# Range restricted old and young lineages show the southern Western Ghats to be both a museum and a cradle of diversity for woody plants

**DOI:** 10.1101/2022.12.11.516896

**Authors:** Abhishek Gopal, D. K. Bharti, Navendu Page, Kyle G. Dexter, Ramanathan Krishnamani, Ajith Kumar, Jahnavi Joshi

## Abstract

The Western Ghats (WG) mountain chain is a global biodiversity hotspot with high diversity and endemicity of woody plants. The latitudinal breadth of the WG offers an opportunity to determine the evolutionary drivers of latitudinal diversity patterns. We examined the spatial patterns of evolutionary diversity using complementary phylogenetic diversity and endemism measures. To examine if different regions of the WG serve as a museum or cradle of evolutionary diversity, we examined the distribution of 470 species based on distribution modelling and occurrence locations across the entire region. In accordance with the expectation, we found that the southern WG is both a museum and cradle of woody plant evolutionary diversity, as a higher proportion of both old and young evolutionary lineages are restricted to the southern WG. The diversity gradient is likely driven by high geo-climatic stability in the south and phylogenetic niche conservatism for moist and aseasonal sites. This is corroborated by persistent lineage nestedness at almost all evolutionary depths (10–135 million years), and a strong correlation of evolutionary diversity with drought seasonality, precipitation and topographic heterogeneity. Our results highlight the global value of the WG, demonstrating, in particular, the importance of protecting the southern WG – an engine of plant diversification and persistence.

## Introduction

Tropical rainforests are one of the most diverse biomes on earth, accounting for nearly half of all known tree diversity [1]. The high diversity of tropical rainforests can be attributed to two important processes operating on evolutionary timescales, deep biome age and climatic stability, allowing for the persistence and accumulation of lineages, and increased speciation rates [2–5]. Tropical rainforests have thus been referred to as both “museums” and “cradles” of diversity [4].

Within the tropics, there is variation in lineage richness over water availability gradients. The high richness in wet sites is driven by strong phylogenetic niche conservatism for wet and warm environments, where angiosperms putatively originated and where angiosperm lineages persist and diversify [6,7]. Furthermore, the favourable environment in wet and aseasonal sites potentially allows for lineages with different affinities, including dry or extratropical, to disperse and establish, resulting in higher diversity [6,8–10]. There are several examples of this “dry-tolerance pattern” [10], wherein dry-adapted lineages have wider ranges and persist in wetter sites, whereas wet-affiliated lineages have a more restricted distribution within the wetter sites [1,6,7], resulting in a nested distribution of lineages over water availability gradients in the tropics [9–11].

The relative influence of different evolutionary and ecological processes in shaping these diversity patterns can be examined using phylogenetic indices of diversity and endemism [12–14]. Areas with a high proportion of old range-restricted lineages have been speculated to be sites with a museum-like diversity pattern [12,15,16]. Alternatively, a high proportion of range-restricted lineages with short branch lengths indicates recently diverged lineages, reflecting sites with a cradle-like diversity pattern [12,15,16]. Documenting spatial variation in phylogenetic diversity indices has also been useful in highlighting sites for conservation [17,18].

Mountain systems, especially in the tropics, are well suited to examine patterns of origin and accumulation of diversity as there is a high level of topographic and climatic heterogeneity over a relatively short geographic distance [19,20]. This provides opportunities for speciation via ecological opportunity and offers sites for refugia (micro-refugia) during climatic instability [20–22]. Consequently, mountains can be an engine for lineage diversification or a refuge for lineage persistence, or both [20,23]. Due to their ancient age, latitudinal breadth, gradients in historical climatic stability and current environment, and topographic heterogeneity, the Western Ghats (WG) mountains represent an ideal system to study evolutionary patterns in the distribution and diversity of old versus young lineages.

The Western Ghats (WG) mountain chain in peninsular India (8°N–21°N) is thought to be one of the oldest yet most dynamic regions of differentiation for the flora and fauna of tropical Asia [24]. It is considered a biodiversity hotspot due to its high diversity and endemism [25]. The diversity and distribution of the extant taxa in the WG are shaped by its complex geo-climatic history, including the breakup of Gondwana, their northward movement across latitudes before merging with Asia (∼50 mya), extensive volcanic activity in the late Cretaceous, and the rise of the Himalayas, monsoon intensification and aridification in the Cenozoic [24,26–28]. Like other mountain chains, the WG have been a refuge during climatic instability, especially the southern Western Ghats, due to their climatic stability and topographic heterogeneity [28,29 and reference therein; 30]. Based on the phytogeography and geographic breaks, the WG are broadly divided into three subdivisions, namely the northern, central, and southern WG [31]. The prominent gaps in the mountain chain are the Goa gap (primarily a climatic barrier) separating the central WG and the northern WG, and the Palghat gap separating the southern WG and the central WG.

The aridification of the Indian peninsula in the Miocene led to the contraction of wet forests to the western slopes of the WG [28,32], which now harbour most of its diversity of evergreen woody plants. The wet forests within this mountain chain are relatively well-defined and form a cohesive biogeographic unit in an otherwise dry peninsula [33]. Over 60% of the woody plant species within these forests are endemic, although endemic clades at the generic level are relatively few (for example, *Agasthiyamalaia, Humboldtia, Otonephelium, Poeciloneuron*) [34]. The interaction of the southwest and northeast monsoons with topography creates a gradient of increasing temperature and precipitation seasonality from the south to the north [11,33,35]. The influence of past geo-climatic events and the current climatic gradient shape the observed taxonomic trend of decreasing woody plant diversity with increasing latitude in the WG [11,36,37].

Here, we use spatial analyses of phylogenetic diversity indices to examine the role of age, stability and phylogenetic niche conservatism in shaping the woody plant diversity gradient in the WG. In particular, we address the following questions.

1. Does the latitudinal gradient in past geo-climatic instability and current seasonality affect the evolutionary diversity of woody plants, resulting in higher persistence and recent divergence of lineages at lower latitudes? The latitudinal gradient in the WG serves as a proxy for historical and current climatic stability and topographic heterogeneity. We predicted that geo-climatic stability and topographic heterogeneity in the southern WG would lead to range-restricted old and young lineages at lower latitudes. In contrast, the northern latitudes are likely to have younger lineages and few range-restricted lineages due to past geo-climatic instability and a present-day climate that is more seasonal.
2. Do woody plant lineages show a nested distribution over water availability and seasonality gradients at multiple evolutionary depths? Water availability is one of the key determinants shaping lineage distribution, resulting in a turnover or nestedness of lineages [7,10]. In the WG there is a strong seasonality gradient, with the northern portion having 6–7 months of dry season, whereas the southern portion has a dry season of only up to 2–3 months, and this gradient has been consistent over geological time (since Eocene-Oligocene) [29 and references therein,33,38]. Taxonomically, there is evidence of a nested distribution due to this seasonality gradient [9,11]; however, it has not been examined if these nested trends persist at deeper evolutionary levels. We examined the distribution pattern of the extant lineages at different evolutionary depths (10 to 135 mya) to examine to which evolutionary depth the observed nested pattern of distribution persists in the WG. In the Cenozoic, the Eocene-Oligocene period is associated with the onset of a seasonality gradient in the subcontinent [29 and references therein], and as the majority of the speciation events occurred during the Miocene, we predict that nestedness will attenuate and potentially be absent at evolutionary depths that are older than the Miocene. Older lineages will have had sufficient time to adapt to the ‘new’ dry and seasonal conditions, while not all younger lineages that have arisen in moist and aseasonal environments since the Miocene would have had time to adapt to dry and seasonal conditions.
3. What are the environmental correlates of evolutionary diversity? In order to understand the potential mechanisms underlying any latitudinal patterns in evolutionary diversity, we examined if the current climate, specifically drought seasonality, mean annual precipitation and topographic heterogeneity, are correlated with evolutionary diversity measures. These predictors have been shown to affect species distributions within the wet gradient in the WG [9,11,36]. As such, we examined if these drivers show a relationship with evolutionary diversity as well. We expected that areas with lower drought seasonality, high annual precipitation and higher topographic heterogeneity, would all show higher evolutionary diversity.
4. Additionally, we examined the contribution of different lineages to the spatial patterns of phylogenetic diversity and if their contribution varies at different evolutionary depths.

To test these predictions, we expand upon the previous phylogenetic studies [29,38], which were restricted to more southern latitudes (south of 16° N), by including the northern limit of the wet gradient in the WG (up to 19° N). The inclusion of higher latitudes is critical in understanding the influence of environmental extremes on evolutionary diversity. Furthermore, the insights from previous phylogenetic studies primarily used metrics of phylogenetic community structure, i.e. how divergent or closely related are species in a given area. While such metrics are insightful in terms of ecological and evolutionary drivers of community assembly, they do not represent well the richness dimension of evolutionary history [14,18,39]. Here, we aim to provide a more comprehensive understanding of the evolutionary diversity patterns and drivers of WG woody plants, using indices sensitive to different aspects of phylogenetic richness.

## 2. Materials and Methods

### 2.1. Dataset

We compiled occurrence datasets from three primary sources [11,40,41], which consist of plots ranging from 0.06 ha to 1 ha in size. Additional occurrence data were obtained from the published literature (Fig. S1). From the compiled dataset, we excluded species with clear dry deciduous and thorny scrub affinities. We focussed on evergreen areas within the Western Ghats, defined as areas where most species are evergreen as defined by their phenology. All species names were validated manually according to The Plant List database (https://www.theplantlist.org/), and synonyms were removed. The final revised list consisted of 470 species with 9,448 occurrence locations at a ∼1 × 1 kmresolution (duplicate occurrences of a species in a grid cell were removed).

### 2.2. Species distribution modelling

To estimate diversity at the scale of the WG and to account for the variable sampling effort across datasets, we employed a presence-only distribution modelling approach using Maxent version 3.4.1 [42]. For each species with >3 occurrence locations, Maxent models were implemented for the full extent of peninsular India (8°–24°N, 68°–91°E) at ∼1 × 1 km resolution, with predictors selected from the WorldClim database of bioclimatic variables [43], specifically mean annual precipitation, mean precipitation of the driest quarter, mean precipitation of the warmest quarter, mean annual temperature, mean temperature of the warmest quarter, mean temperature of the driest quarter, and elevation. These predictors have been shown to affect the species composition of woody plants in the Western Ghats [9,44]. We chose background ‘pseudo-absence’ locations from a bias layer that accounted for variation in sampling effort (see Appendix S2 for more information).

The predictions of habitat suitability for the evergreen woody plants obtained from the best model (Appendix S2) were clipped to the extent of the WG, and the continuous predictions at 1 × 1 km resolution were aggregated to a 10 × 10 km resolution. The species-level maps were stacked to obtain a species richness map of the WG, extending from 8° to 19° with 1756, 10 × 10 kmgrid cells. To get a conservative estimate of diversity for evergreen woody plants, we removed 508 grid cells with less than ten species (minimum observed species richness in the raw plot dataset), which may not represent evergreen sites and thus may be associated with poor model predictability (Fig. S2). The richness maps at 10 × 10 km resolution were used for calculating indices of evolutionary diversity for 348 species. For 12 species with poor model performance and 110 species with ≤ 3 occurrences (largely representing point endemics), only the observed occurrence points were considered for calculating evolutionary diversity and endemism indices. Finally, we clipped the latitudinal extent of the species distribution to the known occurrence locations to get a conservative estimate of species range. The resulting richness maps of clipped versus unclipped data were similar in terms of the richness trends from south to north. However, we decided to focus on the former to get a conservative estimate of species distributions [45]. The code for the SDMs was modified from Bharti et al. [46] (https://github.com/bhartidk/centipede_diversity_endemism). Spatial data were processed using the packages “rgeos” [47], “raster” [48], “sf” [49], “sp” [50], and Maxent models were run using the package “ENMeval” [51] and “dismo” [52] in R 4.1.1 [53]. For more details regarding the species distribution models, refer to Appendix S2.

### 2.3. Phylogenetic tree for WG woody plants

The Smith and Brown, [54] megaphylogeny of seed plants was used to create a phylogeny for the WG woody plants (470 species). Following the recommendations of Qian and Jin, [55], we used the function “phylo.maker”, build.node.1 function and scenario 3 from the R package “V.PhyloMaker” [56] to create the phylogeny, which was then used to calculate the evolutionary diversity indices. In total, 149 species were present in the megaphylogeny and 321 species not initially present were bound to the tree. One genus, *Hydnocarpus* (3 species in the dataset), was added to the tree using “bind.relative” function. This method of using a phylogeny that is well-resolved above species level to calculate evolutionary diversity metrics is robust to unresolved species-level phylogenetic relationships [55,57].

### 2.4. Evolutionarydiversity indices

Evolutionary diversity was quantified using three indices that focus on the richness dimension of evolutionary diversity [14], 1) Faith’s phylogenetic diversity (PD), 2) Time-integrated lineage diversity (TILD), and 3) Phylogenetic endemism (PE). PD is the summation of all branch lengths found in a grid cell [58]. TILD is calculated by integrating the area under a lineage through time plot where the number of lineages is first log transformed [18]. As such, it reduces weighting diversity in recent evolutionary time and shifts it towards deeper evolutionary time. TILD is a useful ‘deep-time’ complement to PD, as PD is strongly driven by young lineage diversity, i.e. the number of species in a sample [6,18]. PE is estimated by weighting each branch length in the phylogeny by the total descendant clade’s range [59]. PE, by integrating the range size information with evolutionary history, is a useful measure to examine how spatially restricted or widely distributed the evolutionary history is in a landscape. As PE is highly sensitive to the area occupied by a species, we excluded species without distribution predictions (122 out of 470 species; see section 2.2). Given that most of the excluded data are point endemics in the southern WG, their inclusion would have only strengthened the observed trends (see Results). Lastly, the total number of species per grid cell was used as a measure of species richness (SR). The indices were calculated using the packages “phyloregion” [60], “ape” [61], and “picante” [62] in R. Data wrangling and tree visualisation were done in R using “tidyverse” [63], and “ggtree [64] packages.

### 2.5. Lineage accumulation and latitudinal trends

The time-calibrated phylogenetic tree was sliced at 10 million year intervals (myr) intervals from the present to 135 million years ago (mya), which is the root age of the WG woody angiosperm phylogeny. The sliced trees at each evolutionary depth were used to examine the accumulation of extant lineages at different evolutionary depths using lineage through time plots (LTT), and the latitudinal distribution of lineages at each evolutionary depth. The method outlined here simply examines the present distribution and diversity of extant lineages at different evolutionary depths [13,65,66]. It does not examine the complete historical diversity or the true range sizes of the ancestral lineages, which would require adequate fossil data. For these analyses, we included both the predicted distribution of species and species for which only occurrence locations were known. The tree was sliced using R code modified from Daru et al. [65], and the lineage age and latitudinal trends were examined using R code modified from Griffiths et al. [13].

### 2.6. Nested distribution of lineage at different evolutionary depths

To examine the nested distribution of lineages at different evolutionary depths, we used the metacommunity approach of Leibold and Mikkelson [67]. The presence of all lineages at each evolutionary depth was ascertained for latitudinal bins and a latitudinally ordered lineage presence-absence matrix was used. Nestedness or turnover was assessed by examining how many times a lineage is replaced at two adjacent sites compared to the average number of replacements when the matrix was randomly re-sorted 1000 times. This was done for each evolutionary depth ranging from 10 to 135 mya. The null matrix was created using the “r1” null model, where the lineage richness of a site is held constant (rows), and the lineage ranges (columns) are filled based on their marginal probability values. Higher observed values than expected indicate turnover, while the converse indicates nestedness [68]. This analysis was done using the function “Turnover” from the “metacom” package in R [69]. Although this analysis is in terms of the lineage diversity, the approach used is ecological in its inference, wherein, nestedness refers to differences in richness among sites, i.e. if lineages in one site are a subset of lineages found in another site and turnover refers to replacement of lineages across sites.We are not using the term nestedness as the phylogenetic literature uses it, that is when one clade is found ‘nested’ within another clade.

### 2.7. Spatial and temporal contribution of key lineages to PD

To identify the key lineages contributing to PD, the branch lengths of each lineage (super order and family level) were summed within 1° latitudinal bins to evaluate their contribution to total PD. Furthermore, at the super order level, in addition to the spatial contribution to PD, we also examined the contributions temporally by summing the branch length of each lineage from a sliced tree from 135 mya to the current time period (see section 2.5). The node identities of the ancestral lineages were identified primarily by its most recent common ancestor (“getMRCA” function in “ape” package in R) and by visually examining the phylogenetic tree. We excluded lineages with only one extant taxon.

### 2.8. Correlates of Phylogenetic Diversity

We examined the environmental correlates of the evolutionary diversity in the WG at 10 × 10 km resolution, using the following predictors, topographic heterogeneity (calculated using WorldClim elevation layer), and mean annual precipitation, from the WorldClim database and mean climatic water deficit (CWD) [70]. Topographic heterogeneity was measured using topographic ruggedness index following Riley et al. [71], wherein the average difference between the focal cells and its neighbouring cells were calculated using the function “terrain” from the raster package. Topographic heterogeneity was calculated at 1 × 1 km resolution and aggregated to 10 × 10 km resolution with higher values indicating high ruggedness. CWD was correlated with temperature and precipitation seasonality in the study area (Fig. S3). CWD reflects the water stress experienced by plants during the dry season and as such is a useful measure to quantify drought. It is measured as the difference between rainfall and evapotranspiration during dry months only (in millimetres per year). It is always a negative number, with higher negative values implying a greater water deficit. These predictors have been shown to affect woody plant diversity patterns in the WG [9,11,36,38] and are also not correlated among themselves (Fig. S3). One grid cell with no predictor information from the WorldClim database was removed.

## 3. Results

### 3.1. Patterns of evolutionary diversity and endemism

All evolutionary diversity indices showed a trend of increasing diversity from northern latitudes to southern latitudes (Fig. 1). Both SR and PD show a similar increase with decreasing latitude (Fig. 1. A, B). The gradient for TILD relative to the other indices was less stark, but with the drier and the seasonal northern sites still having the lowest TILD values (Fig. 1D). On average, the grids in the northern WG had ∼1.4 times lower TILD (Fig. S4D), and ∼3 and ∼6 times lower PD and SR than the grids in the southern WG (Fig. S4. A, B). The sites in the central WG had a comparable spread of PD and SR values to sites in the southern WG, but on average had ∼1.5 times less PD and SR as compared to sites in the southern WG (Fig. S4 A, B). Compared to the other diversity measures, PE shows the starkest contrast in the north-south gradient (Fig. 1C). The average PE value of sites in the southern WG was ∼6 and ∼2 times higher than the sites in the northern WG and central WG, respectively (Fig. S4C). All indices, despite being sensitive to different aspects of the phylogenetic tree, showed very high correlations with each other (Pearson’s correlation > 0.9, *p* < 0.05; Fig. S5). These results highlight that the southern WG are the sites which have a high diversity of old, young and range-restricted lineages.

**Figure 1.**
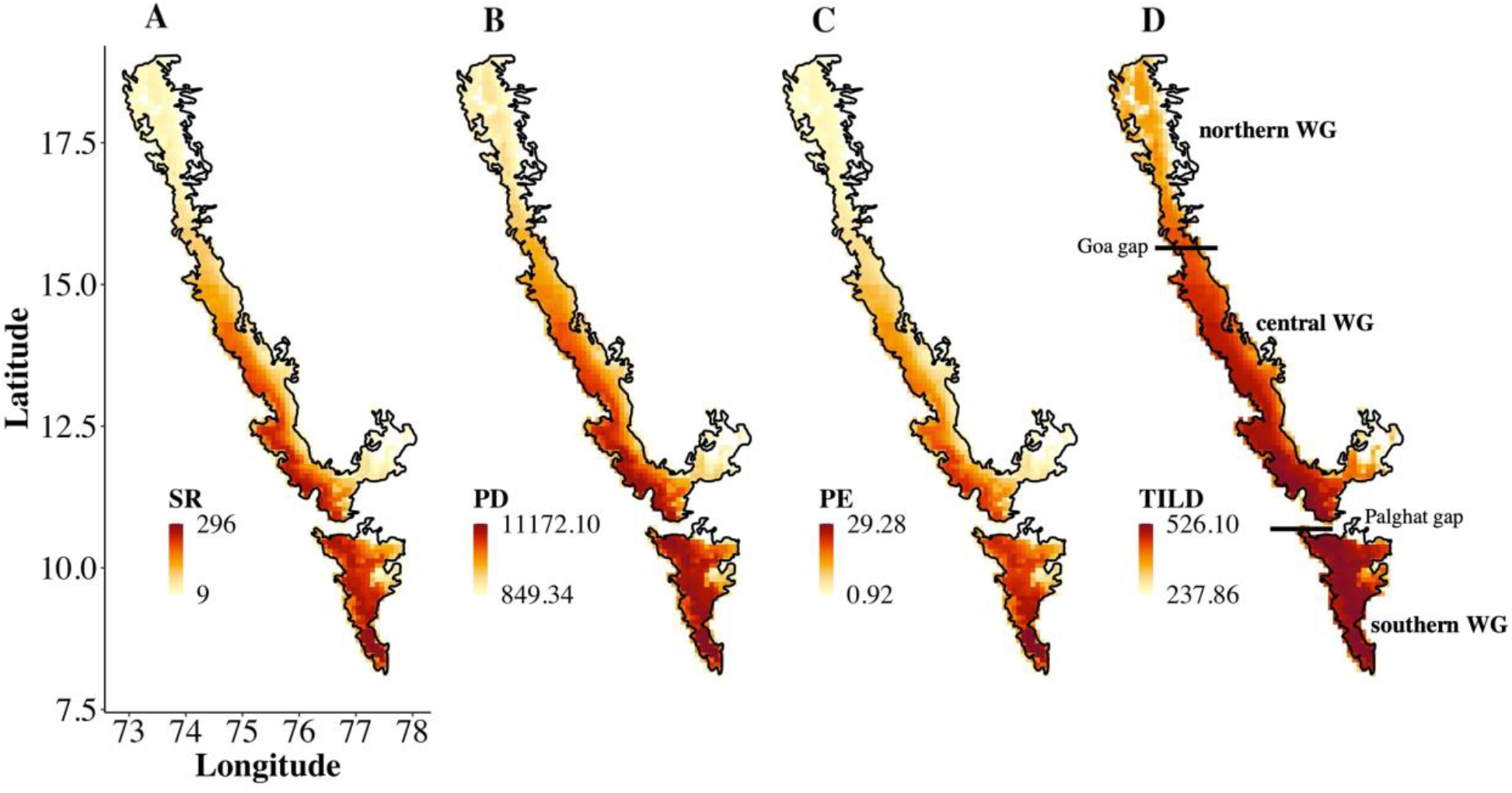
Patterns of the richness of woody plants in the Western Ghats at 10 × 10 km resolution. A. Species richness (SR), B. Phylogenetic diversity (PD), C. Phylogenetic endemism (PE), and D. Time integrated lineage diversity (TILD). The darker colours indicate higher values. PE only included the species for which the SDMs were created (348 species). Previously postulated biogeographic sub-divisions based on geographic and climatic breaks in the mountain chain are shown in panel D.

### 3.2. Spatial and temporal patterns of lineage diversity and latitudinal trends at different evolutionary depths

The lineage through time plot for the extant lineages, shows that the accumulation of lineages is higher at lower latitudes across all evolutionary depths, with almost an order of magnitude higher lineage richness in the current time period (Fig. 2A).

**Figure 2.**
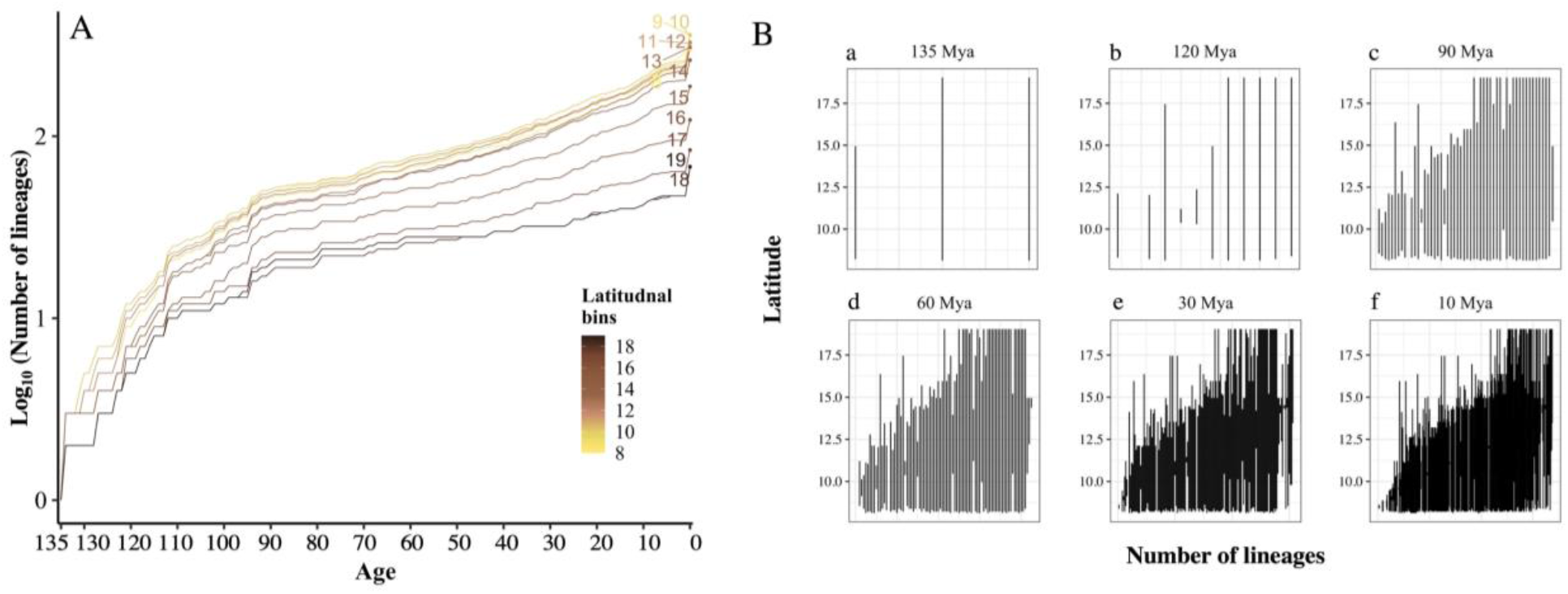
A. Lineage through time plot for the extant taxa in 1° latitudinal bins at 1 million year interval. This shows that the lower latitudes consistently have a higher accumulation of lineages at all evolutionary depths. B. Latitudinal range of lineages at selected evolutionary depths. At the evolutionary depth of period 135 mya (B. a), there are three ancestral lineages representing

In terms of the latitudinal ranges, the lower latitudes show a nested distribution of lineages at all evolutionary depths (Fig. 2B). This is consistent with statistical analysis which shows that lineages consistently display lower turnover than expected along the latitudinal gradient of the WG (Table 1).

**Table 1.** Observed and expected turnover (mean turnover of 1000 randomly sorted matrix) of lineages at each evolutionary depth along the latitudinal gradient. The *p* value tests if the observed lineage turnover along the latitudinal gradient differs from a random expectation. The observed turnover is significantly lower than the expected turnover at all evolutionary depths, indicating a nested lineage distribution at all evolutionary depths along the latitudinal gradient of the Western Ghats.

| Age (mya) | Number of lineages | Observed turnover | Expected turnover | *p* |
| --- | --- | --- | --- | --- |
| 135 | 3 | 0 | 3.91 | < 0.01 |
| 120 | 12 | 0 | 218.41 | < 0.01 |
| 90 | 54 | 95 | 3039.79 | < 0.01 |
| 60 | 85 | 638 | 7952.76 | < 0.01 |
| 30 | 153 | 10750 | 30247.79 | < 0.01 |
| 10 | 337 | 59524 | 145479.55 | < 0.01 |

We further dissected these results to examine the lineage richness patterns above and below 13° latitude, the midpoint of our study region. The lineage richness was consistently higher at the lower latitudes, with the lower latitudes contributing 42% – 34% of the total lineage richness from 120 myr to 10 myr(Fig. 2B; Table 2). There are no lineages restricted to above 13° latitude that are older than 60 myr(Fig. 2B. d; Table 2). There is a striking richness contrast in 30 myr old lineages, with lower latitudes having seven times the lineage richness compared to higher latitudes (Fig. 2B. e; Table 2). The lower latitudes have both older and younger, range-restricted lineages. For example, five lineages that are 120 myr old occur at lower latitudes but none above 13°. Among lineages that are 10 myr old, 114 lineages occur at lower latitudes but only 13 lineages occur at higher latitudes.

**Table 2.** Total lineage richness and lineage restricted to north and south of 13° latitude at selected evolutionary depths. See Table S1 for all evolutionary depths.

| Age (mya) | Total lineages | Num. of lineages $\leq 13^\circ$ | Num. of lineages $> 13^\circ$ |
| --- | --- | --- | --- |
| 135 | 3 | 0 | 0 |
| 120 | 12 | 5 | 0 |
| 90 | 54 | 11 | 0 |
| 60 | 85 | 18 | 1 |
| 30 | 153 | 35 | 5 |
| 10 | 337 | 114 | 13 |

### 3.3. Correlates of evolutionary diversity

Examining the relationship of environmental variables with evolutionary diversity indicates that the CWD is a strong correlate of all diversity indices (Fig. 3; Pearson’s correlation, *r* = 0.74, 0.78, and 0.82 for TILD, PD, and PE respectively with *p* < 0.01). There was a weaker relationship of annual precipitation (Fig. 3; *r* = 0.26, 0.23, 0.13 for TILD, PD, and PE respectively with *p* <0.01) and a moderate correlation of topographic heterogeneity with all diversity indices (Fig. 3; *r* = 0.38, 0.47, 0.50 for TILD, PD, and PE respectively with *p* < 0.01). While CWD is a strong determinant for evolutionary diversity, in terms of annual precipitation, there is a peak in diversity between ∼2000–4000 mm of precipitation. The evolutionary diversity increased with topographic heterogeneity as expected (Fig. 3).

**Figure 3.**
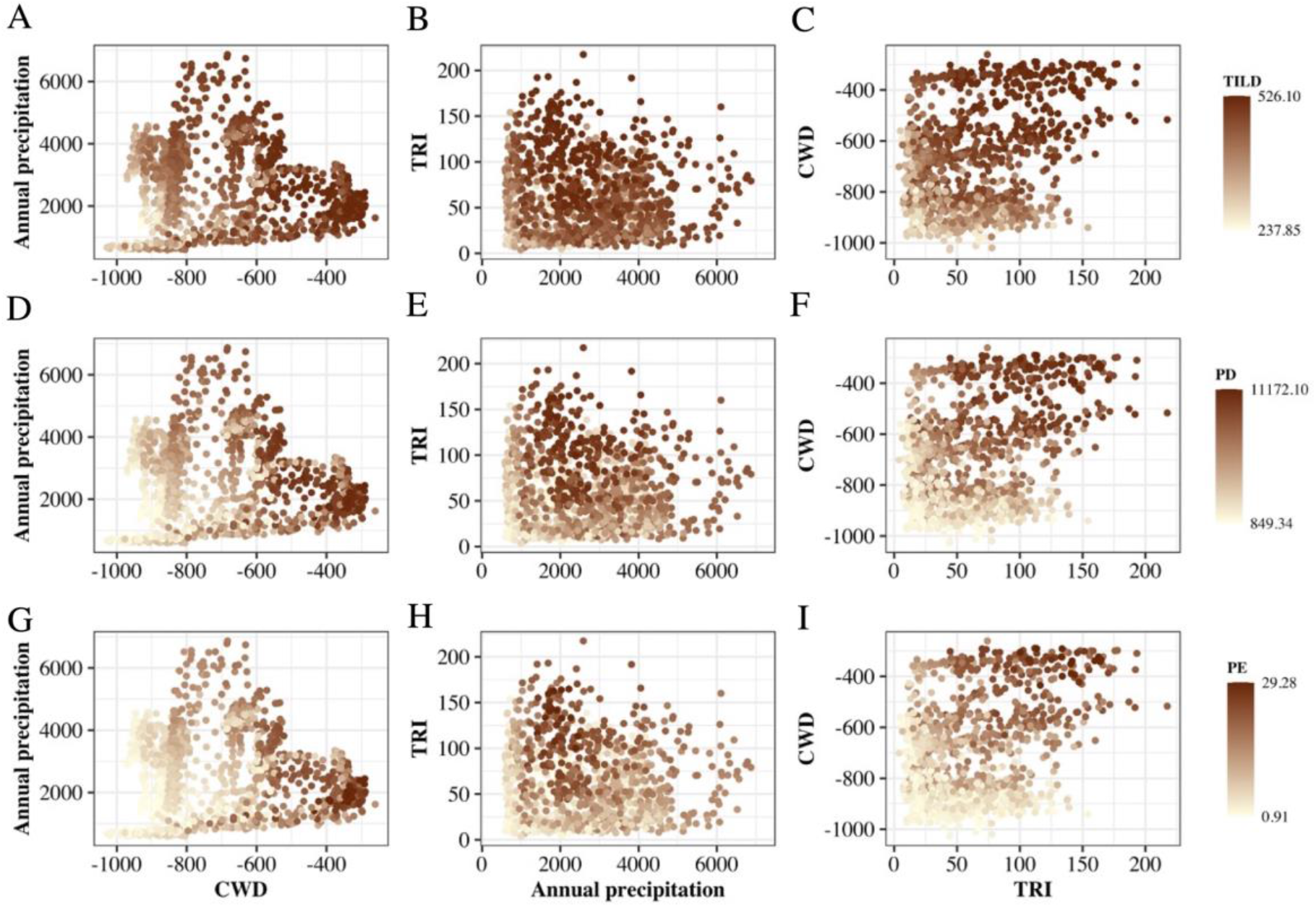
Distribution of different evolutionary indices along the axes of climatic water deficit (CWD), annual precipitation and topographic ruggedness index (TRI). A–C, time-integrated lineage diversity (TILD), D–F phylogenetic diversity (PD), and G–I, phylogenetic endemism (PE). Each point is a measure of the respective evolutionary diversity at a 10 × 10 km grid cell.

### 3.4. Spatial and temporal contribution of key lineages to PD

Our WG woody plant dataset consists of 35 orders, 75 families, and 209 genera. There is a clade-level disparity in the richness of lineages (Table S2) and their contribution to PD (Fig. 4). At the oldest evolutionary depths (120 mya and before), the largest contribution to PD at all latitudes is by basal Eudicots and Magnolids (Fig. 4A). The relative importance of these lineages at younger evolutionary depths is eclipsed by the superorders of Asteridae and Rosidae, which contribute most to PD at younger evolutionary depths across latitudes (Fig. 4A). At the family level, ten speciose families contributed to more than half of the PD (Fig. 4B). Examining the contribution of key families to PD with respect to the latitudinal bins shows that these key families are present throughout the latitudinal extent, except for Fabaceae, which drop off at 15° (Fig. 4B). The relative contribution of the key families to PD varies, however, across the latitudinal regions. For example, in the northern WG, Euphorbiaceae, Lauraceae, Phyllanthaceae, and Rutaceae contribute the most to PD (Fig. 4B). In contrast, the contribution of different families to PD is higher at the lower latitudes, with Annonaceae and Myrtaceae having a relatively higher proportion of PD in both the southern WG and the central WG (Fig. 4B).

**Figure 4.**
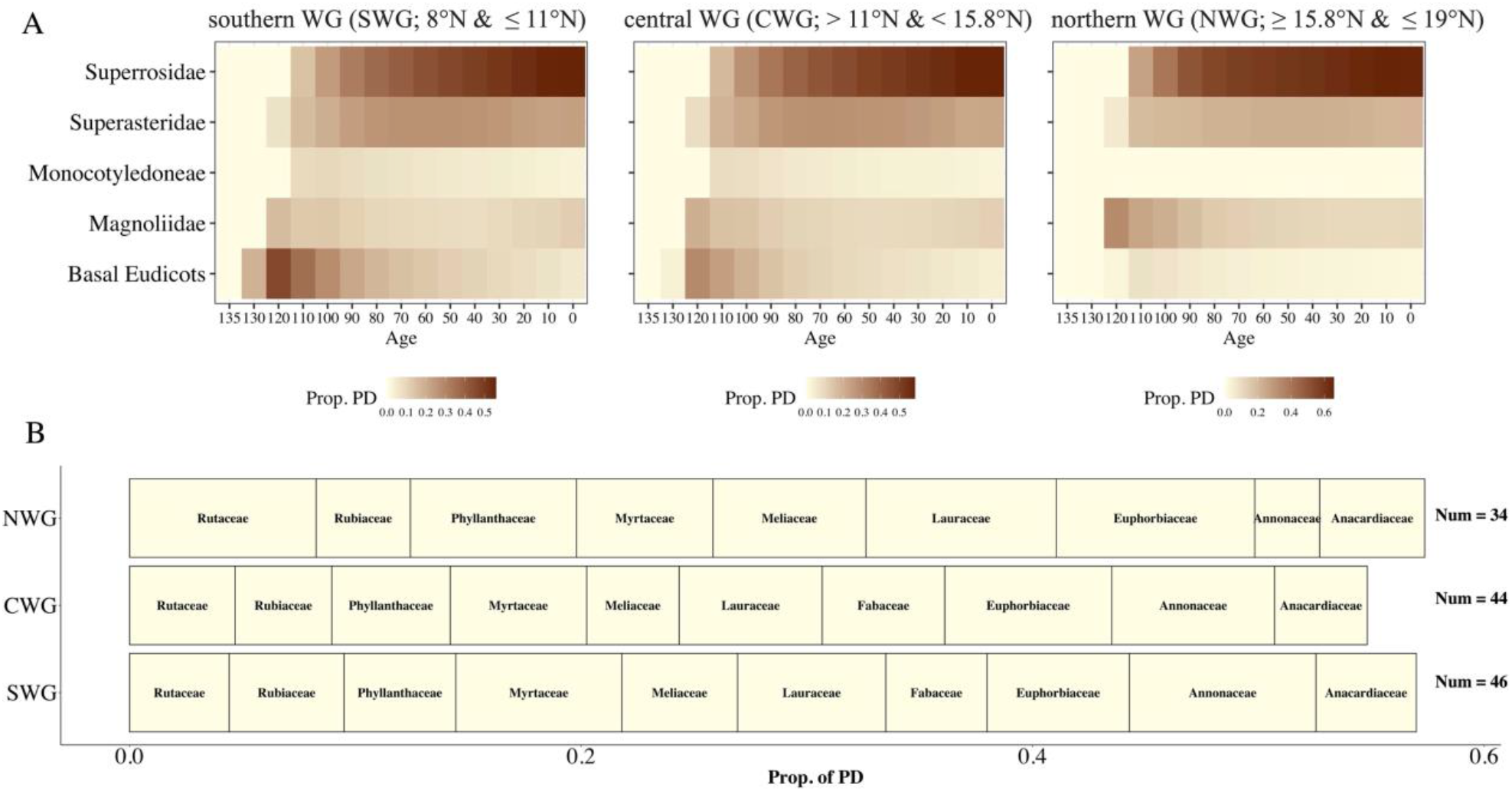
A. Superorder level contribution to phylogenetic diversity (PD) in different biogeographic subdivisions at different evolutionary depths. The colours indicate the proportion of phylogenetic diversity (PD) contributed by each super order at each evolutionary depth and in each biogeographic subdivision. B. Key families and their contribution to PD with respect to each latitudinal bin. The size of each bin reflects the contribution of PD with respect to each biogeographic zone (NWG: northern Western Ghats; CWG: central Western Ghats; SWG: southern Western Ghats). Number on the right-hand side of figure B refers to the total number of families found in each biogeographic subdivision.

## 4. Discussion

We find that the southern WG are a hotbed of evolutionary diversity, facilitating both the persistence and recent divergence of lineages. This latitudinal asymmetry in lineage diversity is likely to be driven by phylogenetic niche conservatism for moist and aseasonal sites, which in congruence with higher geo-climatic stability and higher topographic heterogeneity in the lower latitudes, limits extinction and allows for higher diversification rates. Across all evolutionary depths, some lineages do manage to occur in the more seasonal and dry northern WG, but other lineages are restricted to areas of the WG with low seasonality and higher water availability, resulting in the nested distribution pattern with respect to latitude that persists at all evolutionary depths. Current environmental variables in terms of drought seasonality, precipitation and topographic heterogeneity shape the evolutionary diversity. These results are in line with studies in the Americas that show that within the tropics, water availability is a stronger driver of the evolutionary diversity of woody plants than temperature [7,72].

### 4.1. Southern WG as a museum and a cradle of diversity

Evolutionary diversity measures show a decreasing trend of diversity from lower to higher latitudes for woody plants in the WG. TILD, compared to the other indices, drops off only in the most aseasonal and dry sites of the northern regions of the WG, indicating that over the evolutionary history of the WG, older lineages have been able to colonise most of the WG. Both PD and SR show almost identical patterns of reduced diversity in the northern latitudes, with PE showing the starkest reduction. Given that all indices are strongly correlated with each other, it is clear that the lower latitudes harbour both range-restricted old and young lineages. Our results build on previous studies in the WG [29,38] by emphasising that the high evolutionary diversity at lower latitudes is shaped by the significant contribution of range-restricted lineages.

Spatially restricted evolutionary history, with long or short branch lengths, indicates older or recent in-situ divergence, respectively [59,73,74]. Areas with greater topographic complexity and environmental heterogeneity contribute to ancient and recent divergence by acting as refugia for the persistence of lineages and providing the ecological opportunity for speciation [15,20–22,75]. Historic climatic stability may also aid in the persistence of older lineages across paleoclimatic oscillations, resulting in higher PE [20,73,76,77], due to fewer extinctions and greater time for species accumulation. Our results are in line with global and regional studies, wherein mountains, due to a gradient in historical and current climatic stability coupled with environmental heterogeneity, harbour high phylogenetic endemism and exhibit both persistence and recent divergence of lineages [12,15,77,78].

### 4.2. Phylogenetic niche conservatism for wet, aseasonal sites is likely to shape the latitudinal asymmetry in evolutionary diversity

Niche conservatism for wet and aseasonal sites [6,7] is likely to be one of the key mechanisms underlying the observed asymmetry in evolutionary diversity from lower to higher latitudes. While some older lineages are present throughout the WG, 42%–21% of these lineages, which are 120 myr old and 60 myr old respectively, are found only below 13°N (Table 2). At an evolutionary depth of 30 myr, there is seven times higher lineage richness at lower latitudes compared to higher latitudes. Of the lineages which are 10 myr old, 34% (out of 337 lineages) are found only below 13° (Table 2). Overall, these results suggest deep niche conservatism for wet and aseasonal lower latitudes, even within the relatively wet evergreen portion of the WG, which are the focus of our study. For example, key clades such as Magnolids, Rosids and Asterids, while present throughout the WG, are the highest contributors to PD in the lower latitudes (Fig. 4A). These results are statistically confirmed by the nestedness analysis of lineages at different evolutionary depths. It is not surprising that we do not find a pattern of turnover at any evolutionary depth, as previous studies at the shallowest evolutionary depth (i.e. species) find a pattern of nestedness in the WG [9,11]. In fact, it would be impossible to have turnover at deeper evolutionary depths if there is nestedness at the lowest evolutionary depths. However, it is surprising that the nestedness persists at all evolutionary depths in the WG up to the age of angiosperms themselves (135 mya). Our results are consistent with those of a previous study that showed that the precipitation niches of lineages are conserved between the WG and Central America [79].

### 4.3. Seasonal drought, annual precipitation, and topographic heterogeneity shape evolutionary diversity

Water availability, quantified here using values of mean annual precipitation and CWD, is the key driver of evolutionary diversity in the tropics [7,10,72]. Our results indicate that water availability is likely to limit lineage distribution even within the relatively wetter portions of the WG that we study here, especially for lineages with putative origins in the moist end of the gradient [6,7,72,79]. This is corroborated by recent studies, which show that precipitation and tolerance to drought limit the distribution of evergreen tree species in WG [9,11]. This results in a nested distribution, wherein species with wider drought tolerances can persist throughout the WG, whereas species with narrow climatic tolerance are restricted to the lower latitudes due to the ecophysiological barrier at higher latitudes.

The importance of topographic heterogeneity across the WG mountains, is in accordance with the expectation that more heterogeneous sites have higher evolutionary diversity due to increased ecological opportunity resulting in higher speciation [20–22]. There is some evidence for this in the WG, in which species’ elevation ranges decrease from higher to lower latitudes, suggesting higher niche packing in the lower latitudes [11]. While these predictors are likely to be important for evolutionary diversity indices, it is difficult, however, to truly tease apart their influence as they all covary together with the latitudinal gradient and affect species distribution [9,11].

To summarise, the high taxonomic endemism in the lower latitudes of the WG [11,37] is indicative of the role of in-situ divergences in shaping evolutionary diversity trends. Additionally, the higher evolutionary diversity in the lower latitudes is also due to the persistence of old lineages and the presence of lineages with both narrow and wide climatic tolerance (Fig. 3). In contrast, the northern communities have the subset of lineages that can persist at climatic extremes, which exist there either via range expansion from the lower latitudes or dispersal events.

### 4.4. Insights from biogeographic history of WG in shaping the extant evolutionary diversity

Biogeographic processes, such as vicariance and dispersal, coupled with geological events, are the primary determinants shaping regional diversity and diversification dynamics [80]. The WG woody plant community is thought to be shaped by ancient Gondwanan vicariance, dispersal into India from Southeast Asia and in-situ speciation [27 and references therein]. It has been argued that many ancient lineages with Gondwanan affinity went extinct in the Cretaceous due to volcanic activity, creating empty niches which were occupied by Southeast Asian lineages with wet forest affinity through multiple dispersal events after the Indian plate collided with Asia [27,28,81,82]. Within the WG, the southern WG, in addition to aiding the persistence of lineages, is likely to be characterised by older and recent dispersal events. The environmentally diverse yet generally favourable conditions there likely eased successful establishment, followed by in-situ divergence. The northern WG, in contrast, is likely to be shaped by extinction and range-expansion events from the southern WG and potentially elsewhere, but with less frequent successful establishment.

There has been criticism of the use of the terms “museum” and “cradle”, as the dichotomous definition is likely to be too simplistic to decipher complex patterns occurring over evolutionary time scales [5,83]. In the case of the WG, our results are unequivocal that the lower latitudes are likely to be both a museum and cradle of diversity, as indicated by old and young range-restricted lineages. The northern WG, however, may ultimately be found to exhibit a more complex pattern, given its historic climatic instability and high current climatic seasonality. A more detailed clade-level examination might shed light on which of the processes influence the observed trends and if the lineages found here are characterised by different ages and variable diversification rates or constant diversification rates [1].

### 4.5. Conservation implications

Our study is a comprehensive examination of the evolutionary diversity of the woody plants of the WG, highlighting the potential mechanisms driving the observed diversity gradient and the conservation value of this region. PD, while recognised as an important measure to conserve biodiversity [17,58], is less applied in relation to taxonomic-based conservation measures. Previous efforts to map species-level endemics [84] can now be extended to include PE and PD at the scale of WG. Based on our results here, the identification of priority sites based on species richness and evolutionary diversity are likely to overlap; nonetheless, the inclusion of evolutionary history can further augment and strengthen the support for existing protected areas.

All measures of evolutionary diversity point towards exceptional diversity at lower latitudes. The northern WG, representing the extreme limit of where wet forest can occur, have unsurprisingly low diversity. Nonetheless, the northern WG represents ∼5000 myr of evolutionary history in terms of woody plants alone (calculated as summation of branch length of lineages found in this region, in units of millions of years). While there are very few endemic woody species in the northern WG, it harbours relatively higher endemism of other plant groups, such as shrubs and herbaceous species, including endemic radiations [85,86]. As our study focuses only on the standing woody plants (generally > 10 cm diameter at breast height) and their evolutionary diversity, future studies that include other groups such as pteridophytes, gymnosperms, and other plant habits (herbs, lianas, climbers etc.) to the dataset are needed to obtain a more holistic view of the spatial distribution of the evolutionary diversity of plants in the WG. Shrubs, in particular, show much higher endemism, and the WG are likely to hold one of the richest understory shrub communities globally [87]. However, obtaining reliable occurrence locations and phylogenetic data for these species is likely to be a critical bottleneck.

### 4.6. Going beyond occurrence data, the use of presence-based species distribution modelling and their limitations

Estimating species ranges is a non-trivial endeavour [88]. While distribution models like Maxent are likely to overpredict, occurrence information from plot-based data or herbaria is likely to underpredict distributions [45]. The true range of species is likely to lie between these two extremes. We acknowledge the likely overpredictions due to species distribution models, based on presence-only data, as a potential source of bias in this study. However, using model transferability and performance as a criterion to select the models [89], having a species-specific model fitting and tuning, limiting the latitudinal species extent to known occurrence locations, and removing grid cells having species less than the minimum number of species in a plot, the richness estimates reported here are at best a conservative estimate of the evolutionary diversity indices. Additionally, rare species, where occurrence locations were not modelled (those with only 1–3 occurrences), predominantly occur in the southern WG and have narrow distributions. The inclusion of the ‘true’ ranges of these species is unlikely to change the outcome of our study but would rather further strengthen the observed patterns of higher richness in the southern WG.

## 5. Conclusion

In summary, examining the spatial dimension of evolutionary diversity of evergreen woody plants indicates that the lower latitudes of the WG harbour substantial diversity of old and young range-restricted lineages. These results support previous studies on mountains playing a dual role of persistence and divergence, mediated by region-specific biogeographic processes. Examining the lineage-age distribution trends and the correlates of phylogenetic diversity indicates the role of niche conservatism for wet and aseasonal sites in shaping the distribution of older and younger lineages. These results further give support to the idea that precipitation is a key driver of the evolutionary diversity of woody plants in the tropics, both globally and regionally as well. Our results highlight the exceptional evolutionary diversity of the WG mountains and the role of southern WG in the persistence and generation of woody plant diversity.

## Data accessibility

The raw data used for the study were taken from three published studies and were supplemented with occurrence locations from secondary data. For the dataset from Page and Shanker, [11], both plot data and secondary locations, are available as a supplementary file in the original publication. The dataset from Ramesh et al. [41] is accessible as a data paper in the original publication. The dataset provided by Krishnamani et al. [40] is available on request to the respective authors. The scripts for running the SDMs were modified using the code from Bharti et al. [46]: https://github.com/bhartidk/centipede_diversity_endemism. The supporting figures and tables, and the processed data are available in Appendix S1, S2, and S3 respectively. The code used for the analysis is accessible in the GitHub Repository at https://github.com/abhitims/Evolutionary_diversity_of_WG_woody_plants.

## Authors’ contribution

AG and JJ conceptualised the study with inputs from KGD, NP and DKB. NP, RK, and AJ contributed to the data. Data compilation and curation was done by AG with inputs and support from NP, DKB, and JJ. Analysis was done by AG, with inputs from JJ, KGD, DKB and NP. AG, JJ, KGD, DKB, and NP, interpreted the results. AG wrote the first draft of the manuscript with inputs from all the authors. JJ secured the funding for the study.

## Acknowledgements

AG acknowledges the support of the Student Conference on Conservation Science, Cambridge, UK and the Miriam Rothschild Travel Bursary Programme for facilitating and supporting a one-month internship at the University of Edinburgh. AG thanks the University of Edinburgh and the administrative team at Crew Building for facilitating a smooth stay during the internship period. AG thanks researchers at PLEEBs (PLant Evolutionary Ecologists and Biogeographers), Bruno Garcia Luize, researchers of the Evol-Eco lab, Divya B., RishiddhJhaveri, and Hannah Krupa for discussions and inputs. AG thanks Pragyadeep Roy, S. Manu, Onkar Kulkarni, Vinay Teja, and Souparna Chakrabarty for their help in setting up the analysis on the server. AG thanks Dr. Rohit Naniwadekar for his input and feedback on the manuscript and analyses. AG thanks Hannah Krupa for proofreading the manuscript. We thank Ramesh et al. for keeping their dataset open.

