## Supplementary Table 1 for "Range restricted old and young lineages show the southern Western Ghats to be both a museum and a cradle of diversity for woody plants"

**Appendix S1**

Range restricted old and young lineages show the southern Western Ghats to be a both museum and a cradle of diversity for woody plants

Author list: Abhishek Gopal[1,2*], D. K. Bharti[1✢], Navendu Page[3✢], Kyle G. Dexter[4,5], Ramanathan Krishnamani[6], Ajith Kumar[7], Jahnavi Joshi[1,2*]

Affiliations:
[1] CSIR-Centre for Cellular and Molecular Biology, Uppal Road, Hyderabad
[2] Academy of Scientific and Innovative Research (AcSIR), Ghaziabad, India
[3] Wildlife Institute of India, Dehradun, India
[4] School of GeoSciences, University of Edinburgh, Edinburgh, UK
[5] Tropical Diversity Section, Royal Botanic Garden Edinburgh, Edinburgh, UK
[6] The Rainforest Initiative, Coimbatore, Tamil Nadu, India
[7] Centre for Wildlife Studies, Bangalore, Karnataka, India

[✢]D. K. Bharti and Navendu Page contributed equally to the study.

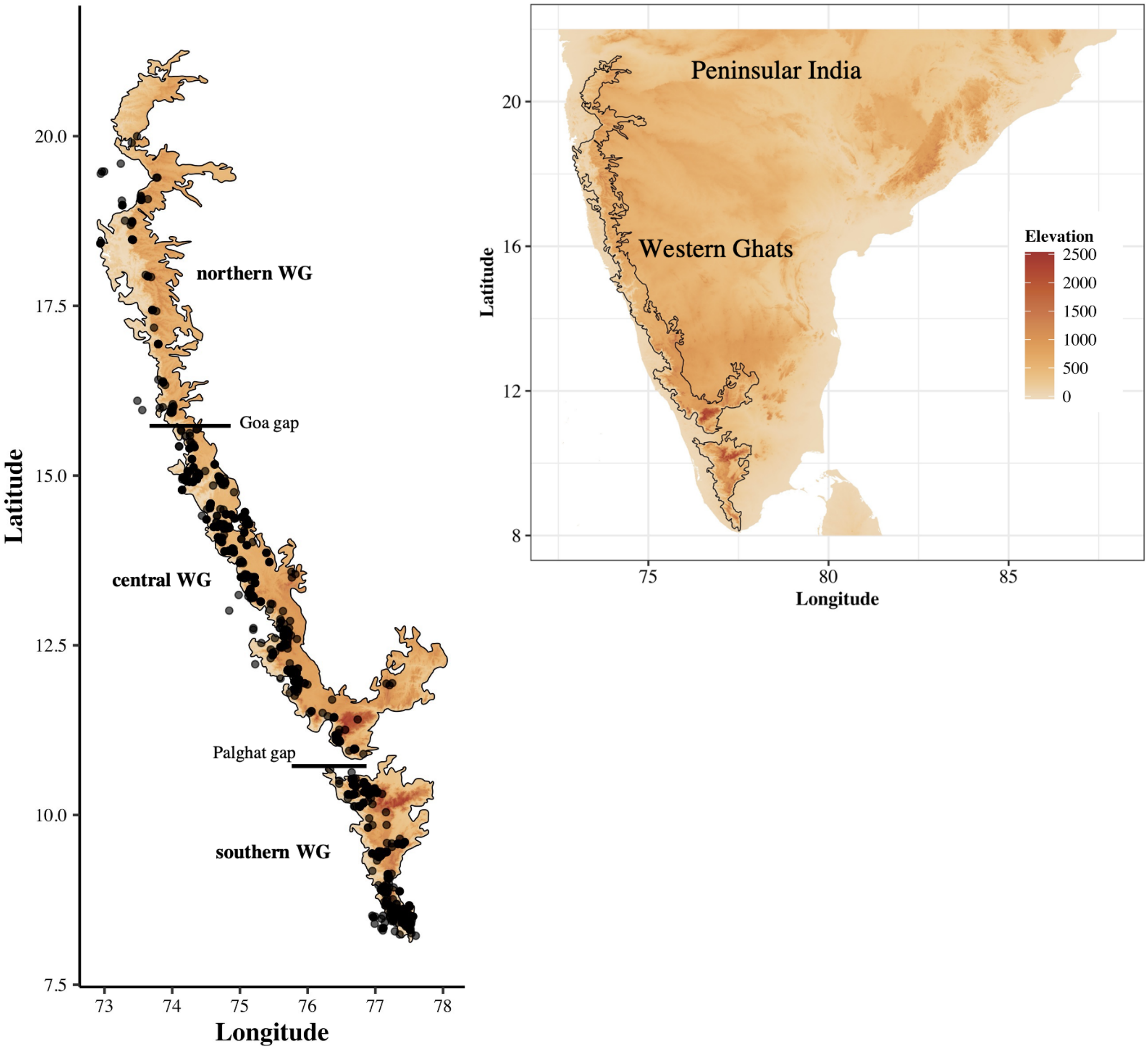


Fig S1. Study site and the elevation profile of peninsular India and the WG. Points indicate the unique occurrence location of species in a 1 x 1 km grid cell. The occurrence locations were compiled from Page and Shanker, 2020, Ramesh et al., 2010, and Krishnamani et al., 2004.

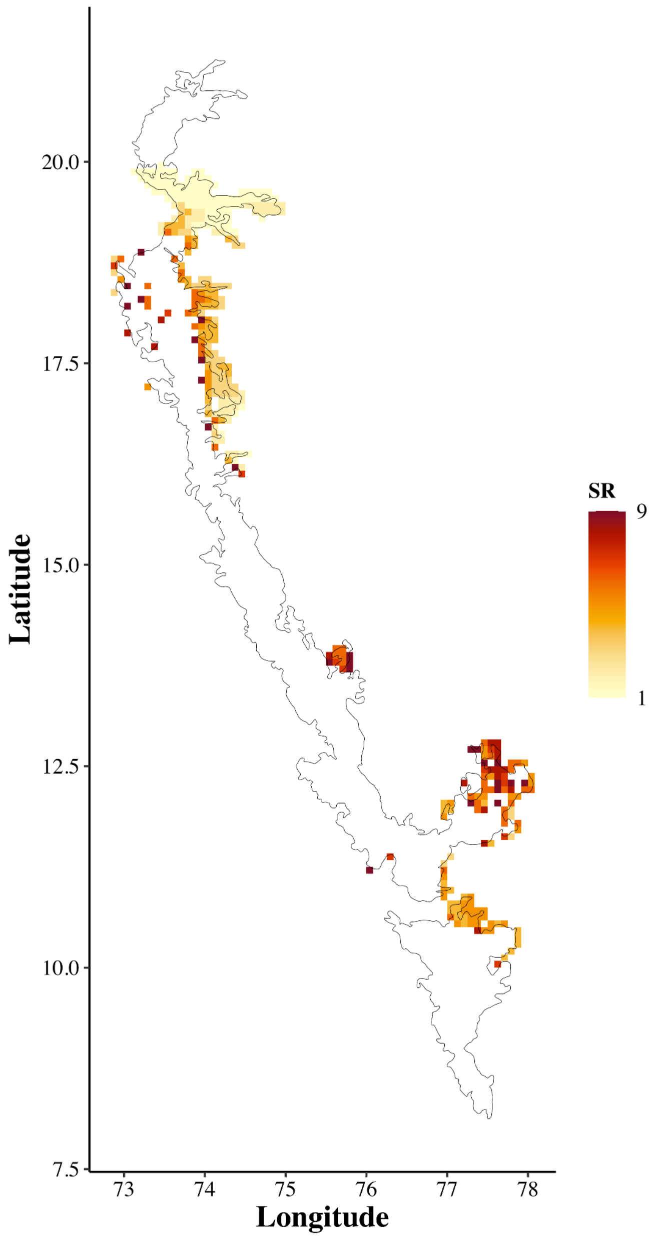


Fig S2. Grids which have less than 10 species per 10 x 10 km grid cell. These grids were removed to get a more conservative estimate of the richness measure of evergreen species in the Western Ghats.

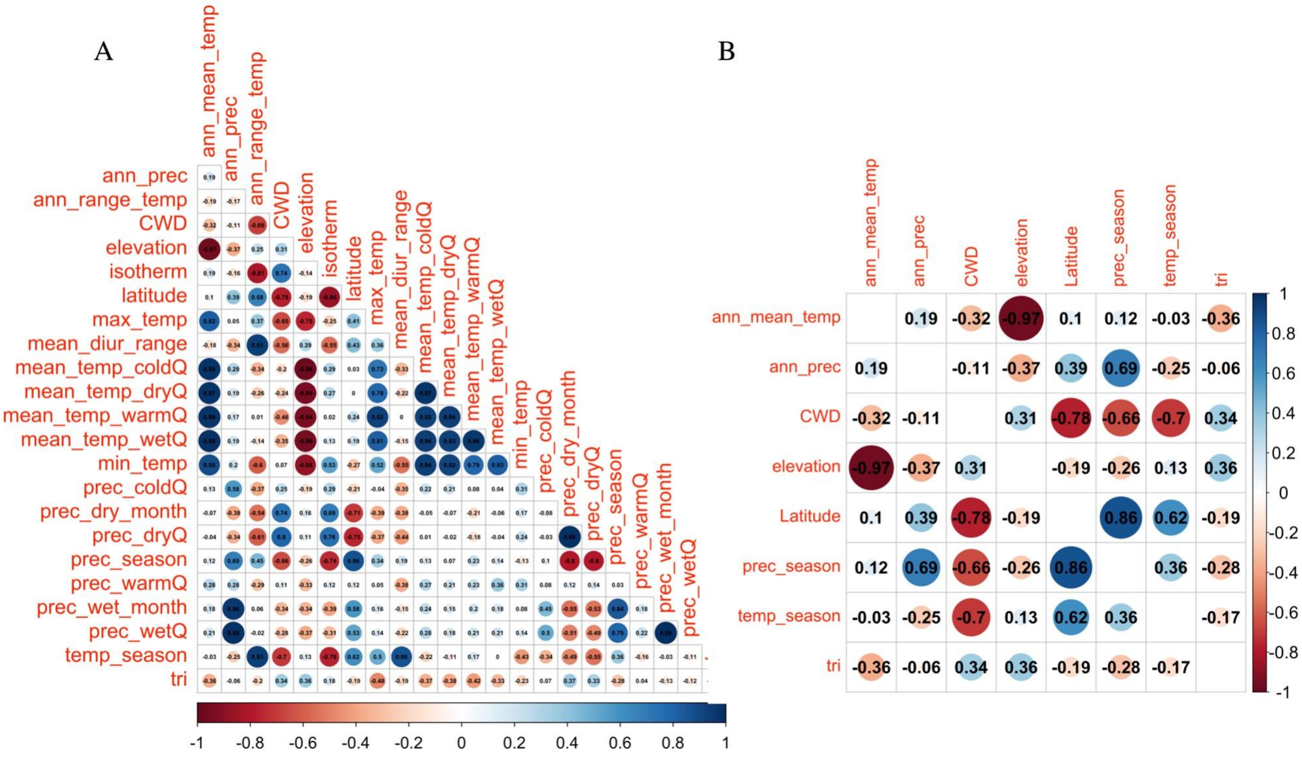


Fig S3. Pairwise correlation between all the (A) WorldClim variables and (B) select WorldClim variables used for analysis. Variable code: **ann_mean_temp:** annual mean temperature, **ann_prec**: annual precipitation**, ann_range_temp**: annual range of temperature**, isotherm**: isothermality,, **max_temp**: maximum temperature of the warmest month, **mean_diur_range**: mean diurnal range of temperature,**mean_temp_coldQ**: mean temperature of the coldest quarter, **mean_temp_warmQ**: mean temperature of the warmest quarter, **mean_temp_dryQ**: mean temperature of the driest quarter, **mean_temp_wetQ**: mean temperature of the wettest quarter, **min_temp**: minimum temperature of the coldest month, **prec_coldQ**: precipitation of the coldest quarter, **prec_dry_month**: precipitation of the driest month, **prec_dryQ**: precipitation of the driest quarter, **prec_season**: precipitation seasonality, **prec_warmQ**: precipitation of the warmest quarter, **prec_wet_month**: precipitation of the wettest month, **prec_wetQ**: precipitation of the wettest quarter, **temp_season**: temperature seasonality, **CWD**: mean climatic water deficit, **tri**: Topographic ruggedness index.

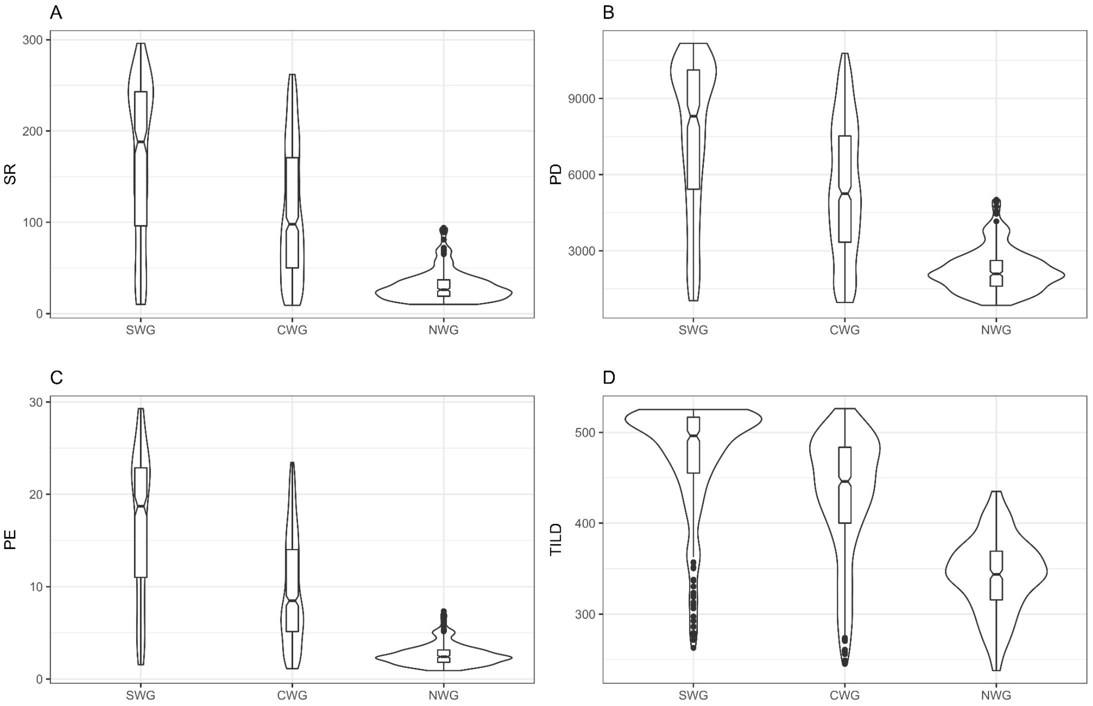


Fig S4. Patterns of the median richness of woody plants in the three biogeographic subdivisions in the Western Ghats (WG) at a 10 x 10 km resolution. A) Species richness (SR), B) Phylogenetic diversity (PD), C) Phylogenetic endemism (PE), and D) Time integrated lineage diversity (TILD), per 10 x 10 km grid cell. Shown is a wedge plot embedded within a violin plot. SWG: southern WG ≤ 11°N, CWG: central WG, > 11°N and < 15.8°N, NWG: northern WG: ≥ 15.8°N.

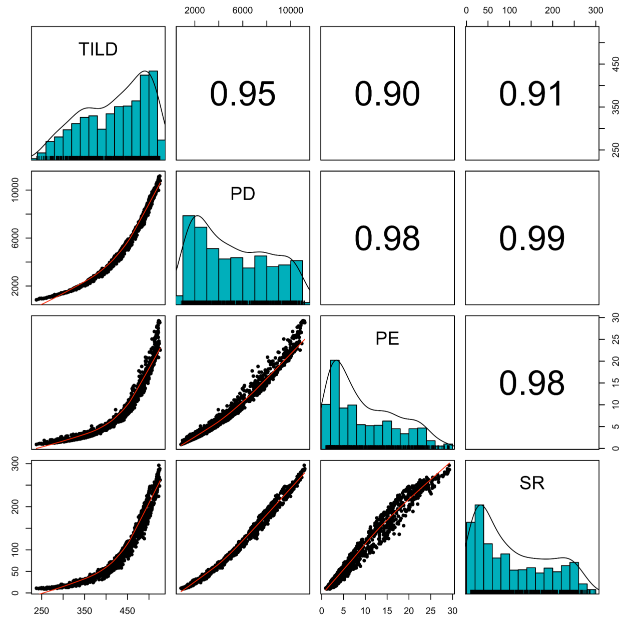


Fig S5. Pearson’s correlation between the different phylogenetic indices

Table S1. Lineage richness and latitudinal trends at all evolutionary depths

| Age (mya) | Total lineages | Num. of lineages ≤ 13° | Num. of lineages > 13° |
|---|---|---|---|
| 135 | 3 | 0 | 0 |
| 130 | 5 | 2 | 0 |
| 120 | 12 | 5 | 0 |
| 110 | 25 | 7 | 0 |
| 100 | 34 | 7 | 0 |
| 90 | 54 | 11 | 0 |
| 80 | 60 | 13 | 0 |
| 70 | 70 | 15 | 0 |
| 60 | 85 | 18 | 1 |
| 50 | 100 | 21 | 2 |
| 40 | 120 | 24 | 3 |
| 30 | 153 | 35 | 5 |
| 20 | 228 | 68 | 8 |
| 10 | 337 | 114 | 13 |

Table S2. Family-level summary of evergreen woody plants in the dataset with the number and proportion of endemics per family. The endemicity was assigned following Ramesh et al., 1997 , Davidar et al., 2018, Page and Shanker, 2020, Kew plants of the world (https://powo.science.kew.org), online flora-BIOTIK (http://flora-peninsula-indica.ces.iisc.ac.in/species.php?id=A&cat=all), and based on inputs by NP.

| Sno. | Order | Family | Number of species | Number of endemics | Proportion of endemics |
|---|---|---|---|---|---|
| 1 | Apiales | Araliaceae | 4 | 3 | 0.75 |
| 2 | Apiales | Pittosporaceae | 2 | 2 | 1 |
| 3 | Aquifoliales | Stemonuraceae | 2 | 1 | 0.5 |
| 4 | Arecales | Arecaceae | 4 | 2 | 0.5 |
| 5 | Asparagales | Asparagaceae | 1 | 0 | 0 |
| 6 | Asterales | Asteraceae | 2 | 1 | 0.5 |
| 7 | Boraginales | Boraginaceae | 1 | 0 | 0 |
| 8 | Brassicales | Capparaceae | 1 | 1 | 1 |
| 9 | Buxales | Buxaceae | 1 | 0 | 0 |
| 10 | Celastrales | Celastraceae | 9 | 7 | 0.78 |
| 11 | Cornales | Nyssaceae | 1 | 1 | 1 |
| 12 | Crossosomatales | Staphyleaceae | 2 | 1 | 0.5 |
| 13 | Cucurbitales | Tetramelaceae | 1 | 0 | 0 |
| 14 | Dilleniales | Dilleniaceae | 1 | 1 | 1 |
| 15 | Dipsacales | Adoxaceae | 1 | 0 | 0 |
| 16 | Ericales | Ebenaceae | 21 | 12 | 0.57 |
| 17 | Ericales | Pentaphylacaceae | 1 | 0 | 0 |
| 18 | Ericales | Primulaceae | 7 | 4 | 0.57 |
| 19 | Ericales | Sapotaceae | 10 | 5 | 0.5 |
| 20 | Ericales | Symplocaceae | 7 | 5 | 0.71 |
| 21 | Ericales | Theaceae | 1 | 1 | 1 |

| | | | | | |
|---|---|---|---|---|---|
| 22 | Fabales | Fabaceae | 10 | 7 | 0.7 |
| 23 | Fabales | Polygalaceae | 1 | 1 | 1 |
| 24 | Gentianales | Apocynaceae | 4 | 2 | 0.5 |
| 25 | Gentianales | Gentianaceae | 1 | 0 | 0 |
| 26 | Gentianales | Rubiaceae | 43 | 34 | 0.79 |
| 27 | Icacinales | Icacinaceae | 1 | 0 | 0 |
| 28 | Lamiales | Bignoniaceae | 2 | 1 | 0.5 |
| 29 | Lamiales | Lamiaceae | 3 | 0 | 0 |
| 30 | Lamiales | Oleaceae | 7 | 5 | 0.71 |
| 31 | Laurales | Lauraceae | 43 | 35 | 0.81 |
| 32 | Magnoliales | Annonaceae | 26 | 23 | 0.88 |
| 33 | Magnoliales | Magnoliaceae | 2 | 1 | 0.5 |
| 34 | Magnoliales | Myristicaceae | 4 | 4 | 1 |
| 35 | Malpighiales | Calophyllaceae | 6 | 5 | 0.83 |
| 36 | Malpighiales | Centroplacaceae | 1 | 0 | 0 |
| 37 | Malpighiales | Chrysobalanaceae | 2 | 2 | 1 |
| 38 | Malpighiales | Clusiaceae | 7 | 5 | 0.71 |
| 39 | Malpighiales | Dichapetalaceae | 1 | 0 | 0 |
| 40 | Malpighiales | Erythroxylaceae | 2 | 2 | 1 |
| 41 | Malpighiales | Euphorbiaceae | 27 | 15 | 0.56 |
| 42 | Malpighiales | Flacourtiaceae | 3 | 3 | 1 |
| 43 | Malpighiales | Ochnaceae | 1 | 0 | 0 |
| 44 | Malpighiales | Phyllanthaceae | 16 | 8 | 0.5 |
| 45 | Malpighiales | Putranjivaceae | 6 | 6 | 1 |
| 46 | Malpighiales | Rhizophoraceae | 2 | 1 | 0.5 |
| 47 | Malpighiales | Salicaceae | 6 | 3 | 0.5 |
| 48 | Malpighiales | Violaceae | 1 | 0 | 0 |
| 49 | Malvales | Dipterocarpaceae | 10 | 10 | 1 |
| 50 | Malvales | Malvaceae | 9 | 4 | 0.44 |

| | | | | | |
|---|---|---|---|---|---|
| 51 | Malvales | Thymelaeaceae | 1 | 0 | 0 |
| 52 | Metteniusales | Metteniusaceae | 1 | 0 | 0 |
| 53 | Myrtales | Combretaceae | 4 | 2 | 0.5 |
| 54 | Myrtales | Lythraceae | 1 | 1 | 1 |
| 55 | Myrtales | Melastomataceae | 13 | 12 | 0.92 |
| 56 | Myrtales | Myrtaceae | 29 | 25 | 0.86 |
| 57 | Oxalidales | Elaeocarpaceae | 5 | 4 | 0.8 |
| 58 | Pandanales | Pandanaceae | 1 | 0 | 0 |
| 59 | Proteales | Proteaceae | 1 | 0 | 0 |
| 60 | Ranunculales | Menispermaceae | 1 | 0 | 0 |
| 61 | Rosales | Cannabaceae | 3 | 0 | 0 |
| 62 | Rosales | Moraceae | 13 | 2 | 0.15 |
| 63 | Rosales | Rosaceae | 1 | 0 | 0 |
| 64 | Rosales | Urticaceae | 4 | 0 | 0 |
| 65 | Sabiales | Sabiaceae | 2 | 0 | 0 |
| 66 | Santalales | Olacaceae | 2 | 2 | 1 |
| 67 | Santalales | Santalaceae | 1 | 0 | 0 |
| 68 | Sapindales | Anacardiaceae | 16 | 13 | 0.81 |
| 69 | Sapindales | Burseraceae | 1 | 0 | 0 |
| 70 | Sapindales | Meliaceae | 19 | 6 | 0.32 |
| 71 | Sapindales | Rutaceae | 13 | 5 | 0.38 |
| 72 | Sapindales | Sapindaceae | 8 | 1 | 0.12 |
| 73 | Sapindales | Simaroubaceae | 1 | 0 | 0 |
| 74 | Saxifragales | Daphniphyllaceae | 1 | 1 | 1 |
| 75 | Vitales | Vitaceae | 1 | 0 | 0 |

Table S3. Species-level summary and endemism status of evergreen woody plants in the dataset. The endemicity was assigned following Ramesh et al., 1997 , Davidar et al., 2018, Page and Shanker, 2020, Kew plants of the world (https://powo.science.kew.org), online flora (http://flora-peninsula-indica.ces.iisc.ac.in/species.php?id=A&cat=all), and based on inputs by NP.

| **Sno.** | **Order** | **Family** | **Genus** | **Species** | **Endemicity** |
|---|---|---|---|---|---|
| 1 | Apiales | Araliaceae | *Schefflera* | *Schefflera capitata* | 1 |
| 2 | Apiales | Araliaceae | *Schefflera* | *Schefflera micrantha* | 1 |
| 3 | Apiales | Araliaceae | *Schefflera* | *Schefflera racemosa* | 1 |
| 4 | Apiales | Araliaceae | *Schefflera* | *Schefflera venulosa* | 0 |
| 5 | Apiales | Pittosporaceae | *Pittosporum* | *Pittosporum dasycaulon* | 1 |
| 6 | Apiales | Pittosporaceae | *Pittosporum* | *Pittosporum neelgherrense* | 1 |
| 7 | Aquifoliales | Stemonuraceae | *Gomphandra* | *Gomphandra coriacea* | 1 |
| 8 | Aquifoliales | Stemonuraceae | *Gomphandra* | *Gomphandra tetrandra* | 0 |
| 9 | Arecales | Arecaceae | *Arenga* | *Arenga wightii* | 1 |
| 10 | Arecales | Arecaceae | *Caryota* | *Caryota urens* | 0 |
| 11 | Arecales | Arecaceae | *Corypha* | *Corypha umbraculifera* | 0 |
| 12 | Arecales | Arecaceae | *Pinanga* | *Pinanga dicksonii* | 1 |
| 13 | Asparagales | Asparagaceae | *Dracaena* | *Dracaena elliptica* | 0 |
| 14 | Asterales | Asteraceae | *Monosis* | *Monosis travancorica* | 1 |
| 15 | Asterales | Asteraceae | *Monosis* | *Monosis wightiana* | 0 |
| 16 | Boraginales | Boraginaceae | *Ehretia* | *Ehretia canarensis* | 0 |
| 17 | Brassicales | Capparaceae | *Capparis* | *Capparis baducca* | 1 |
| 18 | Buxales | Buxaceae | *Sarcococca* | *Sarcococca pruniformis* | 0 |
| 19 | Celastrales | Celastraceae | *Cassine* | *Cassine paniculata* | 0 |
| 20 | Celastrales | Celastraceae | *Euonymus* | *Euonymus angulatus* | 1 |
| 21 | Celastrales | Celastraceae | *Euonymus* | *Euonymus dichotomus* | 1 |
| 22 | Celastrales | Celastraceae | *Euonymus* | *Euonymus indicus* | 1 |
| 23 | Celastrales | Celastraceae | *Glyptopetalum* | *Glyptopetalum grandiflorum* | 1 |
| 24 | Celastrales | Celastraceae | *Lophopetalum* | *Lophopetalum wightianum* | 0 |

| | | | | | |
|---|---|---|---|---|---|
| 25 | Celastrales | Celastraceae | *Maytenus* | *Maytenus rothiana* | 1 |
| 26 | Celastrales | Celastraceae | *Microtropis* | *Microtropis latifolia* | 1 |
| 27 | Celastrales | Celastraceae | *Microtropis* | *Microtropis stocksii* | 1 |
| 28 | Cornales | Nyssaceae | *Mastixia* | *Mastixia arborea* | 1 |
| 29 | Crossosomatales | Staphyleaceae | *Turpinia* | *Turpinia cochinchinensis* | 0 |
| 30 | Crossosomatales | Staphyleaceae | *Turpinia* | *Turpinia malabarica* | 1 |
| 31 | Cucurbitales | Tetramelaceae | *Tetrameles* | *Tetrameles nudiflora* | 0 |
| 32 | Dilleniales | Dilleniaceae | *Dillenia* | *Dillenia bracteata* | 1 |
| 33 | Dipsacales | Adoxaceae | *Viburnum* | *Viburnum punctatum* | 0 |
| 34 | Ericales | Ebenaceae | *Diospyros* | *Diospyros barberi* | 0 |
| 35 | Ericales | Ebenaceae | *Diospyros* | *Diospyros bourdillonii* | 1 |
| 36 | Ericales | Ebenaceae | *Diospyros* | *Diospyros buxifolia* | 0 |
| 37 | Ericales | Ebenaceae | *Diospyros* | *Diospyros candolleana* | 1 |
| 38 | Ericales | Ebenaceae | *Diospyros* | *Diospyros crumenata* | 0 |
| 39 | Ericales | Ebenaceae | *Diospyros* | *Diospyros ebenum* | 0 |
| 40 | Ericales | Ebenaceae | *Diospyros* | *Diospyros foliolosa* | 1 |
| 41 | Ericales | Ebenaceae | *Diospyros* | *Diospyros ghatensis* | 1 |
| 42 | Ericales | Ebenaceae | *Diospyros* | *Diospyros insignis* | 0 |
| 43 | Ericales | Ebenaceae | *Diospyros* | *Diospyros kedam* | 1 |
| 44 | Ericales | Ebenaceae | *Diospyros* | *Diospyros malabarica* | 0 |
| 45 | Ericales | Ebenaceae | *Diospyros* | *Diospyros montana* | 1 |
| 46 | Ericales | Ebenaceae | *Diospyros* | *Diospyros neilgerrensis* | 0 |
| 47 | Ericales | Ebenaceae | *Diospyros* | *Diospyros nilagirica* | 1 |
| 48 | Ericales | Ebenaceae | *Diospyros* | *Diospyros oocarpa* | 1 |
| 49 | Ericales | Ebenaceae | *Diospyros* | *Diospyros paniculata* | 1 |
| 50 | Ericales | Ebenaceae | *Diospyros* | *Diospyros pruriens* | 0 |
| 51 | Ericales | Ebenaceae | *Diospyros* | *Diospyros ridleyi* | 0 |
| 52 | Ericales | Ebenaceae | *Diospyros* | *Diospyros saldanhae* | 1 |
| 53 | Ericales | Ebenaceae | *Diospyros* | *Diospyros sylvatica* | 1 |
| 54 | Ericales | Ebenaceae | *Diospyros* | *Diospyros vera* | 1 |
| 55 | Ericales | Pentaphylacaceae | *Eurya* | *Eurya nitida* | 0 |

| | | | | | |
|---|---|---|---|---|---|
| 56 | Ericales | Primulaceae | *Ardisia* | *Ardisia missionis* | 1 |
| 57 | Ericales | Primulaceae | *Ardisia* | *Ardisia pauciflora* | 1 |
| 58 | Ericales | Primulaceae | *Ardisia* | *Ardisia rhomboidea* | 1 |
| 59 | Ericales | Primulaceae | *Ardisia* | *Ardisia solanacea* | 0 |
| 60 | Ericales | Primulaceae | *Ardisia* | *Ardisia stonei* | 1 |
| 61 | Ericales | Primulaceae | *Maesa* | *Maesa indica* | 0 |
| 62 | Ericales | Primulaceae | *Rapanea* | *Rapanea wightiana* | 0 |
| 63 | Ericales | Sapotaceae | *Chrysophyllum* | *Chrysophyllum flexuosum* | 0 |
| 64 | Ericales | Sapotaceae | *Chrysophyllum* | *Chrysophyllum roxburghii* | 0 |
| 65 | Ericales | Sapotaceae | *Isonandra* | *Isonandra lanceolata* | 0 |
| 66 | Ericales | Sapotaceae | *Isonandra* | *Isonandra perrottetiana* | 1 |
| 67 | Ericales | Sapotaceae | *Madhuca* | *Madhuca bourdillonii* | 1 |
| 68 | Ericales | Sapotaceae | *Madhuca* | *Madhuca neriifolia* | 1 |
| 69 | Ericales | Sapotaceae | *Mimusops* | *Mimusops elengi* | 0 |
| 70 | Ericales | Sapotaceae | *Palaquium* | *Palaquium bourdillonii* | 1 |
| 71 | Ericales | Sapotaceae | *Palaquium* | *Palaquium ellipticum* | 1 |
| 72 | Ericales | Sapotaceae | *Xantolis* | *Xantolis tomentosa* | 0 |
| 73 | Ericales | Symplocaceae | *Symplocos* | *Symplocos cochinchinensis* | 0 |
| 74 | Ericales | Symplocaceae | *Symplocos* | *Symplocos macrophylla* | 1 |
| 75 | Ericales | Symplocaceae | *Symplocos* | *Symplocos monantha* | 1 |
| 76 | Ericales | Symplocaceae | *Symplocos* | *Symplocos pulchra* | 1 |
| 77 | Ericales | Symplocaceae | *Symplocos* | *Symplocos racemosa* | 0 |
| 78 | Ericales | Symplocaceae | *Symplocos* | *Symplocos rosea* | 1 |
| 79 | Ericales | Symplocaceae | *Symplocos* | *Symplocos wynadense* | 1 |
| 80 | Ericales | Theaceae | *Gordonia* | *Gordonia obtusa* | 1 |
| 81 | Fabales | Fabaceae | *Acrocarpus* | *Acrocarpus fraxinifolius* | 0 |
| 82 | Fabales | Fabaceae | *Archidendron* | *Archidendron bigeminum* | 0 |
| 83 | Fabales | Fabaceae | *Cynometra* | *Cynometra bourdillonii* | 1 |
| 84 | Fabales | Fabaceae | *Cynometra* | *Cynometra travancorica* | 1 |
| 85 | Fabales | Fabaceae | *Humboldtia* | *Humboldtia brunonis* | 1 |
| 86 | Fabales | Fabaceae | *Humboldtia* | *Humboldtia decurrens* | 1 |

| | | | | | |
|---|---|---|---|---|---|
| 87 | Fabales | Fabaceae | *Humboldtia* | *Humboldtia vahliana* | 1 |
| 88 | Fabales | Fabaceae | *Kingiodendron* | *Kingiodendron pinnatum* | 1 |
| 89 | Fabales | Fabaceae | *Ormosia* | *Ormosia travancorica* | 1 |
| 90 | Fabales | Fabaceae | *Saraca* | *Saraca asoca* | 0 |
| 91 | Fabales | Polygalaceae | *Xanthophyllum* | *Xanthophyllum arnottianum* | 1 |
| 92 | Gentianales | Apocynaceae | *Alstonia* | *Alstonia scholaris* | 0 |
| 93 | Gentianales | Apocynaceae | *Hunteria* | *Hunteria zeylanica* | 0 |
| 94 | Gentianales | Apocynaceae | *Tabernaemontana* | *Tabernaemontana alternifolia* | 1 |
| 95 | Gentianales | Apocynaceae | *Tabernaemontana* | *Tabernaemontana gamblei* | 1 |
| 96 | Gentianales | Gentianaceae | *Fagraea* | *Fagraea ceilanica* | 0 |
| 97 | Gentianales | Rubiaceae | *Aidia* | *Aidia densiflora* | 0 |
| 98 | Gentianales | Rubiaceae | *Canthium* | *Canthium travancoricum* | 1 |
| 99 | Gentianales | Rubiaceae | *Chassalia* | *Chassalia curviflora* | 0 |
| 100 | Gentianales | Rubiaceae | *Coffea* | *Coffea wightiana* | 1 |
| 101 | Gentianales | Rubiaceae | *Discospermum* | *Discospermum apiocarpum* | 1 |
| 102 | Gentianales | Rubiaceae | *Discospermum* | *Discospermum sphaerocarpum* | 1 |
| 103 | Gentianales | Rubiaceae | *Ixora* | *Ixora alba* | 1 |
| 104 | Gentianales | Rubiaceae | *Ixora* | *Ixora anamalayana* | 1 |
| 105 | Gentianales | Rubiaceae | *Ixora* | *Ixora brachiata* | 1 |
| 106 | Gentianales | Rubiaceae | *Ixora* | *Ixora elongata* | 1 |
| 107 | Gentianales | Rubiaceae | *Ixora* | *Ixora lanceolaria* | 1 |
| 108 | Gentianales | Rubiaceae | *Ixora* | *Ixora nigricans* | 0 |
| 109 | Gentianales | Rubiaceae | *Ixora* | *Ixora polyantha* | 1 |
| 110 | Gentianales | Rubiaceae | *Ixora* | *Ixora sp* | 1 |
| 111 | Gentianales | Rubiaceae | *Lasianthus* | *Lasianthus acuminatus* | 1 |
| 112 | Gentianales | Rubiaceae | *Lasianthus* | *Lasianthus blumeanus* | 1 |
| 113 | Gentianales | Rubiaceae | *Lasianthus* | *Lasianthus capitulatus* | 1 |
| 114 | Gentianales | Rubiaceae | *Lasianthus* | *Lasianthus jackianus* | 1 |
| 115 | Gentianales | Rubiaceae | *Lasianthus* | *Lasianthus oblongifolius* | 1 |
| 116 | Gentianales | Rubiaceae | *Lasianthus* | *Lasianthus parvifolius* | 1 |
| 117 | Gentianales | Rubiaceae | *Lasianthus* | *Lasianthus rostratus* | 1 |

| | | | | | |
|---|---|---|---|---|---|
| 118 | Gentianales | Rubiaceae | *Meyna* | *Meyna laxiflora* | 0 |
| 119 | Gentianales | Rubiaceae | *Mitragyna* | *Mitragyna parvifolia* | 0 |
| 120 | Gentianales | Rubiaceae | *Mitragyna* | *Mitragyna tubulosa* | 1 |
| 121 | Gentianales | Rubiaceae | *Octotropis* | *Octotropis travancorica* | 1 |
| 122 | Gentianales | Rubiaceae | *Pavetta* | *Pavetta indica* | 0 |
| 123 | Gentianales | Rubiaceae | *Pavetta* | *Pavetta sp* | 1 |
| 124 | Gentianales | Rubiaceae | *Prismatomeris* | *Prismatomeris tetrandra* | 0 |
| 125 | Gentianales | Rubiaceae | *Psychotria* | *Psychotria anamallayana* | 1 |
| 126 | Gentianales | Rubiaceae | *Psychotria* | *Psychotria connata* | 1 |
| 127 | Gentianales | Rubiaceae | *Psychotria* | *Psychotria dalzellii* | 1 |
| 128 | Gentianales | Rubiaceae | *Psychotria* | *Psychotria flavida* | 1 |
| 129 | Gentianales | Rubiaceae | *Psychotria* | *Psychotria macrocarpa* | 1 |
| 130 | Gentianales | Rubiaceae | *Psychotria* | *Psychotria nigra* | 1 |
| 131 | Gentianales | Rubiaceae | *Psychotria* | *Psychotria sp* | 1 |
| 132 | Gentianales | Rubiaceae | *Psychotria* | *Psychotria truncata* | 1 |
| 133 | Gentianales | Rubiaceae | *Psydrax* | *Psydrax dicoccos* | 0 |
| 134 | Gentianales | Rubiaceae | *Psydrax* | *Psydrax pergracilis* | 1 |
| 135 | Gentianales | Rubiaceae | *Saprosma* | *Saprosma corymbosum* | 1 |
| 136 | Gentianales | Rubiaceae | *Saprosma* | *Saprosma glomerata* | 1 |
| 137 | Gentianales | Rubiaceae | *Tarenna* | *Tarenna asiatica* | 0 |
| 138 | Gentianales | Rubiaceae | *Tarenna* | *Tarenna nilagirica* | 1 |
| 139 | Gentianales | Rubiaceae | *Wendlandia* | *Wendlandia thyrsoidea* | 1 |
| 140 | Icacinales | Icacinaceae | *Nothapodytes* | *Nothapodytes nimmoniana* | 0 |
| 141 | Lamiales | Bignoniaceae | *Pajanelia* | *Pajanelia longifolia* | 0 |
| 142 | Lamiales | Bignoniaceae | *Stereospermum* | *Stereospermum tetragonum* | 1 |
| 143 | Lamiales | Lamiaceae | *Callicarpa* | *Callicarpa tomentosa* | 0 |
| 144 | Lamiales | Lamiaceae | *Clerodendrum* | *Clerodendrum infortunatum* | 0 |
| 145 | Lamiales | Lamiaceae | *Vitex* | *Vitex altissima* | 0 |
| 146 | Lamiales | Oleaceae | *Chionanthus* | *Chionanthus courtallensis* | 1 |
| 147 | Lamiales | Oleaceae | *Chionanthus* | *Chionanthus linocieroides* | 1 |
| 148 | Lamiales | Oleaceae | *Chionanthus* | *Chionanthus mala-elengi* | 1 |

| | | | | | |
|---|---|---|---|---|---|
| 149 | Lamiales | Oleaceae | *Chionanthus* | *Chionanthus ramiflorus* | 0 |
| 150 | Lamiales | Oleaceae | *Ligustrum* | *Ligustrum perrottetii* | 1 |
| 151 | Lamiales | Oleaceae | *Olea* | *Olea dioica* | 1 |
| 152 | Lamiales | Oleaceae | *Olea* | *Olea paniculata* | 0 |
| 153 | Laurales | Lauraceae | *Actinodaphne* | *Actinodaphne bourdillonii* | 1 |
| 154 | Laurales | Lauraceae | *Actinodaphne* | *Actinodaphne campanulata* | 1 |
| 155 | Laurales | Lauraceae | *Actinodaphne* | *Actinodaphne gullavara* | 0 |
| 156 | Laurales | Lauraceae | *Actinodaphne* | *Actinodaphne hookeri* | 1 |
| 157 | Laurales | Lauraceae | *Actinodaphne* | *Actinodaphne lawsonii* | 1 |
| 158 | Laurales | Lauraceae | *Actinodaphne* | *Actinodaphne salicina* | 1 |
| 159 | Laurales | Lauraceae | *Actinodaphne* | *Actinodaphne tadulingamii* | 1 |
| 160 | Laurales | Lauraceae | *Actinodaphne* | *Actinodaphne wightiana* | 1 |
| 161 | Laurales | Lauraceae | *Alseodaphne* | *Alseodaphne semecarpifolia* | 0 |
| 162 | Laurales | Lauraceae | *Apollonias* | *Apollonias arnottii* | 1 |
| 163 | Laurales | Lauraceae | *Beilschmiedia* | *Beilschmiedia dalzellii* | 0 |
| 164 | Laurales | Lauraceae | *Beilschmiedia* | *Beilschmiedia gemmiflora* | 1 |
| 165 | Laurales | Lauraceae | *Beilschmiedia* | *Beilschmiedia wightii* | 1 |
| 166 | Laurales | Lauraceae | *Cinnamomum* | *Cinnamomum cassia* | 1 |
| 167 | Laurales | Lauraceae | *Cinnamomum* | *Cinnamomum filipedicellatum* | 1 |
| 168 | Laurales | Lauraceae | *Cinnamomum* | *Cinnamomum keralaense* | 1 |
| 169 | Laurales | Lauraceae | *Cinnamomum* | *Cinnamomum macrocarpum* | 1 |
| 170 | Laurales | Lauraceae | *Cinnamomum* | *Cinnamomum malabatrum* | 1 |
| 171 | Laurales | Lauraceae | *Cinnamomum* | *Cinnamomum sulphuratum* | 1 |
| 172 | Laurales | Lauraceae | *Cinnamomum* | *Cinnamomum verum* | 1 |
| 173 | Laurales | Lauraceae | *Cryptocarya* | *Cryptocarya anamalayana* | 1 |
| 174 | Laurales | Lauraceae | *Cryptocarya* | *Cryptocarya beddomei* | 1 |
| 175 | Laurales | Lauraceae | *Cryptocarya* | *Cryptocarya lawsonii* | 1 |
| 176 | Laurales | Lauraceae | *Cryptocarya* | *Cryptocarya stocksii* | 1 |
| 177 | Laurales | Lauraceae | *Cryptocarya* | *Cryptocarya wightiana* | 0 |
| 178 | Laurales | Lauraceae | *Litsea* | *Litsea floribunda* | 1 |
| 179 | Laurales | Lauraceae | *Litsea* | *Litsea ghatica* | 1 |

| | | | | | |
|---|---|---|---|---|---|
| 180 | Laurales | Lauraceae | *Litsea* | *Litsea glabrata* | 1 |
| 181 | Laurales | Lauraceae | *Litsea* | *Litsea keralana* | 1 |
| 182 | Laurales | Lauraceae | *Litsea* | *Litsea laevigata* | 1 |
| 183 | Laurales | Lauraceae | *Litsea* | *Litsea mysorensis* | 1 |
| 184 | Laurales | Lauraceae | *Litsea* | *Litsea oleoides* | 1 |
| 185 | Laurales | Lauraceae | *Litsea* | *Litsea stocksii* | 1 |
| 186 | Laurales | Lauraceae | *Litsea* | *Litsea travancorica* | 1 |
| 187 | Laurales | Lauraceae | *Litsea* | *Litsea udayanii* | 1 |
| 188 | Laurales | Lauraceae | *Litsea* | *Litsea UM* | 1 |
| 189 | Laurales | Lauraceae | *Litsea* | *Litsea venulosa* | 1 |
| 190 | Laurales | Lauraceae | *Litsea* | *Litsea wightiana* | 1 |
| 191 | Laurales | Lauraceae | *Neolitsea* | *Neolitsea fischeri* | 1 |
| 192 | Laurales | Lauraceae | *Neolitsea* | *Neolitsea scrobiculata* | 0 |
| 193 | Laurales | Lauraceae | *Neolitsea* | *Neolitsea zeylanica* | 0 |
| 194 | Laurales | Lauraceae | *Ocotea* | *Ocotea lancifolia* | 0 |
| 195 | Laurales | Lauraceae | *Persea* | *Persea macrantha* | 0 |
| 196 | Magnoliales | Annonaceae | *Alphonsea* | *Alphonsea lutea* | 0 |
| 197 | Magnoliales | Annonaceae | *Cyathocalyx* | *Cyathocalyx zeylanicus* | 1 |
| 198 | Magnoliales | Annonaceae | *Goniothalamus* | *Goniothalamus cardiopetalus* | 1 |
| 199 | Magnoliales | Annonaceae | *Goniothalamus* | *Goniothalamus rhynchantherus* | 1 |
| 200 | Magnoliales | Annonaceae | *Goniothalamus* | *Goniothalamus thwaitesii* | 0 |
| 201 | Magnoliales | Annonaceae | *Goniothalamus* | *Goniothalamus wightii* | 1 |
| 202 | Magnoliales | Annonaceae | *Goniothalamus* | *Goniothalamus wynaadensis* | 1 |
| 203 | Magnoliales | Annonaceae | *Meiogyne* | *Meiogyne pannosa* | 1 |
| 204 | Magnoliales | Annonaceae | *Meiogyne* | *Meiogyne ramarowii* | 1 |
| 205 | Magnoliales | Annonaceae | *Miliusa* | *Miliusa gokhalaei* | 1 |
| 206 | Magnoliales | Annonaceae | *Miliusa* | *Miliusa nilagirica* | 1 |
| 207 | Magnoliales | Annonaceae | *Miliusa* | *Miliusa wightiana* | 1 |
| 208 | Magnoliales | Annonaceae | *Mitrephora* | *Mitrephora grandiflora* | 1 |
| 209 | Magnoliales | Annonaceae | *Mitrephora* | *Mitrephora heyneana* | 0 |
| 210 | Magnoliales | Annonaceae | *Orophea* | *Orophea erythrocarpa* | 1 |

| | | | | | |
|---|---|---|---|---|---|
| 211 | Magnoliales | Annonaceae | *Orophea* | *Orophea malabarica* | 1 |
| 212 | Magnoliales | Annonaceae | *Orophea* | *Orophea sivarajanii* | 1 |
| 213 | Magnoliales | Annonaceae | *Orophea* | *Orophea thomsonii* | 1 |
| 214 | Magnoliales | Annonaceae | *Orophea* | *Orophea zeylanica* | 1 |
| 215 | Magnoliales | Annonaceae | *Polyalthia* | *Polyalthia coffeoides* | 1 |
| 216 | Magnoliales | Annonaceae | *Polyalthia* | *Polyalthia fragrans* | 1 |
| 217 | Magnoliales | Annonaceae | *Polyalthia* | *Polyalthia malabarica* | 1 |
| 218 | Magnoliales | Annonaceae | *Polyalthia* | *Polyalthia shendurunii* | 1 |
| 219 | Magnoliales | Annonaceae | *Sageraea* | *Sageraea laurina* | 1 |
| 220 | Magnoliales | Annonaceae | *Sageraea* | *Sageraea thwaitesii* | 1 |
| 221 | Magnoliales | Annonaceae | *Xylopia* | *Xylopia patoniae* | 1 |
| 222 | Magnoliales | Magnoliaceae | *Magnolia* | *Magnolia champaca* | 0 |
| 223 | Magnoliales | Magnoliaceae | *Magnolia* | *Magnolia nilagirica* | 1 |
| 224 | Magnoliales | Myristicaceae | *Gymnacranthera* | *Gymnacranthera canarica* | 1 |
| 225 | Magnoliales | Myristicaceae | *Knema* | *Knema attenuata* | 1 |
| 226 | Magnoliales | Myristicaceae | *Myristica* | *Myristica beddomei* | 1 |
| 227 | Magnoliales | Myristicaceae | *Myristica* | *Myristica malabarica* | 1 |
| 228 | Malpighiales | Calophyllaceae | *Calophyllum* | *Calophyllum apetalum* | 1 |
| 229 | Malpighiales | Calophyllaceae | *Calophyllum* | *Calophyllum austroindicum* | 1 |
| 230 | Malpighiales | Calophyllaceae | *Calophyllum* | *Calophyllum tomentosum* | 0 |
| 231 | Malpighiales | Calophyllaceae | *Mammea* | *Mammea suriga* | 1 |
| 232 | Malpighiales | Calophyllaceae | *Mesua* | *Mesua ferrea* | 1 |
| 233 | Malpighiales | Calophyllaceae | *Poeciloneuron* | *Poeciloneuron indicum* | 1 |
| 234 | Malpighiales | Centroplacaceae | *Bhesa* | *Bhesa indica* | 0 |
| 235 | Malpighiales | Chrysobalanaceae | *Atuna* | *Atuna indica* | 1 |
| 236 | Malpighiales | Chrysobalanaceae | *Atuna* | *Atuna travancorica* | 1 |
| 237 | Malpighiales | Clusiaceae | *Agasthiyamalai* | *Agasthiyamalai pauciflora* | 1 |
| 238 | Malpighiales | Clusiaceae | *Garcinia* | *Garcinia gummi-gutta* | 1 |
| 239 | Malpighiales | Clusiaceae | *Garcinia* | *Garcinia indica* | 1 |
| 240 | Malpighiales | Clusiaceae | *Garcinia* | *Garcinia morella* | 0 |
| 241 | Malpighiales | Clusiaceae | *Garcinia* | *Garcinia rubroechinata* | 1 |

| | | | | | |
|---|---|---|---|---|---|
| 242 | Malpighiales | Clusiaceae | *Garcinia* | *Garcinia spicata* | 0 |
| 243 | Malpighiales | Clusiaceae | *Garcinia* | *Garcinia talbotii* | 1 |
| 244 | Malpighiales | Dichapetalaceae | *Dichapetalum* | *Dichapetalum gelonioides* | 0 |
| 245 | Malpighiales | Erythroxylaceae | *Erythroxylum* | *Erythroxylum moonii* | 1 |
| 246 | Malpighiales | Erythroxylaceae | *Erythroxylum* | *Erythroxylum obtusifolium* | 1 |
| 247 | Malpighiales | Euphorbiaceae | *Agrostistachys* | *Agrostistachys borneensis* | 0 |
| 248 | Malpighiales | Euphorbiaceae | *Agrostistachys* | *Agrostistachys indica* | 0 |
| 249 | Malpighiales | Euphorbiaceae | *Blachia* | *Blachia calycina* | 1 |
| 250 | Malpighiales | Euphorbiaceae | *Blachia* | *Blachia denudata* | 1 |
| 251 | Malpighiales | Euphorbiaceae | *Blachia* | *Blachia umbellata* | 0 |
| 252 | Malpighiales | Euphorbiaceae | *Cleidion* | *Cleidion javanicum* | 0 |
| 253 | Malpighiales | Euphorbiaceae | *Croton* | *Croton chakrabartyi* | 1 |
| 254 | Malpighiales | Euphorbiaceae | *Croton* | *Croton klotzschianus* | 1 |
| 255 | Malpighiales | Euphorbiaceae | *Croton* | *Croton laccifer* | 0 |
| 256 | Malpighiales | Euphorbiaceae | *Croton* | *Croton malabaricus* | 1 |
| 257 | Malpighiales | Euphorbiaceae | *Croton* | *Croton zeylanicus* | 1 |
| 258 | Malpighiales | Euphorbiaceae | *Dimorphocalyx* | *Dimorphocalyx beddomei* | 1 |
| 259 | Malpighiales | Euphorbiaceae | *Dimorphocalyx* | *Dimorphocalyx glabellus* | 1 |
| 260 | Malpighiales | Euphorbiaceae | *Epiprinus* | *Epiprinus mallotiformis* | 1 |
| 261 | Malpighiales | Euphorbiaceae | *Excoecaria* | *Excoecaria oppositifolia* | 1 |
| 262 | Malpighiales | Euphorbiaceae | *Falconeria* | *Falconeria insignis* | 0 |
| 263 | Malpighiales | Euphorbiaceae | *Macaranga* | *Macaranga indica* | 0 |
| 264 | Malpighiales | Euphorbiaceae | *Macaranga* | *Macaranga peltata* | 0 |
| 265 | Malpighiales | Euphorbiaceae | *Mallotus* | *Mallotus aureopunctatus* | 1 |
| 266 | Malpighiales | Euphorbiaceae | *Mallotus* | *Mallotus beddomei* | 1 |
| 267 | Malpighiales | Euphorbiaceae | *Mallotus* | *Mallotus distans* | 1 |
| 268 | Malpighiales | Euphorbiaceae | *Mallotus* | *Mallotus nudiflorus* | 0 |
| 269 | Malpighiales | Euphorbiaceae | *Mallotus* | *Mallotus philippensis* | 0 |
| 270 | Malpighiales | Euphorbiaceae | *Mallotus* | *Mallotus resinosus* | 1 |
| 271 | Malpighiales | Euphorbiaceae | *Mallotus* | *Mallotus rhamnifolius* | 0 |
| 272 | Malpighiales | Euphorbiaceae | *Mallotus* | *Mallotus tetracoccus* | 0 |

| | | | | | |
|---|---|---|---|---|---|
| 273 | Malpighiales | Euphorbiaceae | *Paracroton* | *Paracroton pendulus subsp. zeylanicus* | 1 |
| 274 | Malpighiales | Flacourtiaceae | *Hydnocarpus* | *Hydnocarpus alpina* | 1 |
| 275 | Malpighiales | Flacourtiaceae | *Hydnocarpus* | *Hydnocarpus macrocarpa* | 1 |
| 276 | Malpighiales | Flacourtiaceae | *Hydnocarpus* | *Hydnocarpus pentandrus* | 1 |
| 277 | Malpighiales | Ochnaceae | *Gomphia* | *Gomphia serrata* | 0 |
| 278 | Malpighiales | Phyllanthaceae | *Actephila* | *Actephila excelsa* | 0 |
| 279 | Malpighiales | Phyllanthaceae | *Antidesma* | *Antidesma alexiteria* | 1 |
| 280 | Malpighiales | Phyllanthaceae | *Antidesma* | *Antidesma montanum* | 0 |
| 281 | Malpighiales | Phyllanthaceae | *Aporosa* | *Aporosa indo-acuminata* | 1 |
| 282 | Malpighiales | Phyllanthaceae | *Aporosa* | *Aporosa bourdillonii* | 1 |
| 283 | Malpighiales | Phyllanthaceae | *Aporosa* | *Aporosa cardiosperma* | 0 |
| 284 | Malpighiales | Phyllanthaceae | *Baccaurea* | *Baccaurea courtallensis* | 1 |
| 285 | Malpighiales | Phyllanthaceae | *Bischofia* | *Bischofia javanica* | 0 |
| 286 | Malpighiales | Phyllanthaceae | *Breynia* | *Breynia vitis-idaea* | 0 |
| 287 | Malpighiales | Phyllanthaceae | *Cleistanthus* | *Cleistanthus malabaricus* | 1 |
| 288 | Malpighiales | Phyllanthaceae | *Cleistanthus* | *Cleistanthus travancorensis* | 1 |
| 289 | Malpighiales | Phyllanthaceae | *Glochidion* | *Glochidion ellipticum* | 0 |
| 290 | Malpighiales | Phyllanthaceae | *Glochidion* | *Glochidion heyneanum* | 0 |
| 291 | Malpighiales | Phyllanthaceae | *Glochidion* | *Glochidion hohenackeri* | 1 |
| 292 | Malpighiales | Phyllanthaceae | *Glochidion* | *Glochidion sp (UM)* | 1 |
| 293 | Malpighiales | Phyllanthaceae | *Margaritaria* | *Margaritaria indica* | 0 |
| 294 | Malpighiales | Putranjivaceae | *Drypetes* | *Drypetes confertiflora* | 1 |
| 295 | Malpighiales | Putranjivaceae | *Drypetes* | *Drypetes gardneri* | 1 |
| 296 | Malpighiales | Putranjivaceae | *Drypetes* | *Drypetes malabarica* | 1 |
| 297 | Malpighiales | Putranjivaceae | *Drypetes* | *Drypetes oblongifolia* | 1 |
| 298 | Malpighiales | Putranjivaceae | *Drypetes* | *Drypetes venusta* | 1 |
| 299 | Malpighiales | Putranjivaceae | *Drypetes* | *Drypetes wightii* | 1 |
| 300 | Malpighiales | Rhizophoraceae | *Blepharistemma* | *Blepharistemma serratum* | 1 |
| 301 | Malpighiales | Rhizophoraceae | *Carallia* | *Carallia brachiata* | 0 |
| 302 | Malpighiales | Salicaceae | *Casearia* | *Casearia ovata* | 0 |
| 303 | Malpighiales | Salicaceae | *Casearia* | *Casearia rubescens* | 0 |

| | | | | | |
|---|---|---|---|---|---|
| 304 | Malpighiales | Salicaceae | *Casearia* | *Casearia wynadensis* | 1 |
| 305 | Malpighiales | Salicaceae | *Flacourtia* | *Flacourtia montana* | 1 |
| 306 | Malpighiales | Salicaceae | *Homalium* | *Homalium ceylanicum* | 1 |
| 307 | Malpighiales | Salicaceae | *Scolopia* | *Scolopia crenata* | 0 |
| 308 | Malpighiales | Violaceae | *Rinorea* | *Rinorea bengalensis* | 0 |
| 309 | Malvales | Dipterocarpaceae | *Dipterocarpus* | *Dipterocarpus bourdillonii* | 1 |
| 310 | Malvales | Dipterocarpaceae | *Dipterocarpus* | *Dipterocarpus indicus* | 1 |
| 311 | Malvales | Dipterocarpaceae | *Hopea* | *Hopea canarensis* | 1 |
| 312 | Malvales | Dipterocarpaceae | *Hopea* | *Hopea erosa* | 1 |
| 313 | Malvales | Dipterocarpaceae | *Hopea* | *Hopea glabra* | 1 |
| 314 | Malvales | Dipterocarpaceae | *Hopea* | *Hopea parviflora* | 1 |
| 315 | Malvales | Dipterocarpaceae | *Hopea* | *Hopea ponga* | 1 |
| 316 | Malvales | Dipterocarpaceae | *Hopea* | *Hopea racophloea* | 1 |
| 317 | Malvales | Dipterocarpaceae | *Hopea* | *Hopea utilis* | 1 |
| 318 | Malvales | Dipterocarpaceae | *Vateria* | *Vateria indica* | 1 |
| 319 | Malvales | Malvaceae | *Bombax* | *Bombax ceiba* | 0 |
| 320 | Malvales | Malvaceae | *Durio* | *Durio exarillatus* | 1 |
| 321 | Malvales | Malvaceae | *Heritiera* | *Heritiera papilio* | 1 |
| 322 | Malvales | Malvaceae | *Leptonychia* | *Leptonychia caudata* | 0 |
| 323 | Malvales | Malvaceae | *Pterospermum* | *Pterospermum diversifolium* | 0 |
| 324 | Malvales | Malvaceae | *Pterospermum* | *Pterospermum reticulatum* | 1 |
| 325 | Malvales | Malvaceae | *Pterospermum* | *Pterospermum rubiginosum* | 1 |
| 326 | Malvales | Malvaceae | *Pterygota* | *Pterygota alata* | 0 |
| 327 | Malvales | Malvaceae | *Sterculia* | *Sterculia guttata* | 0 |
| 328 | Malvales | Thymelaeaceae | *Gnidia* | *Gnidia glauca* | 0 |
| 329 | Metteniusales | Metteniusaceae | *Apodytes* | *Apodytes dimidiata* | 0 |
| 330 | Myrtales | Combretaceae | *Terminalia* | *Terminalia bellirica* | 0 |
| 331 | Myrtales | Combretaceae | *Terminalia* | *Terminalia chebula* | 0 |
| 332 | Myrtales | Combretaceae | *Terminalia* | *Terminalia paniculata* | 1 |
| 333 | Myrtales | Combretaceae | *Terminalia* | *Terminalia travancorensis* | 1 |
| 334 | Myrtales | Lythraceae | *Lagerstroemia* | *Lagerstroemia microcarpa* | 1 |

| | | | | | |
|---|---|---|---|---|---|
| 335 | Myrtales | Melastomataceae | *Memecylon* | *Memecylon angustifolium* | 0 |
| 336 | Myrtales | Melastomataceae | *Memecylon* | *Memecylon depressum* | 1 |
| 337 | Myrtales | Melastomataceae | *Memecylon* | *Memecylon gracile* | 1 |
| 338 | Myrtales | Melastomataceae | *Memecylon* | *Memecylon heyneanum* | 1 |
| 339 | Myrtales | Melastomataceae | *Memecylon* | *Memecylon malabaricum* | 1 |
| 340 | Myrtales | Melastomataceae | *Memecylon* | *Memecylon pseudogracile* | 1 |
| 341 | Myrtales | Melastomataceae | *Memecylon* | *Memecylon randerianum* | 1 |
| 342 | Myrtales | Melastomataceae | *Memecylon* | *Memecylon subramanii* | 1 |
| 343 | Myrtales | Melastomataceae | *Memecylon* | *Memecylon subsessile* | 1 |
| 344 | Myrtales | Melastomataceae | *Memecylon* | *Memecylon talbotianum* | 1 |
| 345 | Myrtales | Melastomataceae | *Memecylon* | *Memecylon terminale* | 1 |
| 346 | Myrtales | Melastomataceae | *Memecylon* | *Memecylon umbellatum* | 1 |
| 347 | Myrtales | Melastomataceae | *Memecylon* | *Memecylon wightii* | 1 |
| 348 | Myrtales | Myrtaceae | *Eugenia* | *Eugenia aloysii* | 1 |
| 349 | Myrtales | Myrtaceae | *Eugenia* | *Eugenia codyensis* | 1 |
| 350 | Myrtales | Myrtaceae | *Eugenia* | *Eugenia floccosa* | 1 |
| 351 | Myrtales | Myrtaceae | *Eugenia* | *Eugenia kalamii* | 1 |
| 352 | Myrtales | Myrtaceae | *Eugenia* | *Eugenia mabaeoides* | 1 |
| 353 | Myrtales | Myrtaceae | *Eugenia* | *Eugenia macrocalyx* | 1 |
| 354 | Myrtales | Myrtaceae | *Eugenia* | *Eugenia mooniana* | 0 |
| 355 | Myrtales | Myrtaceae | *Eugenia* | *Eugenia rottleriana* | 1 |
| 356 | Myrtales | Myrtaceae | *Eugenia* | *Eugenia singampattiana* | 1 |
| 357 | Myrtales | Myrtaceae | *Meteoromyrtus* | *Meteoromyrtus wynaadensis* | 1 |
| 358 | Myrtales | Myrtaceae | *Syzygium* | *Syzygium benthamianum* | 1 |
| 359 | Myrtales | Myrtaceae | *Syzygium* | *Syzygium calophyllifolium* | 1 |
| 360 | Myrtales | Myrtaceae | *Syzygium* | *Syzygium caryophyllatum* | 1 |
| 361 | Myrtales | Myrtaceae | *Syzygium* | *Syzygium chavaran* | 1 |
| 362 | Myrtales | Myrtaceae | *Syzygium* | *Syzygium cumini* | 0 |
| 363 | Myrtales | Myrtaceae | *Syzygium* | *Syzygium densiflorum* | 1 |
| 364 | Myrtales | Myrtaceae | *Syzygium* | *Syzygium gardneri* | 1 |
| 365 | Myrtales | Myrtaceae | *Syzygium* | *Syzygium grande* | 0 |

| | | | | | |
|---|---|---|---|---|---|
| 366 | Myrtales | Myrtaceae | *Syzygium* | *Syzygium hemisphericum* | 1 |
| 367 | Myrtales | Myrtaceae | *Syzygium* | *Syzygium laetum* | 1 |
| 368 | Myrtales | Myrtaceae | *Syzygium* | *Syzygium lanceolatum* | 1 |
| 369 | Myrtales | Myrtaceae | *Syzygium* | *Syzygium makul* | 1 |
| 370 | Myrtales | Myrtaceae | *Syzygium* | *Syzygium mundagam* | 1 |
| 371 | Myrtales | Myrtaceae | *Syzygium* | *Syzygium munronii* | 1 |
| 372 | Myrtales | Myrtaceae | *Syzygium* | *Syzygium neesianum* | 1 |
| 373 | Myrtales | Myrtaceae | *Syzygium* | *Syzygium palghatense* | 1 |
| 374 | Myrtales | Myrtaceae | *Syzygium* | *Syzygium rubicundum* | 1 |
| 375 | Myrtales | Myrtaceae | *Syzygium* | *Syzygium travancoricum* | 1 |
| 376 | Myrtales | Myrtaceae | *Syzygium* | *Syzygium zeylanicum* | 0 |
| 377 | Oxalidales | Elaeocarpaceae | *Elaeocarpus* | *Elaeocarpus munroii* | 1 |
| 378 | Oxalidales | Elaeocarpaceae | *Elaeocarpus* | *Elaeocarpus serratus* | 1 |
| 379 | Oxalidales | Elaeocarpaceae | *Elaeocarpus* | *Elaeocarpus tuberculatus* | 0 |
| 380 | Oxalidales | Elaeocarpaceae | *Elaeocarpus* | *Elaeocarpus variabilis* | 1 |
| 381 | Oxalidales | Elaeocarpaceae | *Elaeocarpus* | *Elaeocarpus venustus* | 1 |
| 382 | Pandanales | Pandanaceae | *Pandanus* | *Pandanus furcatus* | 0 |
| 383 | Proteales | Proteaceae | *Helicia* | *Helicia nilagirica* | 0 |
| 384 | Ranunculales | Menispermaceae | *Cocculus* | *Cocculus laurifolius* | 0 |
| 385 | Rosales | Cannabaceae | *Aphananthe* | *Aphananthe cuspidata* | 0 |
| 386 | Rosales | Cannabaceae | *Celtis* | *Celtis philippensis* | 0 |
| 387 | Rosales | Cannabaceae | *Celtis* | *Celtis timorensis* | 0 |
| 388 | Rosales | Moraceae | *Antiaris* | *Antiaris toxicaria* | 0 |
| 389 | Rosales | Moraceae | *Artocarpus* | *Artocarpus gomezianus* | 0 |
| 390 | Rosales | Moraceae | *Artocarpus* | *Artocarpus heterophyllus* | 0 |
| 391 | Rosales | Moraceae | *Artocarpus* | *Artocarpus hirsutus* | 1 |
| 392 | Rosales | Moraceae | *Ficus* | *Ficus beddomei* | 1 |
| 393 | Rosales | Moraceae | *Ficus* | *Ficus callosa* | 0 |
| 394 | Rosales | Moraceae | *Ficus* | *Ficus exasperata* | 0 |
| 395 | Rosales | Moraceae | *Ficus* | *Ficus hispida* | 0 |
| 396 | Rosales | Moraceae | *Ficus* | *Ficus microcarpa* | 0 |

| | | | | | |
|---|---|---|---|---|---|
| 397 | Rosales | Moraceae | *Ficus* | *Ficus nervosa* | 0 |
| 398 | Rosales | Moraceae | *Ficus* | *Ficus talbotii* | 0 |
| 399 | Rosales | Moraceae | *Ficus* | *Ficus tsjakela* | 0 |
| 400 | Rosales | Moraceae | *Ficus* | *Ficus virens* | 0 |
| 401 | Rosales | Rosaceae | *Prunus* | *Prunus ceylanica* | 0 |
| 402 | Rosales | Urticaceae | *Boehmeria* | *Boehmeria depauperata* | 0 |
| 403 | Rosales | Urticaceae | *Debregeasia* | *Debregeasia longifolia* | 0 |
| 404 | Rosales | Urticaceae | *Dendrocnide* | *Dendrocnide sinuata* | 0 |
| 405 | Rosales | Urticaceae | *Oreocnide* | *Oreocnide integrifolia* | 0 |
| 406 | Sabiales | Sabiaceae | *Meliosma* | *Meliosma pinnata* | 0 |
| 407 | Sabiales | Sabiaceae | *Meliosma* | *Meliosma simplicifolia* | 0 |
| 408 | Santalales | Olacaceae | *Anacolosa* | *Anacolosa densiflora* | 1 |
| 409 | Santalales | Olacaceae | *Strombosia* | *Strombosia ceylanica* | 1 |
| 410 | Santalales | Santalaceae | *Scleropyrum* | *Scleropyrum pentandrum* | 0 |
| 411 | Sapindales | Anacardiaceae | *Gluta* | *Gluta travancorica* | 1 |
| 412 | Sapindales | Anacardiaceae | *Holigarna* | *Holigarna arnottiana* | 1 |
| 413 | Sapindales | Anacardiaceae | *Holigarna* | *Holigarna beddomei* | 1 |
| 414 | Sapindales | Anacardiaceae | *Holigarna* | *Holigarna ferruginea* | 1 |
| 415 | Sapindales | Anacardiaceae | *Holigarna* | *Holigarna grahamii* | 1 |
| 416 | Sapindales | Anacardiaceae | *Holigarna* | *Holigarna nigra* | 1 |
| 417 | Sapindales | Anacardiaceae | *Mangifera* | *Mangifera indica* | 0 |
| 418 | Sapindales | Anacardiaceae | *Nothopegia* | *Nothopegia aureofulva* | 1 |
| 419 | Sapindales | Anacardiaceae | *Nothopegia* | *Nothopegia beddomei* | 1 |
| 420 | Sapindales | Anacardiaceae | *Nothopegia* | *Nothopegia heyneana* | 0 |
| 421 | Sapindales | Anacardiaceae | *Nothopegia* | *Nothopegia racemosa* | 1 |
| 422 | Sapindales | Anacardiaceae | *Nothopegia* | *Nothopegia travancorica* | 1 |
| 423 | Sapindales | Anacardiaceae | *Semecarpus* | *Semecarpus auriculata* | 1 |
| 424 | Sapindales | Anacardiaceae | *Semecarpus* | *Semecarpus travancorica* | 1 |
| 425 | Sapindales | Anacardiaceae | *Spondias* | *Spondias indica* | 1 |
| 426 | Sapindales | Anacardiaceae | *Spondias* | *Spondias pinnata* | 0 |
| 427 | Sapindales | Burseraceae | *Canarium* | *Canarium strictum* | 0 |

| | | | | | |
|---|---|---|---|---|---|
| 428 | Sapindales | Meliaceae | *Aglaia* | *Aglaia bourdillonii* | 1 |
| 429 | Sapindales | Meliaceae | *Aglaia* | *Aglaia canarana* | 0 |
| 430 | Sapindales | Meliaceae | *Aglaia* | *Aglaia edulis* | 0 |
| 431 | Sapindales | Meliaceae | *Aglaia* | *Aglaia elaeagnoidea* | 0 |
| 432 | Sapindales | Meliaceae | *Aglaia* | *Aglaia lawii* | 0 |
| 433 | Sapindales | Meliaceae | *Aglaia* | *Aglaia perviridis* | 0 |
| 434 | Sapindales | Meliaceae | *Aglaia* | *Aglaia simplicifolia* | 0 |
| 435 | Sapindales | Meliaceae | *Aglaia* | *Aglaia tomentosa* | 0 |
| 436 | Sapindales | Meliaceae | *Aphanamixis* | *Aphanamixis polystachya* | 0 |
| 437 | Sapindales | Meliaceae | *Beddomea* | *Beddomea indica* | 1 |
| 438 | Sapindales | Meliaceae | *Chukrasia* | *Chukrasia tabularis* | 0 |
| 439 | Sapindales | Meliaceae | *Dysoxylum* | *Dysoxylum beddomei* | 1 |
| 440 | Sapindales | Meliaceae | *Dysoxylum* | *Dysoxylum gotadhora* | 0 |
| 441 | Sapindales | Meliaceae | *Dysoxylum* | *Dysoxylum malabaricum* | 1 |
| 442 | Sapindales | Meliaceae | *Heynea* | *Heynea trijuga* | 0 |
| 443 | Sapindales | Meliaceae | *Reinwardtiodendron* | *Reinwardtiodendron anamalaiense* | 1 |
| 444 | Sapindales | Meliaceae | *Toona* | *Toona ciliata* | 0 |
| 445 | Sapindales | Meliaceae | *Turraea* | *Turraea pubescens* | 0 |
| 446 | Sapindales | Meliaceae | *Walsura* | *Walsura trifoliolata* | 1 |
| 447 | Sapindales | Rutaceae | *Acronychia* | *Acronychia pedunculata* | 0 |
| 448 | Sapindales | Rutaceae | *Atalantia* | *Atalantia racemosa* | 0 |
| 449 | Sapindales | Rutaceae | *Atalantia* | *Atalantia wightii* | 1 |
| 450 | Sapindales | Rutaceae | *Clausena* | *Clausena anisata* | 0 |
| 451 | Sapindales | Rutaceae | *Clausena* | *Clausena austroindica* | 1 |
| 452 | Sapindales | Rutaceae | *Clausena* | *Clausena indica* | 1 |
| 453 | Sapindales | Rutaceae | *Glycosmis* | *Glycosmis macrocarpa* | 1 |
| 454 | Sapindales | Rutaceae | *Glycosmis* | *Glycosmis pentaphylla* | 0 |
| 455 | Sapindales | Rutaceae | *Melicope* | *Melicope lunu-ankenda* | 0 |
| 456 | Sapindales | Rutaceae | *Murraya* | *Murraya koenigii* | 0 |
| 457 | Sapindales | Rutaceae | *Murraya* | *Murraya paniculata* | 0 |
| 458 | Sapindales | Rutaceae | *Vepris* | *Vepris bilocularis* | 1 |

| | | | | | |
|---|---|---|---|---|---|
| 459 | Sapindales | Rutaceae | *Zanthoxylum* | *Zanthoxylum rhetsa* | 0 |
| 460 | Sapindales | Sapindaceae | *Allophylus* | *Allophylus cobbe* | 0 |
| 461 | Sapindales | Sapindaceae | *Dimocarpus* | *Dimocarpus longan* | 0 |
| 462 | Sapindales | Sapindaceae | *Dodonaea* | *Dodonaea viscosa* | 0 |
| 463 | Sapindales | Sapindaceae | *Filicium* | *Filicium decipiens* | 0 |
| 464 | Sapindales | Sapindaceae | *Harpullia* | *Harpullia arborea* | 0 |
| 465 | Sapindales | Sapindaceae | *Lepisanthes* | *Lepisanthes erecta* | 0 |
| 466 | Sapindales | Sapindaceae | *Lepisanthes* | *Lepisanthes tetraphylla* | 0 |
| 467 | Sapindales | Sapindaceae | *Otonephelium* | *Otonephelium stipulaceum* | 1 |
| 468 | Sapindales | Simaroubaceae | *Ailanthus* | *Ailanthus triphysa* | 0 |
| 469 | Saxifragales | Daphniphyllaceae | *Daphniphyllum* | *Daphniphyllum neilgherrense* | 1 |
| 470 | Vitales | Vitaceae | *Leea* | *Leea indica* | 0 |
