## Supplementary Table 2 for "Range restricted old and young lineages show the southern Western Ghats to be both a museum and a cradle of diversity for woody plants"

**Appendix S2**

Range restricted old and young lineages show the southern Western Ghats to be both a museum and a cradle of diversity for woody plants

Author list: Abhishek Gopal[1,2*], D. K. Bharti[1✢], Navendu Page[3✢], Kyle G. Dexter[4,5], Ramanathan Krishnamani[6], Ajith Kumar[7], Jahnavi Joshi[1,2*]

Affiliations:
[1] CSIR-Centre for Cellular and Molecular Biology, Uppal Road, Hyderabad
[2] Academy of Scientific and Innovative Research (AcSIR), Ghaziabad, India
[3] Wildlife Institute of India, Dehradun, India
[4] School of GeoSciences, University of Edinburgh, Edinburgh, UK
[5] Tropical Diversity Section, Royal Botanic Garden Edinburgh, Edinburgh, UK
[6] The Rainforest Initiative, Coimbatore, Tamil Nadu, India
[7] Centre for Wildlife Studies, Bangalore, Karnataka, India

[✢]D. K. Bharti and Navendu Page contributed equally to the study.

### S.2. Species distribution modelling

#### S.2.1. Bias layer

A bias layer was created to account for variation in sampling effort across the model extent, which may bias predictions towards areas with greater sampling effort rather than species habitat suitability (Phillips et al., 2009). The bias layer was obtained from a species distribution model for the extent of the Western Ghats using, 1) the occurrence locations of all the species, 2) 19 environmental variables, and elevation data from the WorldClim database (Fick & Hijmans, 2017) as predictor variables, 3) 10,000 random background locations, and 4) default Maxent parameters. The output of this model, the bias layer, consists of habitat suitability predictions, where higher values indicate environments more likely to be sampled and hence represent the sampling bias (Phillips et al., 2009).

#### S.2.2. Model tuning, regularisation and model selection

The continuous predictions at 1 x 1 km resolution were aggregated to 10 x 10 km, and converted into a presence-absence map using a threshold of maximum sum of sensitivity and specificity (Liu et al., 2005). As the default Maxent parameters are unlikely to be appropriate across species, we evaluated models using different combinations of feature classes and tuning parameters for each species (Hallgren et al., 2019; Low et al., 2021; S2.2.). Model evaluation was carried out using test and training datasets obtained from four masked geographically structured partitions for species with at least 20 occurrence locations, and k-1 jackknifing for species with fewer occurrences (Radosavljevic & Anderson, 2014; Shcheglovitova & Anderson, 2013). The best model was chosen by prioritising measures of model transferability over performance (Low et al., 2021).

For each species, we built separate models using six different combinations of feature classes (LQH, LQ, QH, L, Q, H; L–linear, Q–quadratic, H–hinge). These were tuned using ten different regularisation parameters (0.5–5, with an interval of 0.5), as the default Maxent feature classes and regularisation parameters are unlikely to be appropriate across species (Low et al., 2021). Model evaluation included measures of model transferability and performance, namely $AUC_{TEST}$, Area under the receiving operating curve, $OR_{MTP}$, omission rate considering a threshold of minimum training presence, $AUC_{DIFF}$, the difference between AUC test and training points (Low et al., 2021 and references therein). The following sequential filters were used to get the top model, $AUC_{TEST}>0.6$, followed by minimum $OR_{MTP}$, followed by minimum $AUC_{DIFF}$, and maximum $AUC_{TEST}$.

Table S2.1. Model settings & evaluation metrics for the species distribution model run at ~1 x 1 km resolution for 350 woody plant species in the Western Ghats. N- Number of occurrence locations, FC- Feature class combination; Linear (L), Quadratic (Q), Hinge (H), RM: regularization multiplier, $OR_{MTP}$: Omission rate of test presences with a threshold of minimum training presence, $AUC_{TEST}$: average area under the receiver-operator curve (AUC) for test points across partitions, $AUC_{DIFF}$: the difference between the training & the test AUC. The models were selected on the criteria for model transferability and performance. First models with $AUC_{TEST} >$ 0.6 were filtered followed by these sequential criteria: minimum $OR_{MTP}$, minimum $AUC_{DIFF}$ & maximum $AUC_{TEST}$ values. If these criteria resulted in multiple models, the simplest model with the highest regularization value was chosen. See the methods section of the main document for more information.

| **Sno.** | **Species** | **N** | **FC** | **RM** | **$OR_{MTP}$** | **$AUC_{TEST}$** | **$AUC_{DIFF}$** |
|---|---|---|---|---|---|---|---|
| 1 | *Acrocarpus fraxinifolius* | 10 | Q | 1.5 | 0.1 | 0.65 | 0.21 |
| 2 | *Acronychia pedunculata* | 23 | H | 1.5 | 0 | 0.87 | 0.13 |
| 3 | *Actephila excelsa* | 5 | H | 0.5 | 0.2 | 0.84 | 0.19 |
| 4 | *Actinodaphne bourdillonii* | 26 | Q | 2 | 0 | 0.94 | 0.05 |
| 5 | *Actinodaphne gullavara* | 27 | H | 5 | 0 | 0.71 | 0.07 |
| 6 | *Actinodaphne hookeri* | 8 | LQ | 5 | 0.13 | 0.89 | 0.08 |
| 7 | *Actinodaphne tadulingamii* | 14 | LQ | 0.5 | 0.07 | 0.95 | 0.05 |
| 8 | *Actinodaphne wightiana* | 47 | QH | 1 | 0 | 0.91 | 0.05 |
| 9 | *Aglaia bourdillonii* | 7 | LQ | 0.5 | 0.14 | 0.98 | 0.02 |
| 10 | *Aglaia canarana* | 30 | QH | 1.5 | 0 | 0.88 | 0.03 |
| 11 | *Aglaia edulis* | 61 | H | 5 | 0.02 | 0.84 | 0.1 |
| 12 | *Aglaia elaeagnoidea* | 81 | Q | 0.5 | 0.01 | 0.85 | 0.06 |
| 13 | *Aglaia lawii* | 28 | H | 2.5 | 0.07 | 0.68 | 0.15 |
| 14 | *Aglaia perviridis* | 15 | QH | 0.5 | 0.07 | 0.92 | 0.08 |
| 15 | *Aglaia simplicifolia* | 28 | Q | 0.5 | 0.04 | 0.91 | 0.04 |
| 16 | *Agrostistachys borneensis* | 28 | H | 5 | 0.04 | 0.92 | 0.04 |
| 17 | *Agrostistachys indica* | 21 | H | 2.5 | 0 | 0.87 | 0.02 |
| 18 | *Aidia densiflora* | 4 | H | 1 | 0.25 | 0.86 | 0.15 |
| 19 | *Allophylus cobbe* | 10 | H | 4.5 | 0 | 0.94 | 0.04 |

| | | | | | | | |
|---|---|---|---|---|---|---|---|
| 20 | *Alseodaphne semecarpifolia* | 26 | H | 2 | 0.04 | 0.65 | 0.18 |
| 21 | *Alstonia scholaris* | 30 | H | 2 | 0.25 | 0.66 | 0.17 |
| 22 | *Anacolosa densiflora* | 16 | QH | 2 | 0.06 | 0.91 | 0.07 |
| 23 | *Antiaris toxicaria* | 17 | H | 1 | 0.06 | 0.9 | 0.07 |
| 24 | *Antidesma montanum* | 59 | LQH | 3.5 | 0.02 | 0.86 | 0.03 |
| 25 | *Aphanamixis polystachya* | 32 | L | 4 | 0 | 0.83 | 0.08 |
| 26 | *Aphananthe cuspidata* | 10 | H | 3.5 | 0.1 | 0.71 | 0.08 |
| 27 | *Apodytes dimidiata* | 11 | Q | 0.5 | 0.09 | 0.87 | 0.1 |
| 28 | *Aporosa indo-acuminata* | 17 | LQ | 0.5 | 0.06 | 0.91 | 0.07 |
| 29 | *Aporosa cardiosperma* | 54 | Q | 5 | 0.02 | 0.87 | 0.11 |
| 30 | *Archidendron bigeminum* | 26 | H | 1.5 | 0.04 | 0.78 | 0.11 |
| 31 | *Ardisia missionis* | 5 | QH | 0.5 | 0.2 | 0.95 | 0.05 |
| 32 | *Ardisia pauciflora* | 19 | L | 2.5 | 0.05 | 0.91 | 0.06 |
| 33 | *Ardisia rhomboidea* | 12 | Q | 1.5 | 0.08 | 0.98 | 0.02 |
| 34 | *Ardisia solanacea* | 10 | Q | 3 | 0.1 | 0.76 | 0.06 |
| 35 | *Ardisia stonei* | 5 | H | 2 | 0.2 | 0.94 | 0.04 |
| 36 | *Arenga wightii* | 16 | LQH | 0.5 | 0.06 | 0.93 | 0.06 |
| 37 | *Artocarpus gomezianus* | 22 | Q | 3.5 | 0.25 | 0.62 | 0.21 |
| 38 | *Artocarpus heterophyllus* | 60 | QH | 2.5 | 0.02 | 0.84 | 0.06 |
| 39 | *Artocarpus hirsutus* | 62 | H | 5 | 0.05 | 0.75 | 0.14 |
| 40 | *Atalantia racemosa* | 14 | H | 1.5 | 0.07 | 0.79 | 0.16 |
| 41 | *Atalantia wightii* | 31 | L | 0.5 | 0.04 | 0.78 | 0.06 |
| 42 | *Atuna indica* | 6 | LQH | 2.5 | 0 | 1 | 0.02 |
| 43 | *Atuna travancorica* | 8 | H | 3 | 0 | 0.86 | 0.12 |
| 44 | *Baccaurea courtallensis* | 49 | LQ | 5 | 0 | 0.85 | 0.06 |
| 45 | *Beilschmiedia dalzellii* | 67 | QH | 2.5 | 0.03 | 0.84 | 0.1 |
| 46 | *Beilschmiedia wightii* | 18 | L | 5 | 0.06 | 0.91 | 0.07 |
| 47 | *Bhesa indica* | 5 | Q | 0.5 | 0.2 | 0.97 | 0.02 |

| | | | | | | | |
|---|---|---|---|---|---|---|---|
| 48 | *Bischofia javanica* | 31 | L | 0.5 | 0 | 0.82 | 0.11 |
| 49 | *Blachia denudata* | 15 | H | 2.5 | 0.07 | 0.89 | 0.12 |
| 50 | *Blepharistemma serratum* | 12 | L | 5 | 0.08 | 0.74 | 0.09 |
| 51 | *Bombax ceiba* | 13 | H | 3.5 | 0 | 0.63 | 0.24 |
| 52 | *Callicarpa tomentosa* | 47 | H | 5 | 0.08 | 0.77 | 0.17 |
| 53 | *Calophyllum apetalum* | 21 | Q | 1 | 0.05 | 0.93 | 0.09 |
| 54 | *Calophyllum austroindicum* | 10 | LQH | 0.5 | 0.1 | 0.93 | 0.07 |
| 55 | *Calophyllum tomentosum* | 72 | LQ | 4.5 | 0.01 | 0.84 | 0.02 |
| 56 | *Canarium strictum* | 45 | L | 0.5 | 0.05 | 0.83 | 0.04 |
| 57 | *Capparis baducca* | 8 | LQ | 0.5 | 0.25 | 0.89 | 0.05 |
| 58 | *Carallia brachiata* | 21 | H | 2.5 | 0 | 0.83 | 0.06 |
| 59 | *Caryota urens* | 43 | LQH | 0.5 | 0.02 | 0.82 | 0.08 |
| 60 | *Casearia ovata* | 80 | LQ | 0.5 | 0.03 | 0.8 | 0.04 |
| 61 | *Casearia rubescens* | 20 | LQ | 0.5 | 0.05 | 0.81 | 0.1 |
| 62 | *Casearia wynadensis* | 28 | LQ | 1 | 0.04 | 0.92 | 0.03 |
| 63 | *Celtis philippensis* | 16 | L | 4.5 | 0.06 | 0.71 | 0.11 |
| 64 | *Chassalia curviflora* | 4 | L | 1 | 0.5 | 0.65 | 0.28 |
| 65 | *Chionanthus courtallensis* | 8 | Q | 2 | 0.13 | 0.98 | 0.02 |
| 66 | *Chionanthus linocieroides* | 5 | Q | 0.5 | 0.2 | 0.98 | 0.01 |
| 67 | *Chionanthus mala-elengi* | 22 | L | 5 | 0.04 | 0.63 | 0.06 |
| 68 | *Chionanthus ramiflorus* | 24 | QH | 5 | 0 | 0.9 | 0.03 |
| 69 | *Chrysophyllum flexuosum* | 11 | H | 1 | 0.09 | 0.75 | 0.23 |
| 70 | *Chrysophyllum roxburghii* | 25 | L | 5 | 0.08 | 0.72 | 0.12 |
| 71 | *Chukrasia tabularis* | 7 | L | 0.5 | 0.14 | 0.77 | 0.17 |
| 72 | *Cinnamomum filipedicellatum* | 15 | Q | 0.5 | 0.07 | 0.97 | 0.03 |
| 73 | *Cinnamomum keralaense* | 6 | Q | 0.5 | 0.17 | 0.77 | 0.18 |
| 74 | *Cinnamomum macrocarpum* | 7 | H | 1.5 | 0.14 | 0.87 | 0.15 |
| 75 | *Cinnamomum malabatrum* | 134 | Q | 2.5 | 0.02 | 0.81 | 0.03 |

| | | | | | | | |
|---|---|---|---|---|---|---|---|
| 76 | *Cinnamomum sulphuratum* | 9 | L | 1 | 0.11 | 0.72 | 0.23 |
| 77 | *Cinnamomum verum* | 24 | H | 4 | 0 | 0.86 | 0.04 |
| 78 | *Clausena anisata* | 16 | LQ | 0.5 | 0.06 | 0.8 | 0.21 |
| 79 | *Clausena austroindica* | 4 | L | 0.5 | 0.25 | 0.72 | 0.2 |
| 80 | *Clausena indica* | 22 | L | 3 | 0.04 | 0.77 | 0.06 |
| 81 | *Cleidion javanicum* | 12 | LQ | 1 | 0.08 | 0.86 | 0.06 |
| 82 | *Cleistanthus malabaricus* | 6 | H | 4.5 | 0.17 | 0.6 | 0.12 |
| 83 | *Cleistanthus travancorensis* | 9 | L | 2.5 | 0.11 | 0.77 | 0.14 |
| 84 | *Clerodendrum infortunatum* | 18 | L | 4 | 0.06 | 0.83 | 0.16 |
| 85 | *Croton malabaricus* | 41 | H | 5 | 0 | 0.86 | 0.05 |
| 86 | *Croton zeylanicus* | 10 | Q | 2 | 0.1 | 0.9 | 0.05 |
| 87 | *Cryptocarya anamalayana* | 21 | QH | 2.5 | 0.13 | 0.93 | 0.09 |
| 88 | *Cryptocarya beddomei* | 22 | QH | 1 | 0 | 0.9 | 0.01 |
| 89 | *Cryptocarya lawsonii* | 71 | Q | 2.5 | 0 | 0.89 | 0.03 |
| 90 | *Cryptocarya stocksii* | 11 | LQ | 1 | 0.09 | 0.98 | 0.01 |
| 91 | *Cryptocarya wightiana* | 28 | LQH | 1.5 | 0.25 | 0.76 | 0.19 |
| 92 | *Cyathocalyx zeylanicus* | 15 | H | 1.5 | 0.07 | 0.9 | 0.07 |
| 93 | *Cynometra travancorica* | 7 | LQ | 1 | 0.14 | 0.96 | 0.02 |
| 94 | *Dendrocnide sinuata* | 16 | L | 0.5 | 0.06 | 0.91 | 0.05 |
| 95 | *Dichapetalum gelonioides* | 65 | Q | 1.5 | 0.03 | 0.85 | 0.07 |
| 96 | *Dillenia bracteata* | 13 | L | 5 | 0.08 | 0.69 | 0.08 |
| 97 | *Dimocarpus longan* | 118 | Q | 0.5 | 0 | 0.88 | 0.03 |
| 98 | *Dimorphocalyx beddomei* | 19 | LQ | 2 | 0.05 | 0.91 | 0.05 |
| 99 | *Dimorphocalyx glabellus* | 13 | L | 3 | 0.08 | 0.6 | 0.15 |
| 100 | *Diospyros bourdillonii* | 23 | L | 0.5 | 0 | 0.82 | 0.08 |
| 101 | *Diospyros buxifolia* | 33 | H | 3.5 | 0 | 0.66 | 0.1 |
| 102 | *Diospyros candolleana* | 64 | Q | 0.5 | 0.05 | 0.92 | 0.06 |
| 103 | *Diospyros crumenata* | 9 | LQH | 1 | 0.11 | 0.95 | 0.05 |

| | | | | | | | |
|---|---|---|---|---|---|---|---|
| 104 | *Diospyros ebenum* | 25 | LQ | 1 | 0 | 0.83 | 0.02 |
| 105 | *Diospyros ghatensis* | 26 | QH | 4.5 | 0.07 | 0.85 | 0.11 |
| 106 | *Diospyros malabarica* | 6 | H | 3 | 0.17 | 0.65 | 0.21 |
| 107 | *Diospyros montana* | 35 | Q | 2 | 0 | 0.83 | 0.06 |
| 108 | *Diospyros nilagirica* | 11 | Q | 4.5 | 0.09 | 0.6 | 0.03 |
| 109 | *Diospyros oocarpa* | 34 | H | 5 | 0.13 | 0.74 | 0.17 |
| 110 | *Diospyros paniculata* | 49 | L | 4 | 0.21 | 0.73 | 0.2 |
| 111 | *Diospyros pruriens* | 17 | LQ | 1 | 0.06 | 0.96 | 0.02 |
| 112 | *Diospyros ridleyi* | 6 | H | 3 | 0.17 | 0.97 | 0.01 |
| 113 | *Diospyros saldanhae* | 41 | LQH | 1 | 0.03 | 0.88 | 0.06 |
| 114 | *Diospyros sylvatica* | 58 | L | 4.5 | 0 | 0.74 | 0.02 |
| 115 | *Diospyros vera* | 13 | LQH | 3 | 0.08 | 0.93 | 0.03 |
| 116 | *Dipterocarpus bourdillonii* | 8 | LQ | 2.5 | 0.13 | 0.76 | 0.15 |
| 117 | *Dipterocarpus indicus* | 55 | Q | 1 | 0 | 0.9 | 0.08 |
| 118 | *Discospermum apiocarpum* | 5 | Q | 1 | 0.2 | 0.75 | 0.31 |
| 119 | *Discospermum sphaerocarpum* | 15 | Q | 3.5 | 0.07 | 0.92 | 0.07 |
| 120 | *Dracaena elliptica* | 5 | H | 0.5 | 0.2 | 0.91 | 0.08 |
| 121 | *Drypetes confertiflora* | 24 | QH | 4 | 0 | 0.73 | 0.05 |
| 122 | *Drypetes malabarica* | 18 | LQH | 0.5 | 0.06 | 0.89 | 0.07 |
| 123 | *Drypetes oblongifolia* | 24 | H | 3 | 0 | 0.89 | 0.04 |
| 124 | *Drypetes venusta* | 44 | LQ | 3 | 0 | 0.86 | 0.04 |
| 125 | *Drypetes wightii* | 13 | H | 1 | 0.08 | 0.91 | 0.07 |
| 126 | *Durio exarillatus* | 40 | L | 0.5 | 0 | 0.9 | 0.04 |
| 127 | *Dysoxylum gotadhora* | 19 | Q | 3.5 | 0.05 | 0.74 | 0.12 |
| 128 | *Dysoxylum malabaricum* | 54 | Q | 0.5 | 0.02 | 0.77 | 0.11 |
| 129 | *Elaeocarpus munroii* | 12 | QH | 1 | 0.08 | 0.87 | 0.12 |
| 130 | *Elaeocarpus serratus* | 35 | LQ | 0.5 | 0.08 | 0.83 | 0.1 |
| 131 | *Elaeocarpus tuberculatus* | 35 | QH | 2 | 0.08 | 0.88 | 0.11 |

| | | | | | | | |
|---|---|---|---|---|---|---|---|
| 132 | *Elaeocarpus variabilis* | 6 | Q | 1.5 | 0.17 | 0.9 | 0.06 |
| 133 | *Elaeocarpus venustus* | 6 | H | 4.5 | 0.17 | 0.93 | 0.03 |
| 134 | *Epiprinus mallotiformis* | 19 | L | 0.5 | 0.05 | 0.84 | 0.13 |
| 135 | *Erythroxylum moonii* | 8 | LQ | 0.5 | 0.13 | 0.91 | 0.09 |
| 136 | *Eugenia codyensis* | 6 | LQ | 1.5 | 0.17 | 0.94 | 0.07 |
| 137 | *Eugenia kalamii* | 11 | H | 1 | 0.09 | 0.86 | 0.13 |
| 138 | *Eugenia macrocalyx* | 25 | LQ | 0.5 | 0.08 | 0.97 | 0.03 |
| 139 | *Eugenia mooniana* | 24 | L | 5 | 0 | 0.89 | 0.07 |
| 140 | *Euonymus angulatus* | 5 | L | 3 | 0 | 0.95 | 0.02 |
| 141 | *Euonymus dichotomus* | 7 | Q | 2.5 | 0.14 | 0.9 | 0.12 |
| 142 | *Euonymus indicus* | 26 | Q | 1 | 0.13 | 0.83 | 0.11 |
| 143 | *Eurya nitida* | 7 | LQ | 2 | 0.14 | 0.97 | 0.01 |
| 144 | *Excoecaria oppositifolia* | 7 | H | 0.5 | 0.14 | 0.96 | 0.03 |
| 145 | *Falconeria insignis* | 6 | H | 4 | 0.17 | 0.74 | 0.07 |
| 146 | *Ficus beddomei* | 15 | L | 0.5 | 0.07 | 0.83 | 0.13 |
| 147 | *Ficus callosa* | 17 | L | 1 | 0.06 | 0.65 | 0.18 |
| 148 | *Ficus exasperata* | 5 | LQH | 0.5 | 0.2 | 0.87 | 0.14 |
| 149 | *Ficus microcarpa* | 7 | H | 2 | 0.29 | 0.66 | 0.23 |
| 150 | *Ficus nervosa* | 52 | Q | 3.5 | 0.02 | 0.8 | 0.1 |
| 151 | *Ficus talbotii* | 4 | L | 1.5 | 0.25 | 0.73 | 0.1 |
| 152 | *Ficus virens* | 16 | Q | 1 | 0.06 | 0.76 | 0.16 |
| 153 | *Filicium decipiens* | 7 | LQH | 3 | 0.14 | 0.94 | 0.07 |
| 154 | *Flacourtia montana* | 79 | H | 2 | 0.04 | 0.79 | 0.16 |
| 155 | *Garcinia gummi-gutta* | 68 | Q | 4 | 0 | 0.86 | 0.02 |
| 156 | *Garcinia indica* | 33 | H | 5 | 0.16 | 0.78 | 0.18 |
| 157 | *Garcinia morella* | 77 | Q | 1 | 0.03 | 0.91 | 0.04 |
| 158 | *Garcinia rubroechinata* | 5 | H | 5 | 0 | 0.93 | 0.02 |
| 159 | *Garcinia talbotii* | 39 | LQ | 1 | 0.03 | 0.79 | 0.1 |

| | | | | | | | |
|---|---|---|---|---|---|---|---|
| 160 | *Glochidion ellipticum* | 74 | H | 5 | 0.04 | 0.73 | 0.11 |
| 161 | *Gluta travancorica* | 13 | Q | 1 | 0.08 | 0.98 | 0.02 |
| 162 | *Glycosmis macrocarpa* | 19 | H | 1 | 0.05 | 0.85 | 0.1 |
| 163 | *Glycosmis pentaphylla* | 36 | H | 1 | 0.08 | 0.89 | 0.06 |
| 164 | *Gomphandra coriacea* | 19 | QH | 1 | 0.05 | 0.94 | 0.06 |
| 165 | *Gomphandra tetrandra* | 45 | LQ | 0.5 | 0 | 0.89 | 0.04 |
| 166 | *Goniothalamus cardiopetalus* | 19 | H | 3 | 0.05 | 0.86 | 0.08 |
| 167 | *Goniothalamus rhynchantherus* | 4 | H | 4.5 | 0.25 | 0.99 | 0.01 |
| 168 | *Goniothalamus wightii* | 8 | QH | 1.5 | 0.13 | 0.96 | 0.05 |
| 169 | *Gordonia obtusa* | 18 | QH | 1 | 0.06 | 0.88 | 0.13 |
| 170 | *Gymnacranthera canarica* | 10 | Q | 5 | 0.1 | 0.6 | 0.11 |
| 171 | *Helicia nilagirica* | 4 | Q | 3 | 0.25 | 0.65 | 0.06 |
| 172 | *Heritiera papilio* | 25 | QH | 1 | 0.08 | 0.86 | 0.12 |
| 173 | *Heynea trijuga* | 22 | H | 1.5 | 0.05 | 0.85 | 0.06 |
| 174 | *Holigarna arnottiana* | 73 | QH | 0.5 | 0.08 | 0.86 | 0.12 |
| 175 | *Holigarna ferruginea* | 25 | H | 3 | 0 | 0.94 | 0.04 |
| 176 | *Holigarna grahamii* | 79 | QH | 4.5 | 0.01 | 0.88 | 0.07 |
| 177 | *Holigarna nigra* | 38 | Q | 3.5 | 0.06 | 0.84 | 0.08 |
| 178 | *Homalium ceylanicum* | 18 | Q | 1.5 | 0.06 | 0.85 | 0.14 |
| 179 | *Hopea canarensis* | 15 | LQH | 1.5 | 0.07 | 0.98 | 0.02 |
| 180 | *Hopea erosa* | 6 | H | 5 | 0.17 | 0.87 | 0.09 |
| 181 | *Hopea glabra* | 7 | H | 5 | 0 | 0.77 | 0.11 |
| 182 | *Hopea parviflora* | 39 | H | 1.5 | 0.03 | 0.8 | 0.08 |
| 183 | *Hopea ponga* | 42 | Q | 1.5 | 0.08 | 0.92 | 0.09 |
| 184 | *Hopea racophloea* | 4 | Q | 0.5 | 0.25 | 0.8 | 0.2 |
| 185 | *Humboldtia brunonis* | 19 | QH | 1 | 0.05 | 0.97 | 0.02 |
| 186 | *Humboldtia decurrens* | 4 | LQH | 1 | 0.25 | 0.93 | 0.06 |
| 187 | *Humboldtia vahliana* | 6 | Q | 3 | 0.17 | 0.63 | 0.05 |

| | | | | | | | |
|---|---|---|---|---|---|---|---|
| 188 | *Hunteria zeylanica* | 12 | QH | 0.5 | 0.08 | 0.97 | 0.02 |
| 189 | *Hydnocarpus alpina* | 18 | QH | 1 | 0.06 | 0.87 | 0.12 |
| 190 | *Hydnocarpus macrocarpa* | 5 | LQH | 0.5 | 0.2 | 0.89 | 0.11 |
| 191 | *Hydnocarpus pentandrus* | 73 | QH | 3.5 | 0 | 0.82 | 0.05 |
| 192 | *Isonandra lanceolata* | 30 | QH | 4.5 | 0 | 0.88 | 0.04 |
| 193 | *Ixora alba* | 6 | H | 0.5 | 0.33 | 0.79 | 0.17 |
| 194 | *Ixora brachiata* | 87 | Q | 3 | 0 | 0.84 | 0.09 |
| 195 | *Ixora elongata* | 23 | LQH | 1.5 | 0 | 0.96 | 0.03 |
| 196 | *Ixora lanceolaria* | 6 | L | 2 | 0.17 | 0.63 | 0.11 |
| 197 | *Ixora nigricans* | 50 | Q | 5 | 0.06 | 0.81 | 0.09 |
| 198 | *Kingiodendron pinnatum* | 19 | LQH | 1 | 0.05 | 0.94 | 0.04 |
| 199 | *Knema attenuata* | 127 | LQ | 2 | 0.01 | 0.83 | 0.05 |
| 200 | *Lagerstroemia microcarpa* | 49 | Q | 5 | 0.06 | 0.81 | 0.13 |
| 201 | *Lasianthus acuminatus* | 17 | H | 1 | 0.06 | 0.95 | 0.05 |
| 202 | *Lasianthus jackianus* | 13 | H | 3.5 | 0.08 | 0.96 | 0.04 |
| 203 | *Lasianthus rostratus* | 4 | QH | 0.5 | 0.25 | 1 | 0 |
| 204 | *Leea indica* | 67 | H | 5 | 0.01 | 0.79 | 0.09 |
| 205 | *Lepisanthes erecta* | 8 | QH | 3 | 0.13 | 0.93 | 0.04 |
| 206 | *Lepisanthes tetraphylla* | 58 | L | 4 | 0.02 | 0.81 | 0.07 |
| 207 | *Leptonychia caudata* | 8 | H | 1.5 | 0.13 | 0.84 | 0.12 |
| 208 | *Ligustrum perrottetii* | 4 | H | 0.5 | 0.25 | 0.93 | 0.08 |
| 209 | *Litsea floribunda* | 73 | Q | 0.5 | 0.01 | 0.92 | 0.04 |
| 210 | *Litsea ghatica* | 8 | LQH | 3 | 0 | 0.96 | 0.02 |
| 211 | *Litsea glabrata* | 16 | L | 1.5 | 0.06 | 0.94 | 0.03 |
| 212 | *Litsea keralana* | 23 | L | 0.5 | 0.1 | 0.89 | 0.1 |
| 213 | *Litsea laevigata* | 61 | H | 5 | 0.03 | 0.78 | 0.16 |
| 214 | *Litsea mysorensis* | 36 | LQH | 1.5 | 0.03 | 0.88 | 0.05 |
| 215 | *Litsea oleoides* | 43 | LQ | 1.5 | 0.05 | 0.86 | 0.06 |

| | | | | | | | |
|---|---|---|---|---|---|---|---|
| 216 | *Litsea stocksii* | 17 | H | 3.5 | 0.06 | 0.89 | 0.08 |
| 217 | *Litsea travancorica* | 9 | L | 1 | 0.11 | 0.97 | 0.01 |
| 218 | *Litsea wightiana* | 9 | QH | 3 | 0.11 | 0.92 | 0.11 |
| 219 | *Lophopetalum wightianum* | 32 | L | 0.5 | 0 | 0.85 | 0.09 |
| 220 | *Macaranga indica* | 11 | L | 1 | 0.09 | 0.91 | 0.05 |
| 221 | *Macaranga peltata* | 76 | QH | 2 | 0.04 | 0.8 | 0.07 |
| 222 | *Madhuca bourdillonii* | 7 | L | 0.5 | 0.29 | 0.67 | 0.19 |
| 223 | *Madhuca neriifolia* | 10 | Q | 0.5 | 0.1 | 0.89 | 0.1 |
| 224 | *Maesa indica* | 7 | L | 0.5 | 0.14 | 0.86 | 0.12 |
| 225 | *Magnolia nilagirica* | 7 | L | 5 | 0.14 | 0.69 | 0.02 |
| 226 | *Mallotus aureopunctatus* | 13 | LQ | 2 | 0.08 | 0.77 | 0.13 |
| 227 | *Mallotus beddomei* | 11 | H | 1 | 0.09 | 0.92 | 0.12 |
| 228 | *Mallotus nudiflorus* | 10 | Q | 0.5 | 0.1 | 0.76 | 0.18 |
| 229 | *Mallotus philippensis* | 73 | QH | 0.5 | 0.03 | 0.79 | 0.09 |
| 230 | *Mallotus resinosus* | 16 | LQ | 0.5 | 0.06 | 0.92 | 0.06 |
| 231 | *Mallotus rhamnifolius* | 4 | H | 1.5 | 0.25 | 0.89 | 0.16 |
| 232 | *Mammea suriga* | 11 | H | 4.5 | 0.09 | 0.8 | 0.13 |
| 233 | *Mangifera indica* | 93 | LQH | 1 | 0.01 | 0.81 | 0.05 |
| 234 | *Margaritaria indica* | 13 | QH | 3 | 0.08 | 0.73 | 0.09 |
| 235 | *Mastixia arborea* | 47 | LQ | 1.5 | 0 | 0.91 | 0.04 |
| 236 | *Maytenus rothiana* | 13 | L | 4.5 | 0.08 | 0.75 | 0.06 |
| 237 | *Meiogyne pannosa* | 47 | Q | 2 | 0.02 | 0.89 | 0.06 |
| 238 | *Meiogyne ramarowii* | 40 | H | 5 | 0 | 0.86 | 0.04 |
| 239 | *Melicope lunu-ankenda* | 26 | Q | 0.5 | 0 | 0.8 | 0.07 |
| 240 | *Meliosma pinnata* | 6 | H | 2 | 0.17 | 0.93 | 0.03 |
| 241 | *Memecylon gracile* | 16 | LQ | 2 | 0.06 | 0.83 | 0.09 |
| 242 | *Memecylon heyneanum* | 18 | QH | 2.5 | 0.06 | 0.86 | 0.06 |
| 243 | *Memecylon malabaricum* | 13 | LQ | 0.5 | 0.08 | 0.87 | 0.11 |

| | | | | | | | |
|---|---|---|---|---|---|---|---|
| 244 | *Memecylon pseudogracile* | 11 | H | 0.5 | 0.09 | 0.99 | 0.01 |
| 245 | *Memecylon randerianum* | 12 | LQ | 0.5 | 0.08 | 0.84 | 0.14 |
| 246 | *Memecylon subsessile* | 7 | L | 0.5 | 0.14 | 0.95 | 0.08 |
| 247 | *Memecylon talbotianum* | 33 | Q | 4.5 | 0 | 0.78 | 0.1 |
| 248 | *Memecylon terminale* | 7 | H | 0.5 | 0.14 | 0.98 | 0.02 |
| 249 | *Memecylon umbellatum* | 32 | L | 5 | 0.03 | 0.77 | 0.11 |
| 250 | *Memecylon wightii* | 27 | LQ | 1 | 0 | 0.91 | 0.02 |
| 251 | *Mesua ferrea* | 59 | LQH | 4.5 | 0 | 0.91 | 0.01 |
| 252 | *Meteoromyrtus wynaadensis* | 6 | L | 0.5 | 0.33 | 0.75 | 0.18 |
| 253 | *Meyna laxiflora* | 15 | L | 4.5 | 0.07 | 0.87 | 0.08 |
| 254 | *Microtropis latifolia* | 35 | QH | 5 | 0.03 | 0.9 | 0.01 |
| 255 | *Miliusa gokhalaei* | 6 | LQH | 2.5 | 0.17 | 0.96 | 0.03 |
| 256 | *Miliusa nilagirica* | 11 | LQ | 2.5 | 0.09 | 0.91 | 0.03 |
| 257 | *Mimusops elengi* | 51 | QH | 2.5 | 0.02 | 0.86 | 0.1 |
| 258 | *Mitragyna parvifolia* | 7 | QH | 2 | 0.14 | 0.68 | 0.22 |
| 259 | *Mitragyna tubulosa* | 4 | LQ | 0.5 | 0.25 | 0.93 | 0.06 |
| 260 | *Mitrephora grandiflora* | 8 | Q | 1.5 | 0.13 | 0.76 | 0.27 |
| 261 | *Murraya koenigii* | 16 | H | 1.5 | 0.06 | 0.84 | 0.11 |
| 262 | *Murraya paniculata* | 9 | LQH | 1 | 0.11 | 0.83 | 0.11 |
| 263 | *Myristica beddomei* | 112 | Q | 5 | 0.02 | 0.86 | 0.02 |
| 264 | *Myristica malabarica* | 58 | Q | 3.5 | 0.02 | 0.68 | 0.11 |
| 265 | *Neolitsea fischeri* | 13 | LQ | 0.5 | 0.08 | 0.92 | 0.06 |
| 266 | *Neolitsea zeylanica* | 13 | QH | 1 | 0.08 | 0.86 | 0.13 |
| 267 | *Nothapodytes nimmoniana* | 27 | Q | 5 | 0.11 | 0.84 | 0.14 |
| 268 | *Nothopegia aureofulva* | 4 | H | 0.5 | 0.25 | 1 | 0 |
| 269 | *Nothopegia beddomei* | 77 | LQH | 3 | 0 | 0.88 | 0.06 |
| 270 | *Nothopegia racemosa* | 72 | H | 4 | 0 | 0.84 | 0.02 |
| 271 | *Nothopegia travancorica* | 27 | LQ | 2 | 0.11 | 0.74 | 0.14 |

| | | | | | | | |
|---|---|---|---|---|---|---|---|
| 272 | *Ocotea lancifolia* | 27 | L | 2.5 | 0 | 0.94 | 0.06 |
| 273 | *Octotropis travancorica* | 5 | LQH | 0.5 | 0.2 | 0.99 | 0.01 |
| 274 | *Olea dioica* | 97 | QH | 5 | 0.02 | 0.84 | 0.09 |
| 275 | *Oreocnide integrifolia* | 14 | QH | 3 | 0.07 | 0.94 | 0.03 |
| 276 | *Ormosia travancorica* | 16 | L | 0.5 | 0.06 | 0.83 | 0.13 |
| 277 | *Orophea erythrocarpa* | 9 | LQ | 0.5 | 0.11 | 0.89 | 0.08 |
| 278 | *Orophea sivarajanii* | 6 | Q | 2.5 | 0 | 0.94 | 0.01 |
| 279 | *Orophea thomsonii* | 12 | QH | 4.5 | 0.08 | 0.93 | 0.02 |
| 280 | *Otonephelium stipulaceum* | 42 | Q | 1 | 0 | 0.9 | 0.07 |
| 281 | *Pajanelia longifolia* | 9 | LQ | 0.5 | 0.11 | 0.93 | 0.05 |
| 282 | *Palaquium bourdillonii* | 6 | L | 0.5 | 0.17 | 0.97 | 0.03 |
| 283 | *Palaquium ellipticum* | 91 | LQH | 3.5 | 0.04 | 0.86 | 0.04 |
| 284 | *Pandanus furcatus* | 23 | QH | 1 | 0.04 | 0.88 | 0.05 |
| 285 | *Paracroton pendulus subsp. zeylanicus* | 46 | LQ | 0.5 | 0 | 0.83 | 0.05 |
| 286 | *Pavetta sp* | 8 | H | 5 | 0.13 | 0.9 | 0.05 |
| 287 | *Persea macrantha* | 99 | QH | 2 | 0 | 0.88 | 0.05 |
| 288 | *Pinanga dicksonii* | 20 | Q | 1 | 0.05 | 0.92 | 0.09 |
| 289 | *Poeciloneuron indicum* | 33 | QH | 0.5 | 0.16 | 0.96 | 0.05 |
| 290 | *Polyalthia coffeoides* | 21 | LQ | 1 | 0 | 0.91 | 0.02 |
| 291 | *Polyalthia fragrans* | 57 | LQ | 1 | 0 | 0.86 | 0.06 |
| 292 | *Polyalthia malabarica* | 13 | H | 0.5 | 0.08 | 0.95 | 0.03 |
| 293 | *Prismatomeris tetrandra* | 5 | L | 5 | 0 | 0.77 | 0.03 |
| 294 | *Prunus ceylanica* | 31 | H | 4 | 0 | 0.84 | 0.06 |
| 295 | *Psychotria anamallayana* | 8 | LQ | 1 | 0.13 | 0.95 | 0.04 |
| 296 | *Psychotria dalzellii* | 23 | H | 4.5 | 0.04 | 0.75 | 0.07 |
| 297 | *Psychotria flavida* | 7 | L | 1 | 0.14 | 0.65 | 0.21 |
| 298 | *Psychotria nigra* | 73 | Q | 1.5 | 0.01 | 0.88 | 0.05 |
| 299 | *Psychotria truncata* | 16 | LQH | 0.5 | 0.06 | 0.9 | 0.11 |

| | | | | | | | |
|---|---|---|---|---|---|---|---|
| 300 | *Psydrax dicoccos* | 39 | H | 4 | 0.05 | 0.79 | 0.14 |
| 301 | *Pterospermum diversifolium* | 16 | QH | 2 | 0.06 | 0.81 | 0.15 |
| 302 | *Pterospermum reticulatum* | 22 | Q | 0.5 | 0.15 | 0.71 | 0.08 |
| 303 | *Pterospermum rubiginosum* | 8 | L | 3.5 | 0.13 | 0.77 | 0.1 |
| 304 | *Pterygota alata* | 17 | LQH | 1.5 | 0.06 | 0.92 | 0.06 |
| 305 | *Rapanea wightiana* | 8 | L | 0.5 | 0.13 | 0.91 | 0.04 |
| 306 | *Reinwardtiodendron anamalaiense* | 72 | LQH | 4 | 0.03 | 0.84 | 0.04 |
| 307 | *Sageraea laurina* | 5 | L | 0.5 | 0.2 | 0.97 | 0.03 |
| 308 | *Sageraea thwaitesii* | 16 | H | 1 | 0.06 | 0.94 | 0.05 |
| 309 | *Saprosma corymbosum* | 7 | QH | 0.5 | 0.14 | 0.99 | 0.01 |
| 310 | *Saprosma glomerata* | 14 | LQ | 2 | 0.07 | 0.94 | 0.02 |
| 311 | *Schefflera micrantha* | 7 | LQ | 0.5 | 0.14 | 0.91 | 0.04 |
| 312 | *Scolopia crenata* | 22 | LQ | 3 | 0.04 | 0.82 | 0.1 |
| 313 | *Semecarpus auriculata* | 15 | LQ | 0.5 | 0.07 | 0.9 | 0.1 |
| 314 | *Semecarpus travancorica* | 7 | H | 3 | 0.14 | 0.94 | 0.03 |
| 315 | *Sterculia guttata* | 19 | LQ | 1.5 | 0.05 | 0.71 | 0.15 |
| 316 | *Stereospermum tetragonum* | 40 | H | 4 | 0.15 | 0.72 | 0.2 |
| 317 | *Strombosia ceylanica* | 44 | H | 5 | 0 | 0.78 | 0.03 |
| 318 | *Symplocos cochinchinensis* | 6 | H | 3 | 0.17 | 0.7 | 0.08 |
| 319 | *Symplocos macrophylla* | 9 | H | 0.5 | 0.11 | 1 | 0.01 |
| 320 | *Symplocos racemosa* | 57 | LQ | 0.5 | 0.05 | 0.91 | 0.02 |
| 321 | *Symplocos rosea* | 15 | H | 1.5 | 0.07 | 0.91 | 0.07 |
| 322 | *Syzygium benthamianum* | 9 | Q | 2.5 | 0.11 | 0.94 | 0.07 |
| 323 | *Syzygium caryophyllatum* | 8 | LQ | 0.5 | 0.13 | 0.96 | 0.02 |
| 324 | *Syzygium cumini* | 75 | QH | 3.5 | 0.05 | 0.79 | 0.09 |
| 325 | *Syzygium densiflorum* | 11 | Q | 0.5 | 0.09 | 0.87 | 0.12 |
| 326 | *Syzygium gardneri* | 105 | LQH | 5 | 0.01 | 0.85 | 0.07 |
| 327 | *Syzygium grande* | 11 | QH | 0.5 | 0.09 | 0.9 | 0.1 |

| | | | | | | | |
|---|---|---|---|---|---|---|---|
| 328 | *Syzygium hemisphericum* | 47 | Q | 2 | 0.02 | 0.83 | 0.03 |
| 329 | *Syzygium laetum* | 78 | LQ | 4 | 0 | 0.89 | 0.03 |
| 330 | *Syzygium lanceolatum* | 10 | Q | 0.5 | 0.1 | 0.96 | 0.03 |
| 331 | *Syzygium mundagam* | 27 | H | 3 | 0.25 | 0.87 | 0.11 |
| 332 | *Syzygium munronii* | 23 | L | 0.5 | 0 | 0.87 | 0.07 |
| 333 | *Syzygium rubicundum* | 6 | Q | 1.5 | 0.17 | 0.7 | 0.14 |
| 334 | *Tabernaemontana alternifolia* | 70 | QH | 5 | 0 | 0.85 | 0.05 |
| 335 | *Tabernaemontana gamblei* | 10 | LQH | 0.5 | 0.1 | 0.95 | 0.04 |
| 336 | *Tarenna nilagirica* | 9 | LQH | 1.5 | 0.11 | 0.87 | 0.09 |
| 337 | *Terminalia bellirica* | 33 | H | 5 | 0.03 | 0.83 | 0.06 |
| 338 | *Terminalia paniculata* | 41 | Q | 5 | 0.1 | 0.83 | 0.11 |
| 339 | *Terminalia travancorensis* | 11 | LQ | 0.5 | 0.09 | 0.9 | 0.07 |
| 340 | *Tetrameles nudiflora* | 10 | H | 4.5 | 0 | 0.72 | 0.25 |
| 341 | *Toona ciliata* | 18 | H | 3.5 | 0.06 | 0.78 | 0.09 |
| 342 | *Turpinia malabarica* | 19 | L | 1 | 0.05 | 0.79 | 0.08 |
| 343 | *Vateria indica* | 50 | LQH | 5 | 0 | 0.82 | 0.09 |
| 344 | *Vepris bilocularis* | 43 | LQ | 2 | 0.02 | 0.82 | 0.06 |
| 345 | *Vitex altissima* | 47 | LQH | 2.5 | 0 | 0.81 | 0.08 |
| 346 | *Walsura trifoliolata* | 15 | Q | 3 | 0.07 | 0.91 | 0.04 |
| 347 | *Xanthophyllum arnottianum* | 53 | L | 0.5 | 0 | 0.86 | 0.07 |
| 348 | *Xantolis tomentosa* | 40 | LQH | 4.5 | 0.05 | 0.86 | 0.08 |

Table S2.2. The permutation importance of predictor variables for each species' distribution model is shown. Variable code: Annual mean temp: annual mean temperature, Annual prec: annual precipitation; Mean temp dryQ: mean temperature of the driest quarter; Mean temp warmQ: mean temperature of the warmest quarter; Prec dryQ: precipitation of the driest quarter; Prec warmQ: precipitation of the warmest quarter.

| Sno. | Species | Annual mean temp. | Annual prec. | Elevation | Mean temp. dryQ | Mean temp warmQ | Prec dryQ | Prec warmQ |
|---|---|---|---|---|---|---|---|---|
| 1 | *Acrocarpus fraxinifolius* | 0 | 38.02 | 0 | 0 | 61.98 | 0 | 0 |
| 2 | *Acronychia pedunculata* | 0 | 0 | 0 | 0 | 0 | 100 | 0 |
| 3 | *Actephila excelsa* | 0 | 14.84 | 7.42 | 0 | 0 | 54.24 | 23.5 |
| 4 | *Actinodaphne bourdillonii* | 0 | 40.24 | 0 | 0 | 57.24 | 2.52 | 0 |
| 5 | *Actinodaphne gullavara* | 1.19 | 0 | 0 | 0 | 0 | 83.57 | 15.24 |
| 6 | *Actinodaphne hookeri* | 0 | 0 | 0 | 0 | 33.73 | 66.27 | 0 |
| 7 | *Actinodaphne tadulingamii* | 0 | 0.83 | 4.79 | 3.54 | 90.74 | 0 | 0.11 |
| 8 | *Actinodaphne wightiana* | 0 | 13.24 | 20.95 | 0 | 53.56 | 6.59 | 5.66 |
| 9 | *Aglaia bourdillonii* | 0 | 0.97 | 0 | 0 | 0 | 95.92 | 3.12 |
| 10 | *Aglaia canarana* | 0 | 11.47 | 17.2 | 0 | 25.03 | 46.3 | 0 |
| 11 | *Aglaia edulis* | 0 | 62.25 | 37.46 | 0 | 0 | 0.01 | 0.28 |
| 12 | *Aglaia elaeagnoidea* | 7.81 | 5.39 | 20.03 | 3.56 | 62.38 | 0.68 | 0.15 |
| 13 | *Aglaia lawii* | 31.45 | 53.55 | 15 | 0 | 0 | 0 | 0 |
| 14 | *Aglaia perviridis* | 0 | 18.81 | 51.09 | 3.59 | 8.73 | 1.8 | 15.98 |
| 15 | *Aglaia simplicifolia* | 0.3 | 0.16 | 8.52 | 20.41 | 69.53 | 0 | 1.08 |
| 16 | *Agrostistachys borneensis* | 0 | 0.74 | 0 | 0 | 75.08 | 24.18 | 0 |
| 17 | *Agrostistachys indica* | 0 | 0 | 2.49 | 7.17 | 61.88 | 18.48 | 9.99 |
| 18 | *Aidia densiflora* | 0 | 0 | 0 | 0 | 0 | 56.41 | 43.59 |
| 19 | *Allophylus cobbe* | 0 | 0 | 31.18 | 0 | 0 | 68.82 | 0 |
| 20 | *Alseodaphne semecarpifolia* | 12.99 | 0 | 0 | 0 | 56.99 | 18.69 | 11.34 |
| 21 | *Alstonia scholaris* | 2.83 | 31.18 | 0 | 0.73 | 0 | 0 | 65.26 |
| 22 | *Anacolosa densiflora* | 0 | 0.43 | 24.09 | 0 | 0 | 74.05 | 1.43 |
| 23 | *Antiaris toxicaria* | 0 | 45.98 | 11.31 | 1.77 | 0 | 40.94 | 0 |

| | | | | | | | | |
|---|---|---|---|---|---|---|---|---|
| 24 | *Antidesma montanum* | 0 | 12.67 | 6.39 | 0 | 70.44 | 10.49 | 0.01 |
| 25 | *Aphanamixis polystachya* | 0 | 4.99 | 0 | 0 | 83.64 | 11.37 | 0 |
| 26 | *Aphananthe cuspidata* | 0 | 0 | 100 | 0 | 0 | 0 | 0 |
| 27 | *Apodytes dimidiata* | 0 | 0.64 | 22.74 | 0 | 73.79 | 0 | 2.83 |
| 28 | *Aporosa indo-acuminata* | 0 | 1.25 | 16.22 | 0 | 45.58 | 28.4 | 8.56 |
| 29 | *Aporosa cardiosperma* | 0 | 1.92 | 31.66 | 0.2 | 47.64 | 0 | 18.59 |
| 30 | *Archidendron bigeminum* | 0 | 37.96 | 40.23 | 0 | 0 | 12.17 | 9.64 |
| 31 | *Ardisia missionis* | 0 | 87.49 | 9.58 | 0 | 0 | 2.74 | 0.2 |
| 32 | *Ardisia pauciflora* | 0 | 0 | 0 | 0 | 88.35 | 11.65 | 0 |
| 33 | *Ardisia rhomboidea* | 0 | 29.66 | 0 | 0 | 24.37 | 45.96 | 0 |
| 34 | *Ardisia solanacea* | 0 | 0 | 0 | 0 | 0 | 75.95 | 24.05 |
| 35 | *Ardisia stonei* | 0 | 1.41 | 44.26 | 0 | 0 | 54.33 | 0 |
| 36 | *Arenga wightii* | 0 | 1.94 | 40.46 | 0 | 42.94 | 8.82 | 5.84 |
| 37 | *Artocarpus gomezianus* | 0 | 2.54 | 22.66 | 0 | 35.76 | 0 | 39.05 |
| 38 | *Artocarpus heterophyllus* | 0 | 7.9 | 14.59 | 0 | 74.54 | 2.96 | 0.02 |
| 39 | *Artocarpus hirsutus* | 0 | 52.98 | 11.25 | 5.43 | 30.34 | 0 | 0 |
| 40 | *Atalantia racemosa* | 0 | 74.35 | 20.03 | 0 | 0 | 5.62 | 0 |
| 41 | *Atalantia wightii* | 0 | 16.88 | 18.43 | 0 | 61.46 | 3.23 | 0 |
| 42 | *Atuna indica* | 0 | 3.13 | 5.26 | 0 | 0 | 91.61 | 0 |
| 43 | *Atuna travancorica* | 0 | 0 | 4.38 | 0 | 0 | 95.62 | 0 |
| 44 | *Baccaurea courtallensis* | 0 | 0 | 30.59 | 0 | 43.32 | 1.2 | 24.89 |
| 45 | *Beilschmiedia dalzellii* | 0 | 5 | 32.71 | 0 | 61.92 | 0.37 | 0 |
| 46 | *Beilschmiedia wightii* | 8.17 | 40.37 | 0 | 0 | 51.46 | 0 | 0 |
| 47 | *Bhesa indica* | 0 | 3.96 | 0.3 | 0 | 95.48 | 0.26 | 0 |
| 48 | *Bischofia javanica* | 0 | 20.4 | 3.21 | 6.35 | 66.73 | 0.93 | 2.37 |
| 49 | *Blachia denudata* | 0 | 75.01 | 5.33 | 0 | 0 | 0 | 19.66 |
| 50 | *Blepharistemma serratum* | 0 | 0 | 0 | 0 | 0 | 0 | 100 |
| 51 | *Bombax ceiba* | 0 | 0 | 0 | 0 | 0 | 100 | 0 |
| 52 | *Callicarpa tomentosa* | 0 | 56.5 | 0 | 0 | 43.17 | 0 | 0.33 |

| | | | | | | | | |
|---|---|---|---|---|---|---|---|---|
| 53 | *Calophyllum apetalum* | 0 | 0.21 | 31.64 | 0 | 57.72 | 0.52 | 9.9 |
| 54 | *Calophyllum austroindicum* | 0.91 | 1.16 | 0.38 | 0 | 0 | 87.69 | 9.85 |
| 55 | *Calophyllum tomentosum* | 0 | 13.89 | 19.32 | 0 | 58.57 | 5.43 | 2.79 |
| 56 | *Canarium strictum* | 0 | 19.94 | 13.68 | 8.71 | 46.18 | 10.85 | 0.64 |
| 57 | *Capparis baducca* | 0 | 2.64 | 62.65 | 0 | 27.61 | 0.88 | 6.22 |
| 58 | *Carallia brachiata* | 34.41 | 61.99 | 3.6 | 0 | 0 | 0 | 0 |
| 59 | *Caryota urens* | 0 | 8.84 | 23.51 | 0.72 | 54.57 | 4.41 | 7.94 |
| 60 | *Casearia ovata* | 0 | 5.37 | 23.07 | 0 | 67.89 | 2.79 | 0.88 |
| 61 | *Casearia rubescens* | 0 | 0.1 | 6.57 | 0 | 84.16 | 2.41 | 6.76 |
| 62 | *Casearia wynadensis* | 0 | 10.65 | 9.6 | 0.06 | 36.31 | 39.81 | 3.56 |
| 63 | *Celtis philippensis* | 0 | 0 | 0 | 0 | 0 | 41.25 | 58.75 |
| 64 | *Chassalia curviflora* | 0 | 0 | 0 | 0 | 27.55 | 59.04 | 13.41 |
| 65 | *Chionanthus courtallensis* | 0 | 1.54 | 0 | 0 | 0 | 98.46 | 0 |
| 66 | *Chionanthus linocieroides* | 0 | 7.26 | 2.24 | 0 | 0 | 88.09 | 2.42 |
| 67 | *Chionanthus mala-elengi* | 0 | 16.17 | 0 | 0 | 83.83 | 0 | 0 |
| 68 | *Chionanthus ramiflorus* | 0 | 30.6 | 21.8 | 0 | 22.54 | 0 | 25.05 |
| 69 | *Chrysophyllum flexuosum* | 0 | 43.26 | 46.53 | 9.92 | 0 | 0 | 0.29 |
| 70 | *Chrysophyllum roxburghii* | 0 | 9.74 | 0 | 0 | 85.01 | 0 | 5.25 |
| 71 | *Chukrasia tabularis* | 0 | 0.74 | 0 | 0 | 37.63 | 61.63 | 0 |
| 72 | *Cinnamomum filipedicellatum* | 0 | 0 | 14.73 | 0 | 71.88 | 13.39 | 0.01 |
| 73 | *Cinnamomum keralaense* | 0 | 0 | 0 | 0 | 98.3 | 1.7 | 0 |
| 74 | *Cinnamomum macrocarpum* | 0 | 0 | 0.92 | 0 | 0 | 99.08 | 0 |
| 75 | *Cinnamomum malabatrum* | 0 | 12 | 19.33 | 0 | 62.9 | 5.77 | 0 |
| 76 | *Cinnamomum sulphuratum* | 0 | 0 | 0 | 0 | 0 | 100 | 0 |
| 77 | *Cinnamomum verum* | 0.76 | 50.75 | 0 | 0 | 0 | 40.69 | 7.8 |
| 78 | *Clausena anisata* | 0 | 6.67 | 25.92 | 0 | 61.97 | 1.58 | 3.87 |
| 79 | *Clausena austroindica* | 0 | 0 | 0 | 0 | 8.2 | 52.63 | 39.17 |
| 80 | *Clausena indica* | 0 | 19.92 | 0 | 0 | 80.08 | 0 | 0 |
| 81 | *Cleidion javanicum* | 0 | 0 | 6.17 | 0 | 69.04 | 24.79 | 0 |

| | | | | | | | | |
|---|---|---|---|---|---|---|---|---|
| 82 | *Cleistanthus malabaricus* | 0 | 100 | 0 | 0 | 0 | 0 | 0 |
| 83 | *Cleistanthus travancorensis* | 0 | 49.74 | 0 | 50.26 | 0 | 0 | 0 |
| 84 | *Clerodendrum infortunatum* | 0 | 0 | 0 | 0 | 100 | 0 | 0 |
| 85 | *Croton malabaricus* | 0 | 6.03 | 21.65 | 1.59 | 0 | 65.95 | 4.78 |
| 86 | *Croton zeylanicus* | 0 | 0 | 1.03 | 0 | 97.11 | 1.86 | 0 |
| 87 | *Cryptocarya anamalayana* | 0 | 9.65 | 0 | 0 | 29.9 | 60.45 | 0 |
| 88 | *Cryptocarya beddomei* | 0 | 24.93 | 23.62 | 0 | 3.56 | 30.24 | 17.65 |
| 89 | *Cryptocarya lawsonii* | 0 | 6.43 | 9.25 | 0 | 81.73 | 0.6 | 2 |
| 90 | *Cryptocarya stocksii* | 0 | 4.47 | 6.18 | 0 | 87.19 | 0 | 2.17 |
| 91 | *Cryptocarya wightiana* | 0 | 59.71 | 4.41 | 0 | 0 | 20.66 | 15.22 |
| 92 | *Cyathocalyx zeylanicus* | 0 | 11.78 | 38.77 | 0.19 | 0 | 48.29 | 0.97 |
| 93 | *Cynometra travancorica* | 0.4 | 0 | 39.3 | 0 | 43.9 | 0 | 16.4 |
| 94 | *Dendrocnide sinuata* | 0 | 1.49 | 0.91 | 2.65 | 94.95 | 0 | 0 |
| 95 | *Dichapetalum gelonioides* | 2.03 | 14.76 | 17.67 | 0.02 | 62.32 | 3.21 | 0 |
| 96 | *Dillenia bracteata* | 0 | 0 | 0 | 0 | 47.61 | 0 | 52.39 |
| 97 | *Dimocarpus longan* | 14.49 | 5.49 | 16.97 | 0 | 62.66 | 0.35 | 0.04 |
| 98 | *Dimorphocalyx beddomei* | 0 | 0 | 20.6 | 0 | 77.58 | 1.82 | 0 |
| 99 | *Dimorphocalyx glabellus* | 0 | 100 | 0 | 0 | 0 | 0 | 0 |
| 100 | *Diospyros bourdillonii* | 0 | 2.49 | 18.39 | 0 | 55.17 | 22.44 | 1.53 |
| 101 | *Diospyros buxifolia* | 7.98 | 33.55 | 0 | 1.04 | 32.68 | 7.24 | 17.51 |
| 102 | *Diospyros candolleana* | 16.49 | 5.74 | 17.45 | 4.39 | 45.88 | 0 | 10.06 |
| 103 | *Diospyros crumenata* | 0 | 8.12 | 20.61 | 0 | 0 | 70 | 1.28 |
| 104 | *Diospyros ebenum* | 0 | 13.08 | 9.7 | 0 | 41.04 | 33.69 | 2.5 |
| 105 | *Diospyros ghatensis* | 0 | 17.18 | 34.76 | 0 | 0 | 33.3 | 14.76 |
| 106 | *Diospyros malabarica* | 0 | 0 | 100 | 0 | 0 | 0 | 0 |
| 107 | *Diospyros montana* | 0 | 2.84 | 26.59 | 0 | 46.6 | 4.37 | 19.6 |
| 108 | *Diospyros nilagirica* | 0 | 0 | 0 | 0 | 0 | 0 | 100 |
| 109 | *Diospyros oocarpa* | 0 | 89.25 | 10.75 | 0 | 0 | 0 | 0 |
| 110 | *Diospyros paniculata* | 0 | 73.42 | 0 | 0 | 2.37 | 21.38 | 2.83 |

| | | | | | | | | |
|---|---|---|---|---|---|---|---|---|
| 111 | *Diospyros pruriens* | 0 | 8.45 | 6.78 | 0 | 71.38 | 13.35 | 0.05 |
| 112 | *Diospyros ridleyi* | 0 | 89.12 | 10.88 | 0 | 0 | 0 | 0 |
| 113 | *Diospyros saldanhae* | 0 | 25.05 | 15.67 | 0 | 41.06 | 5.85 | 12.36 |
| 114 | *Diospyros sylvatica* | 54.35 | 30.93 | 0 | 0 | 4.66 | 10.06 | 0 |
| 115 | *Diospyros vera* | 0 | 40.06 | 3.94 | 0 | 13.86 | 33.19 | 8.94 |
| 116 | *Dipterocarpus bourdillonii* | 0 | 0 | 0 | 0 | 0 | 100 | 0 |
| 117 | *Dipterocarpus indicus* | 0 | 16.1 | 29.45 | 5.7 | 47.92 | 0.65 | 0.19 |
| 118 | *Discospermum apiocarpum* | 0 | 89.99 | 0 | 0 | 0 | 0 | 10.01 |
| 119 | *Discospermum sphaerocarpum* | 0 | 0 | 0 | 0 | 91.78 | 8.22 | 0 |
| 120 | *Dracaena elliptica* | 0 | 0.91 | 10.53 | 3.62 | 0 | 84.94 | 0 |
| 121 | *Drypetes confertiflora* | 0 | 0 | 34.76 | 0 | 30.3 | 0 | 34.94 |
| 122 | *Drypetes malabarica* | 0 | 14.06 | 10.24 | 1.32 | 0 | 51.39 | 23 |
| 123 | *Drypetes oblongifolia* | 0 | 4.2 | 30.63 | 0.16 | 1.5 | 61.59 | 1.93 |
| 124 | *Drypetes venusta* | 0 | 3.97 | 19.4 | 0 | 59.87 | 0.74 | 16.01 |
| 125 | *Drypetes wightii* | 9.58 | 29.32 | 0.89 | 0 | 0 | 60.21 | 0 |
| 126 | *Durio exarillatus* | 0 | 0.03 | 9.96 | 0 | 66.7 | 22.83 | 0.48 |
| 127 | *Dysoxylum gotadhora* | 63.41 | 10.99 | 1.8 | 23.8 | 0 | 0 | 0 |
| 128 | *Dysoxylum malabaricum* | 0 | 0.8 | 29.33 | 3.49 | 63.49 | 2.36 | 0.53 |
| 129 | *Elaeocarpus munroii* | 0 | 15.12 | 52.7 | 0.59 | 0 | 29.47 | 2.12 |
| 130 | *Elaeocarpus serratus* | 0 | 14.34 | 28.9 | 0 | 49.14 | 3.61 | 4.02 |
| 131 | *Elaeocarpus tuberculatus* | 0 | 26.98 | 8.15 | 0 | 51.81 | 9.1 | 3.96 |
| 132 | *Elaeocarpus variabilis* | 0 | 0 | 0 | 0 | 100 | 0 | 0 |
| 133 | *Elaeocarpus venustus* | 0 | 0 | 0 | 0 | 0 | 100 | 0 |
| 134 | *Epiprinus mallotiformis* | 0 | 4.85 | 0 | 0 | 33.35 | 55.15 | 6.64 |
| 135 | *Erythroxylum moonii* | 0 | 0 | 0 | 0 | 6.75 | 69.69 | 23.57 |
| 136 | *Eugenia codyensis* | 0 | 1.44 | 0 | 0 | 67.03 | 30.47 | 1.05 |
| 137 | *Eugenia kalamii* | 0 | 21.52 | 16.55 | 0 | 0 | 25.88 | 36.05 |
| 138 | *Eugenia macrocalyx* | 0 | 14.01 | 11.49 | 0 | 24.07 | 50.12 | 0.31 |
| 139 | *Eugenia mooniana* | 0 | 0 | 0 | 0 | 75.2 | 24.8 | 0 |

| | | | | | | | | |
|---|---|---|---|---|---|---|---|---|
| 140 | *Euonymus angulatus* | 0 | 0 | 0 | 0 | 99.38 | 0.62 | 0 |
| 141 | *Euonymus dichotomus* | 0 | 17.2 | 0 | 0 | 0 | 82.8 | 0 |
| 142 | *Euonymus indicus* | 0 | 11.96 | 21.04 | 0 | 63.47 | 3.53 | 0 |
| 143 | *Eurya nitida* | 0 | 27.11 | 0 | 0 | 72.89 | 0 | 0 |
| 144 | *Excoecaria oppositifolia* | 0 | 1.35 | 35.76 | 8.33 | 0.22 | 51.2 | 3.14 |
| 145 | *Falconeria insignis* | 0 | 0 | 0 | 0 | 0 | 100 | 0 |
| 146 | *Ficus beddomei* | 0 | 33.4 | 0 | 0 | 66.42 | 0 | 0.17 |
| 147 | *Ficus callosa* | 0 | 37.72 | 0 | 0 | 43.12 | 0 | 19.16 |
| 148 | *Ficus exasperata* | 0 | 16.66 | 0 | 0 | 0 | 18.35 | 64.99 |
| 149 | *Ficus microcarpa* | 0 | 0 | 32.01 | 0 | 0 | 67.99 | 0 |
| 150 | *Ficus nervosa* | 0 | 10.52 | 35.29 | 0 | 54.18 | 0 | 0 |
| 151 | *Ficus talbotii* | 100 | 0 | 0 | 0 | 0 | 0 | 0 |
| 152 | *Ficus virens* | 0 | 10.24 | 0 | 0 | 77.64 | 12.12 | 0 |
| 153 | *Filicium decipiens* | 0 | 0 | 0.95 | 0 | 0 | 99.05 | 0 |
| 154 | *Flacourtia montana* | 0 | 33.25 | 0.5 | 3.61 | 34.97 | 13.51 | 14.16 |
| 155 | *Garcinia gummi-gutta* | 0 | 9.23 | 29.5 | 0 | 59.65 | 1.62 | 0 |
| 156 | *Garcinia indica* | 0 | 46.26 | 0 | 0 | 0 | 32.97 | 20.77 |
| 157 | *Garcinia morella* | 17.14 | 6.03 | 17.83 | 0 | 58.67 | 0 | 0.33 |
| 158 | *Garcinia rubroechinata* | 0 | 0 | 0 | 0 | 0 | 100 | 0 |
| 159 | *Garcinia talbotii* | 0.97 | 10.18 | 32.83 | 10.37 | 41.26 | 0.02 | 4.37 |
| 160 | *Glochidion ellipticum* | 0 | 58.15 | 0 | 0 | 34.53 | 0 | 7.31 |
| 161 | *Gluta travancorica* | 0 | 3.39 | 0.13 | 0 | 0 | 96.48 | 0 |
| 162 | *Glycosmis macrocarpa* | 0 | 11.09 | 43.14 | 4.91 | 9.84 | 31.01 | 0 |
| 163 | *Glycosmis pentaphylla* | 0 | 53.4 | 26.05 | 0 | 18.67 | 1.59 | 0.3 |
| 164 | *Gomphandra coriacea* | 2.44 | 0.22 | 0.42 | 3.29 | 36.33 | 57.29 | 0 |
| 165 | *Gomphandra tetrandra* | 0 | 3.07 | 30.41 | 9.1 | 39.18 | 0.16 | 18.09 |
| 166 | *Goniothalamus cardiopetalus* | 0 | 63.1 | 34.25 | 0 | 0 | 1.06 | 1.58 |
| 167 | *Goniothalamus rhynchantherus* | 0 | 0 | 0 | 0 | 0 | 100 | 0 |
| 168 | *Goniothalamus wightii* | 0 | 0 | 0 | 0 | 0 | 100 | 0 |

| | | | | | | | |
|---|---|---|---|---|---|---|---|
| 169 | *Gordonia obtusa* | 0.62 | 40.13 | 0 | 0 | 9.28 | 46.5 | 3.47 |
| 170 | *Gymnacranthera canarica* | 0 | 0 | 100 | 0 | 0 | 0 | 0 |
| 171 | *Helicia nilagirica* | 0 | 0 | 0 | 0 | 100 | 0 | 0 |
| 172 | *Heritiera papilio* | 8.78 | 0.83 | 3.64 | 0 | 80.87 | 3.05 | 2.83 |
| 173 | *Heynea trijuga* | 0 | 15.3 | 59.52 | 0 | 5.76 | 19.12 | 0.3 |
| 174 | *Holigarna arnottiana* | 12.99 | 15.08 | 12.41 | 4.4 | 31.67 | 14.34 | 9.11 |
| 175 | *Holigarna ferruginea* | 0 | 49.84 | 41.79 | 0 | 0 | 3.51 | 4.86 |
| 176 | *Holigarna grahamii* | 0 | 10.03 | 31.58 | 0 | 57.59 | 0.06 | 0.74 |
| 177 | *Holigarna nigra* | 0 | 5.44 | 8.28 | 0 | 83.45 | 2.83 | 0 |
| 178 | *Homalium ceylanicum* | 0 | 31.71 | 27.29 | 0 | 35.08 | 2.93 | 3 |
| 179 | *Hopea canarensis* | 0 | 13.86 | 0 | 0 | 19.28 | 48.83 | 18.03 |
| 180 | *Hopea erosa* | 0 | 0 | 0 | 0 | 0 | 100 | 0 |
| 181 | *Hopea glabra* | 0 | 0 | 0 | 0 | 0 | 100 | 0 |
| 182 | *Hopea parviflora* | 13.93 | 21.61 | 11.41 | 1.9 | 2.03 | 39.08 | 10.03 |
| 183 | *Hopea ponga* | 0 | 1.97 | 48.4 | 0 | 47.99 | 0 | 1.64 |
| 184 | *Hopea racophloea* | 0 | 0 | 47.23 | 0 | 47.33 | 3.07 | 2.37 |
| 185 | *Humboldtia brunonis* | 0 | 21.29 | 22.78 | 0 | 0 | 44.1 | 11.83 |
| 186 | *Humboldtia decurrens* | 0 | 0 | 9.87 | 0 | 0 | 81.37 | 8.76 |
| 187 | *Humboldtia vahliana* | 0 | 0 | 100 | 0 | 0 | 0 | 0 |
| 188 | *Hunteria zeylanica* | 0.02 | 1.73 | 17.73 | 1.39 | 5.03 | 63.57 | 10.53 |
| 189 | *Hydnocarpus alpina* | 0 | 0 | 20.61 | 4.17 | 0.25 | 30.7 | 44.27 |
| 190 | *Hydnocarpus macrocarpa* | 0 | 72.87 | 0 | 0 | 0 | 27.13 | 0 |
| 191 | *Hydnocarpus pentandrus* | 0 | 0.82 | 29.71 | 0 | 57.34 | 0 | 12.14 |
| 192 | *Isonandra lanceolata* | 0 | 2.34 | 10.31 | 0 | 71.74 | 12.99 | 2.62 |
| 193 | *Ixora alba* | 0 | 10.5 | 34.25 | 0 | 0 | 28.68 | 26.57 |
| 194 | *Ixora brachiata* | 0 | 0.42 | 38.52 | 1.77 | 50.53 | 0 | 8.76 |
| 195 | *Ixora elongata* | 0 | 43.44 | 7.2 | 7.11 | 0 | 42.24 | 0 |
| 196 | *Ixora lanceolaria* | 0 | 0 | 0 | 0 | 100 | 0 | 0 |
| 197 | *Ixora nigricans* | 0 | 11.73 | 8.1 | 0 | 80.17 | 0 | 0 |

| | | | | | | | | |
|---|---|---|---|---|---|---|---|---|
| 198 | *Kingiodendron pinnatum* | 0 | 0 | 69.87 | 0.09 | 19.45 | 0.66 | 9.93 |
| 199 | *Knema attenuata* | 0 | 10.65 | 30.83 | 0 | 55.01 | 2.63 | 0.88 |
| 200 | *Lagerstroemia microcarpa* | 0 | 3.82 | 25.92 | 0 | 49.85 | 11.65 | 8.76 |
| 201 | *Lasianthus acuminatus* | 0 | 46.28 | 20.65 | 0 | 15.21 | 0.02 | 17.85 |
| 202 | *Lasianthus jackianus* | 0 | 36.05 | 40.2 | 0 | 0 | 23.75 | 0 |
| 203 | *Lasianthus rostratus* | 0 | 0 | 0 | 0 | 8.46 | 88.58 | 2.96 |
| 204 | *Leea indica* | 0 | 44.54 | 0 | 0 | 55.46 | 0 | 0 |
| 205 | *Lepisanthes erecta* | 0 | 4.1 | 0 | 0 | 0 | 95.9 | 0 |
| 206 | *Lepisanthes tetraphylla* | 0 | 50.18 | 0 | 0 | 30.53 | 19.29 | 0 |
| 207 | *Leptonychia caudata* | 0 | 0 | 13.79 | 0 | 0 | 86.21 | 0 |
| 208 | *Ligustrum perrottetii* | 0 | 0 | 33.2 | 0.4 | 4.67 | 0 | 61.72 |
| 209 | *Litsea floribunda* | 19.4 | 9.65 | 5 | 0 | 65.29 | 0.13 | 0.53 |
| 210 | *Litsea ghatica* | 19.4 | 35.59 | 0 | 0 | 5.82 | 35.68 | 3.51 |
| 211 | *Litsea glabrata* | 0 | 7.4 | 0 | 0 | 84.22 | 8.38 | 0 |
| 212 | *Litsea keralana* | 0 | 58.26 | 0 | 0 | 9.56 | 32.18 | 0 |
| 213 | *Litsea laevigata* | 0 | 55.03 | 0 | 0 | 43.94 | 0 | 1.03 |
| 214 | *Litsea mysorensis* | 0 | 50.25 | 4.39 | 1.35 | 31.43 | 6.14 | 6.45 |
| 215 | *Litsea oleoides* | 0 | 6.93 | 7.8 | 0 | 81.74 | 1.73 | 1.8 |
| 216 | *Litsea stocksii* | 0 | 27.21 | 72.79 | 0 | 0 | 0 | 0 |
| 217 | *Litsea travancorica* | 0 | 0 | 0.87 | 0 | 0 | 97.99 | 1.14 |
| 218 | *Litsea wightiana* | 0 | 0 | 14.66 | 0 | 0 | 85.34 | 0 |
| 219 | *Lophopetalum wightianum* | 32.7 | 9.78 | 24.98 | 1.55 | 18.99 | 6.8 | 5.2 |
| 220 | *Macaranga indica* | 0 | 0 | 0 | 0 | 56.98 | 43.02 | 0 |
| 221 | *Macaranga peltata* | 0 | 15.84 | 22.88 | 0 | 49.09 | 6.69 | 5.5 |
| 222 | *Madhuca bourdillonii* | 0 | 41.22 | 8.87 | 0 | 0 | 49.91 | 0 |
| 223 | *Madhuca neriifolia* | 0 | 0 | 28.54 | 0 | 28.09 | 43.37 | 0 |
| 224 | *Maesa indica* | 0 | 29.41 | 0 | 0 | 69.85 | 0 | 0.74 |
| 225 | *Magnolia nilagirica* | 0 | 0 | 0 | 0 | 100 | 0 | 0 |
| 226 | *Mallotus aureopunctatus* | 0 | 67.47 | 0 | 0 | 25.55 | 6.98 | 0 |

| | | | | | | | | |
|---|---|---|---|---|---|---|---|---|
| 227 | *Mallotus beddomei* | 0 | 0 | 11.84 | 0 | 0 | 88.16 | 0 |
| 228 | *Mallotus nudiflorus* | 0 | 0 | 31.97 | 0 | 66.52 | 1.51 | 0 |
| 229 | *Mallotus philippensis* | 6.35 | 12.59 | 20.32 | 9.28 | 41.96 | 7.78 | 1.73 |
| 230 | *Mallotus resinosus* | 0 | 10.51 | 59.44 | 5.2 | 0 | 19.44 | 5.41 |
| 231 | *Mallotus rhamnifolius* | 0 | 0 | 24.56 | 0 | 0 | 75.44 | 0 |
| 232 | *Mammea suriga* | 0 | 0 | 0 | 0 | 0 | 100 | 0 |
| 233 | *Mangifera indica* | 0 | 1.52 | 23.36 | 0 | 60.11 | 11.45 | 3.56 |
| 234 | *Margaritaria indica* | 0 | 26.52 | 73.48 | 0 | 0 | 0 | 0 |
| 235 | *Mastixia arborea* | 0 | 16.32 | 18.63 | 0 | 62.38 | 1.21 | 1.46 |
| 236 | *Maytenus rothiana* | 0 | 0 | 0 | 0 | 0 | 100 | 0 |
| 237 | *Meiogyne pannosa* | 0 | 13.74 | 16.92 | 3.22 | 64.33 | 1.79 | 0 |
| 238 | *Meiogyne ramarowii* | 0 | 1.85 | 29.45 | 0 | 0 | 24.17 | 44.53 |
| 239 | *Melicope lunu-ankenda* | 14.69 | 1.74 | 15.15 | 0.8 | 63.31 | 4.31 | 0 |
| 240 | *Meliosma pinnata* | 0 | 3.11 | 54.07 | 0 | 0 | 42.82 | 0 |
| 241 | *Memecylon gracile* | 0 | 0 | 14.73 | 0 | 71.65 | 1.69 | 11.94 |
| 242 | *Memecylon heyneanum* | 0 | 11.93 | 12.84 | 0 | 0 | 75.11 | 0.12 |
| 243 | *Memecylon malabaricum* | 0 | 52.27 | 0 | 0 | 0 | 46.63 | 1.1 |
| 244 | *Memecylon pseudogracile* | 0 | 23.13 | 0.12 | 59.91 | 0 | 9.84 | 6.99 |
| 245 | *Memecylon randerianum* | 0 | 0 | 22.06 | 0 | 31.62 | 17.67 | 28.65 |
| 246 | *Memecylon subsessile* | 24.88 | 0 | 45 | 0 | 0 | 29.83 | 0.29 |
| 247 | *Memecylon talbotianum* | 0 | 7 | 25.45 | 0 | 67.55 | 0 | 0 |
| 248 | *Memecylon terminale* | 0 | 49.19 | 24.97 | 0 | 0 | 25.78 | 0.07 |
| 249 | *Memecylon umbellatum* | 0 | 67.77 | 0 | 0 | 12.1 | 0 | 20.13 |
| 250 | *Memecylon wightii* | 7.85 | 17.8 | 25 | 0.07 | 43.24 | 6.04 | 0 |
| 251 | *Mesua ferrea* | 0 | 5.61 | 9.09 | 0 | 76.58 | 0.24 | 8.48 |
| 252 | *Meteoromyrtus wynaadensis* | 0 | 12.71 | 0 | 0 | 74.19 | 0 | 13.1 |
| 253 | *Meyna laxiflora* | 18.4 | 0 | 0 | 0 | 0 | 53.57 | 28.03 |
| 254 | *Microtropis latifolia* | 0 | 10.63 | 8.1 | 0 | 79.74 | 0 | 1.52 |
| 255 | *Miliusa gokhalaei* | 0 | 5.58 | 1.08 | 0 | 7.88 | 85.46 | 0 |

| | | | | | | | | |
|---|---|---|---|---|---|---|---|---|
| 256 | *Miliusa nilagirica* | 0 | 0 | 10.23 | 4.82 | 84.05 | 0 | 0.89 |
| 257 | *Mimusops elengi* | 0 | 0.31 | 38.95 | 0 | 39.12 | 4.05 | 17.57 |
| 258 | *Mitragyna parvifolia* | 0 | 0 | 18.7 | 0 | 8.57 | 47.63 | 25.1 |
| 259 | *Mitragyna tubulosa* | 0 | 0.35 | 50.65 | 0 | 37.13 | 0 | 11.88 |
| 260 | *Mitrephora grandiflora* | 0 | 0 | 0 | 0 | 0 | 79.9 | 20.1 |
| 261 | *Murraya koenigii* | 0 | 10.21 | 37.3 | 0 | 0 | 52.49 | 0 |
| 262 | *Murraya paniculata* | 0 | 0 | 33.74 | 0 | 0 | 66.26 | 0 |
| 263 | *Myristica beddomei* | 0 | 8.15 | 20.42 | 0 | 66.89 | 4.53 | 0 |
| 264 | *Myristica malabarica* | 0 | 30.27 | 15.76 | 0 | 25.14 | 28.83 | 0 |
| 265 | *Neolitsea fischeri* | 0 | 5.36 | 0 | 0 | 3.1 | 91.54 | 0 |
| 266 | *Neolitsea zeylanica* | 0 | 48.62 | 0 | 0 | 1.35 | 17.65 | 32.38 |
| 267 | *Nothapodytes nimmoniana* | 0 | 43.26 | 0 | 0 | 39.58 | 0 | 17.16 |
| 268 | *Nothopegia aureofulva* | 0 | 0 | 11.77 | 0 | 0 | 88.23 | 0 |
| 269 | *Nothopegia beddomei* | 0 | 14.33 | 24.33 | 0 | 56.28 | 2.42 | 2.63 |
| 270 | *Nothopegia racemosa* | 0 | 44.04 | 0 | 0 | 50.39 | 3.87 | 1.7 |
| 271 | *Nothopegia travancorica* | 0 | 1.44 | 14.61 | 0 | 53.14 | 5.58 | 25.22 |
| 272 | *Ocotea lancifolia* | 0 | 10.8 | 0 | 0 | 59.52 | 28.94 | 0.74 |
| 273 | *Octotropis travancorica* | 0 | 3.49 | 3.33 | 0 | 0 | 93.17 | 0.01 |
| 274 | *Olea dioica* | 0 | 21.1 | 22.95 | 3.36 | 46.78 | 4.61 | 1.21 |
| 275 | *Oreocnide integrifolia* | 0 | 7.69 | 18.9 | 0 | 23.24 | 50.17 | 0 |
| 276 | *Ormosia travancorica* | 0 | 0.76 | 0 | 0 | 22.97 | 76.28 | 0 |
| 277 | *Orophea erythrocarpa* | 0 | 0.51 | 29.77 | 0 | 68.15 | 0 | 1.57 |
| 278 | *Orophea sivarajanii* | 0 | 0 | 5.09 | 0 | 70.65 | 20.62 | 3.64 |
| 279 | *Orophea thomsonii* | 0 | 0 | 33.42 | 0 | 0 | 66.58 | 0 |
| 280 | *Otonephelium stipulaceum* | 0 | 0 | 24.69 | 3.31 | 69.05 | 0.12 | 2.82 |
| 281 | *Pajanelia longifolia* | 0 | 6.6 | 30.86 | 0 | 52.14 | 9.67 | 0.73 |
| 282 | *Palaquium bourdillonii* | 0 | 4.16 | 0 | 0 | 0 | 95.84 | 0 |
| 283 | *Palaquium ellipticum* | 0 | 6.85 | 13.42 | 0 | 70.92 | 3.13 | 5.69 |
| 284 | *Pandanus furcatus* | 0 | 9.97 | 24.9 | 0 | 54.64 | 7.59 | 2.89 |

| | | | | | | | | |
|---|---|---|---|---|---|---|---|---|
| 285 | *Paracroton pendulus subsp. zeylanicus* | 0 | 3.32 | 20.05 | 0 | 51.06 | 1.08 | 24.49 |
| 286 | *Pavetta sp* | 0 | 0 | 100 | 0 | 0 | 0 | 0 |
| 287 | *Persea macrantha* | 1.48 | 10.65 | 16.64 | 0.05 | 67.53 | 2.95 | 0.71 |
| 288 | *Pinanga dicksonii* | 0 | 14.87 | 23.04 | 0 | 59.77 | 2.32 | 0 |
| 289 | *Poeciloneuron indicum* | 0 | 61.79 | 8.98 | 0 | 2.2 | 10.49 | 16.55 |
| 290 | *Polyalthia coffeoides* | 0 | 0 | 13.03 | 0 | 63.29 | 1.76 | 21.93 |
| 291 | *Polyalthia fragrans* | 10.05 | 5.57 | 32.62 | 0 | 39.55 | 3.95 | 8.25 |
| 292 | *Polyalthia malabarica* | 0.16 | 4.26 | 6.08 | 4.11 | 0 | 28.79 | 56.6 |
| 293 | *Prismatomeris tetrandra* | 0 | 0 | 0 | 0 | 0 | 100 | 0 |
| 294 | *Prunus ceylanica* | 0 | 19.68 | 8.23 | 0 | 10.19 | 61.59 | 0.32 |
| 295 | *Psychotria anamallayana* | 0 | 0 | 10.98 | 0 | 55.97 | 33.05 | 0 |
| 296 | *Psychotria dalzellii* | 0.17 | 62.59 | 0 | 0 | 0 | 24.01 | 13.22 |
| 297 | *Psychotria flavida* | 0 | 49.34 | 40.22 | 0 | 0 | 0 | 10.44 |
| 298 | *Psychotria nigra* | 6.38 | 8.04 | 9.65 | 0 | 73.92 | 1.85 | 0.16 |
| 299 | *Psychotria truncata* | 0 | 20.65 | 11.19 | 8.57 | 21.16 | 8.42 | 30 |
| 300 | *Psydrax dicoccos* | 0.29 | 57.38 | 29.17 | 0 | 12.63 | 0.53 | 0 |
| 301 | *Pterospermum diversifolium* | 0 | 44.6 | 40.08 | 0 | 0 | 15.12 | 0.2 |
| 302 | *Pterospermum reticulatum* | 0 | 0.59 | 24.81 | 0 | 54.5 | 16.76 | 3.34 |
| 303 | *Pterospermum rubiginosum* | 0 | 0 | 0 | 0 | 0 | 0 | 100 |
| 304 | *Pterygota alata* | 0 | 3.55 | 36.51 | 10.12 | 0 | 11.59 | 38.24 |
| 305 | *Rapanea wightiana* | 0 | 39.65 | 0 | 0 | 50.21 | 6.75 | 3.4 |
| 306 | *Reinwardtiodendron anamalaiense* | 0 | 16.74 | 16.16 | 0 | 44.37 | 19.16 | 3.57 |
| 307 | *Sageraea laurina* | 25.16 | 2.86 | 55.81 | 0 | 16.15 | 0 | 0.02 |
| 308 | *Sageraea thwaitesii* | 0 | 10.4 | 12.98 | 1.54 | 0 | 75.08 | 0 |
| 309 | *Saprosma corymbosum* | 0 | 2.47 | 4.37 | 0 | 0 | 93.16 | 0 |
| 310 | *Saprosma glomerata* | 0 | 4.49 | 2.87 | 0 | 89.75 | 0 | 2.88 |
| 311 | *Schefflera micrantha* | 0 | 6.96 | 1.46 | 0 | 84.62 | 0.04 | 6.93 |
| 312 | *Scolopia crenata* | 0 | 18.18 | 0 | 0 | 57.39 | 24.42 | 0 |

| | | | | | | | | |
|---|---|---|---|---|---|---|---|---|
| 313 | *Semecarpus auriculata* | 0 | 0 | 41.84 | 0 | 32.28 | 24.48 | 1.41 |
| 314 | *Semecarpus travancorica* | 0 | 0 | 15.03 | 0 | 0 | 84.97 | 0 |
| 315 | *Sterculia guttata* | 0 | 5.7 | 12.09 | 0 | 69.17 | 7.94 | 5.1 |
| 316 | *Stereospermum tetragonum* | 0 | 18.81 | 0 | 0 | 26.49 | 35.16 | 19.54 |
| 317 | *Strombosia ceylanica* | 0 | 21.88 | 20.01 | 0 | 0 | 19.33 | 38.78 |
| 318 | *Symplocos cochinchinensis* | 0 | 0 | 100 | 0 | 0 | 0 | 0 |
| 319 | *Symplocos macrophylla* | 0 | 75.35 | 20.78 | 0 | 0 | 3.41 | 0.45 |
| 320 | *Symplocos racemosa* | 13.39 | 6.34 | 13.33 | 6.39 | 48.66 | 11.72 | 0.17 |
| 321 | *Symplocos rosea* | 0 | 7.53 | 16.36 | 1.81 | 0 | 74.3 | 0 |
| 322 | *Syzygium benthamianum* | 0 | 0.13 | 0 | 0 | 0 | 99.87 | 0 |
| 323 | *Syzygium caryophyllatum* | 0 | 0.76 | 0 | 0 | 8.33 | 62.03 | 28.88 |
| 324 | *Syzygium cumini* | 0 | 2.94 | 20.56 | 0 | 68.18 | 7.17 | 1.16 |
| 325 | *Syzygium densiflorum* | 0 | 0.34 | 0.98 | 0 | 90.92 | 4.39 | 3.37 |
| 326 | *Syzygium gardneri* | 0 | 14.47 | 17.39 | 0 | 51.85 | 7.61 | 8.68 |
| 327 | *Syzygium grande* | 8.84 | 2.99 | 0.98 | 21.62 | 0 | 19.59 | 45.98 |
| 328 | *Syzygium hemisphericum* | 0 | 16.52 | 29.1 | 0 | 48.04 | 6.06 | 0.28 |
| 329 | *Syzygium laetum* | 0 | 17.06 | 22.88 | 0 | 58.54 | 1.16 | 0.36 |
| 330 | *Syzygium lanceolatum* | 0 | 5.92 | 3.46 | 17.64 | 62.75 | 8.15 | 2.08 |
| 331 | *Syzygium mundagam* | 0 | 0 | 7.55 | 0 | 0.35 | 89.3 | 2.8 |
| 332 | *Syzygium munronii* | 0 | 0.2 | 11.31 | 0 | 87.78 | 0 | 0.71 |
| 333 | *Syzygium rubicundum* | 0 | 0 | 0 | 0 | 100 | 0 | 0 |
| 334 | *Tabernaemontana alternifolia* | 0 | 0.45 | 44.47 | 8.7 | 43.73 | 2.63 | 0.02 |
| 335 | *Tabernaemontana gamblei* | 0 | 0.22 | 11.79 | 7.38 | 0 | 74.99 | 5.62 |
| 336 | *Tarenna nilagirica* | 0 | 24.16 | 54.25 | 0 | 0 | 10.56 | 11.04 |
| 337 | *Terminalia bellirica* | 28.71 | 7.03 | 0 | 0 | 0 | 52.08 | 12.18 |
| 338 | *Terminalia paniculata* | 0.26 | 0 | 35.94 | 0 | 46.2 | 0 | 17.59 |
| 339 | *Terminalia travancorensis* | 0 | 2.32 | 33.02 | 0 | 24.29 | 7.52 | 32.85 |
| 340 | *Tetrameles nudiflora* | 0 | 0 | 0 | 0 | 0 | 100 | 0 |
| 341 | *Toona ciliata* | 0 | 46.41 | 53.59 | 0 | 0 | 0 | 0 |

| | | | | | | | | |
|---|---|---|---|---|---|---|---|---|
| 342 | *Turpinia malabarica* | 0 | 31.9 | 0 | 0 | 23.16 | 44.94 | 0 |
| 343 | *Vateria indica* | 0 | 9.48 | 21.97 | 0 | 18.27 | 46.54 | 3.74 |
| 344 | *Vepris bilocularis* | 0 | 18.84 | 13.21 | 0 | 46.98 | 17.77 | 3.2 |
| 345 | *Vitex altissima* | 0 | 15.94 | 0.82 | 0 | 18.75 | 45.72 | 18.76 |
| 346 | *Walsura trifoliolata* | 0 | 0 | 59.71 | 0 | 40.29 | 0 | 0 |
| 347 | *Xanthophyllum arnottianum* | 42.19 | 0.11 | 16.74 | 4.05 | 21 | 15.45 | 0.45 |
| 348 | *Xantolis tomentosa* | 0 | 0 | 15.9 | 0 | 30.24 | 51.93 | 1.93 |
